# PrediTALE: A novel model learned from quantitative data allows for new perspectives on TALE targeting

**DOI:** 10.1101/522458

**Authors:** Annett Erkes, Stefanie Mücke, Maik Reschke, Jens Boch, Jan Grau

## Abstract

Plant-pathogenic *Xanthomonas* bacteria secret transcription activator-like effectors (TALEs) into host cells, where they act as transcriptional activators on plant target genes to support bacterial virulence. TALEs have a unique modular DNA-binding domain composed of tandem repeats. Two amino acids within each tandem repeat, termed repeat-variable diresidues, bind to contiguous nucleotides on the DNA sequence and determine target specificity.

In this paper, we propose a novel approach for TALE target prediction to identify potential virulence targets. Our approach accounts for recent findings concerning TALE targeting, including frame-shift binding by repeats of aberrant lengths, and the flexible strand orientation of target boxes relative to the transcription start of the downstream target gene. The computational model can account for dependencies between adjacent RVD positions. Model parameters are learned from the wealth of quantitative data that have been generated over the last years.

We benchmark the novel approach, termed PrediTALE, using RNA-seq data after *Xanthomonas* infection in rice, and find an overall improvement of prediction performance compared with previous approaches. Using PrediTALE, we are able to predict several novel putative virulence targets. However, we also observe that no target genes are predicted by any prediction tool for several TALEs, which we term orphan TALEs for this reason. We postulate that one explanation for orphan TALEs are incomplete gene annotations and, hence, propose to replace promoterome-wide by genome-wide scans for target boxes. We demonstrate that known targets from promoterome-wide scans may be recovered by genome-wide scans, whereas the latter, combined with RNA-seq data, are able to detect putative targets independent of existing gene annotations.

**Author summary:** Diseases caused by plant-pathogenic *Xanthomonas* bacteria are a serious threat for many important crop plants including rice. Efficiently protecting plants from these pathogens requires a deeper understanding of infection strategies. For many *Xanthomonas* strains, such infection strategies depend on a special class of effector proteins, termed transcription activator-like effectors (TALEs). TALEs may specifically activate genes of the host plant and, by this means, re-program the plant cell for the benefit of the pathogen. Target sequences and, consequently, target genes of a specific TALE may be predicted computationally from its amino acids. Here, we propose a novel approach for TALE target prediction that makes use of several insights into TALE biology but also of broad experimental data gained over the last years. We demonstrate that this approach yields a higher prediction accuracy than previous approaches. We further postulate that a strategy change from a restricted search only considering promoters of annotated genes to a broad genome-wide search is feasible and yields novel targets including previously neglected protein-coding genes but also non-coding RNAs of possibly regulatory function.

## Introduction

Many crop plants including rice can be infected by *Xanthomonas* bacteria causing disease in the affected plants, which results in substantial yield losses. Many strains of *Xanthomonas oryzae* pv. *oryzae* (*Xoo*) and *Xanthomonas oryzae* pv. *oryzicola* (*Xoc*) express a specific type of effector protein called transcription activator-like effectors (TALEs). TALE proteins function as transcription factors in infected host cells [1], and contain a nuclear localization signal, a DNA-binding domain, and an activation domain. The DNA-binding domain consists of tandem repeats that bind to the promoter of plant target genes. Each repeat consists of approximately 34 highly conserved amino acids (AAs), except for the amino acids at position 12 and 13, which are termed repeat variable diresdue (RVD) and are responsible for DNA specificity. The repeat domain forms right-handed superhelical structure, while the RVD is situated within a loop accessing the DNA [2, 3]. Each RVD binds to one nucleotide of the target box [4, 5], where amino acid 13 binds to the sense strand and amino acid 12 stabilizes the repeat structure. Hence, the specificity of each TALE is determined by its RVD sequence. In addition, most known target boxes are directly preceeded by a ‘T’, while ‘C’ and ‘A’ occur with decreasing frequencies, which is also referred to as “position 0” of the target box.

Some repeats deviate from the common length of 34 AAs and have, for this reason, been termed *aberrant* repeats. Aberrant repeats may loop out of the repeat array when a TALE binds to its DNA target box and by this means allow for increased flexibility, also binding to frame-shifted target boxes [6].

Different *Xoo* and *Xoc* strains express different repertoires of TALEs, where a single strain may host up to 27 TALEs [7–10].

Naturally occurring TALEs may activate susceptibility (S) genes that are responsible for bacterial growth, proliferation and disease development, but also disease resistance (R) genes [1].

The names of TALEs and TALE classes are based on the nomenclature introduced by the tool AnnoTALE [11]. TALEs are clustered according to the similarity of their RVD sequence and divided into classes.

Target boxes upstream of all known major virulence targets are located in forward orientation relative to the transcription start site (TSS). Recently, target boxes of TALEs have been reported to be also functional in reverse orientation relative to the transcription start site (TSS) of their target gene [12, 13]. However, reverse binding seems to be rather an exception than a general rule [13]. Accurate predictions of target boxes of TALEs are important for studying naturally occurring TALEs and determining their virulence targets, but also for the identification of target and off-target sequences of artificially designed TALEs. Over the last years, several tools have been designed for the *in-silico* prediction of TALE target boxes based on the RVD sequence of a given TALE and, subsequently, for the identification of target genes.

The TALE-NT suite includes “Target Finder”, a tool for predicting target boxes of TALEs based on their RVD sequence. It is available as online or command line application (http://tale-nt.cac.cornell.edu/) [14, 15]. In Target Finder, predictions are based on a position weight matrix calculated from frequencies of naturally occurring RVD-nucleotide associations. The user can choose whether the target box should start with nucleotide T or C.

Talvez is another prediction tool that uses PWMs to model RVD-nucleotide interactions [16]. It differs from Target Finder in deriving specificities of rare RVDs from those of common RVDs with the same 13th amino acid. Target sequences may only begin with nucleotide T or C, with a lower score assigned in the case of cytosine. In addition, Talvez may explicitly model that mismatches are tolerated to a larger degree if these are located near the C terminus [17]. Users of Talvez can choose between web-based and command line applications.

TALgetter [18] uses a local mixture model to predict TAL target sequences. The specificities were learned from 267 pairs of TALEs and target sites with qualitative information whether the pair is functional or not. According to Streubel *et al.* [19], the efficiencies of different RVDs are non-identical. The TALgetter model adapts a similar concept using an importance term, which is learned independently from the specificity of each RVD. TALgetter is implemented within the Java framework Jstacs [20], and is available as online and command line program.

In the web tool SIFTED [21], specificity data from a large-scale study using protein-binding microarrays (PBMs) were used for training model parameters. For this purpose, 21 TALEs constructed exclusively from the most common four RVDs (NI, HD, NN, NG) were designed and their binding specificity measured on ≈ 5,000-20,000 DNA sequences per protein using PBMs. However, we will not consider SIFTED in the remainder of this manuscripts, as the SIFTED web server is currently unavailable and the limited set of RVDs included into SIFTED does not cover the entire spectrum of those occurring in natural TALEs.

Predictions of all of these approaches still comprise a substantial number of false positive predictions, whereas some of the known target genes cannot be detected by these approaches, yet. During the last years, several quantitative studies of TALE binding and transcriptional activation have been published. The studies included quantitative analyses of target gene activation by TALEs spanning naturally occurring RVDs [19, 22], specificities at position 0 of target boxes [23], complete exploration of all possible combinations of amino acids at RVD positions [24, 25], and systematic analyses of those RVDs frequently used in designer TALEs [21].

In this paper, we aim at developing a novel approach for modelling TALE target specificities based on these quantitative data. This approach, called PrediTALE, explicitly captures putative dependencies between adjacent RVDs, dependencies between the first RVD and position 0 of the target box, and also includes positional effects of mismatch tolerance. In contrast to previous approaches, model parameters are adapted by minimizing the difference between prediction scores and quantitative measurements for pairs of TALEs and target boxes. Like previous approaches, PrediTALE also predicts target boxes in reverse strand orientation relative to the TSS, but applies a small penalty term in this case, following the assumption that functional reverse target boxes are rather rare *in planta*. PrediTALE is the first approach to account for aberrant repeats when predicting TALE targets.

## Materials and methods

### Training data

Pairs of TALEs and putative target boxes were collected from systematic, quantitative experiments reported in [19, 22–25]. Data were further processed as detailed in Supplementary Text S1. Basically, data were grouped by TALE, and the global weight was computed as the maximum assay value for the current TALE divided by the maximum assay value reported for all TALEs with the same 13th AA at any position in the current assay. Target values were computed as the assay value of the current pair of TALE and target box divided by maximum assay value over all tested target boxes for the current TALE.

While the normalization of target values has a mostly technical background as it simplified the selection of initial values during numerical optimization of our model (see below), the definition of global weights influences the optimization result. The choice of global weights has been motivated by the observation that some TALE architectures (e.g., those with long successions of identical RVDs, or 12th AAs not occurring in nature) show a generally lower activity than others, which also affects the influence of measurement noise and, hence, the reliability of assay values. With the choice of global weights proposed here, the influence of such TALEs on the final optimization result is reduced, while such TALEs do not need to be completely removed from the training set.

As detailed in Supplementary Text S1, PBM experiments from [21] were filtered for apparent data quality, normalized log-intensities were used as target values, and global weights were defined uniformly for all putative target boxes from a common PBM experiment.

### Bacterial growth conditions

*Xanthomonas oryzae* pv. *oryzae* (*Xoo*) strains PXO83, PXO142 and ICMP 3125^T^ were cultivated in PSA medium at 28°C.

### Plant growth conditions & inoculation

*Oryza sativa* ssp. *japonica* cv. Nipponbare was grown under glasshouse conditions at 28°C (day) and 25°C (night) at 70% relative humidity (RH). Leaves of 4-week-old plants were infiltrated with a needleless syringe and a bacterial suspension with an OD600 of 0.5 in 10 mM MgCl2 as previously described [26].

### RNA-seq data

Rice cultivar Nipponbare leaves were inoculated with *Xoo* strains PXO83, PXO142, ICMP 3125^T^, or MgCl2 as mock control in five spots in an area of approx. 5 cm using a needleless syringe. Two leaves of three rice plants each were inoculated for each strain and control, respectively. 24h later, samples were taken, frozen in liquid nitrogen, and RNA prepared. Three replicates of this experiment were done on separate days and subjected to RNAseq analysis, separately.

RNA-seq data 48h after inoculation with different *Xoc* strains (BLS256, BLS279, CFBP2286, B8-12, L8, RS105, BXOR1, CFBP7331, CFBP7341, CFBP7342), and mock controls [9] were downloaded from Gene Expression Omnibus available under accession number GSE67588.

RNA-seq data were adapter clipped using cutadapt (v1.15) [27] and quality trimmed using trimmomatic (v0.33) [28] with parameters “SLIDINGWINDOW:4:28 MINLEN:50”. Transcript abundances were computed by kallisto [29] using parameters “–single -b 10 -l 200 -s 40” and the cDNA sequences available from http://rice.plantbiology.msu.edu/pub/data/Eukaryotic_Projects/o_sativa/annotation_dbs/pseudomolecules/version_7.0/all.dir/all.cdna. Differentially expressed genes relative to the respective control samples were determined by the R-package sleuth [30].

For the *Xoo* strains and the respective mock control, replicates have been paired during library preparation and sequencing. Hence, the replicate was considered as an additional factor when computing p-values of differential expression for the *Xoo* samples but not for the *Xoc* samples. Differential expression was aggregated on the level of genes using the parameter target mapping of the sleuth function sleuth_prep(), and b-value, p-value, and Benjamini–Hochberg-corrected q-value were recorded. The b-value reported by sleuth when applying a Wald test is actually a biased estimator of the log-fold change. However, as this is a more commonly understood term, we refer to the b-value as “log-fold change” in the remainder of this manuscript. Gene abundances, and sleuth outputs with regard to differential expression are provided as Supplementary Tables T and U, respectively. RNA-seq reads were also mapped to the rice genome (MSU7) to obtain detailed information about transcript coverage. To this end, adapter clipped and quality trimmed reads were mapped using TopHat2 v2.1.0 [31], and the resulting BAM output files were processed in further analyes described below.

### Model

Let ***r*** = *r*_1_ *r*_2_ … *r*_*L*_ denote the RVD sequence of length *L* of a TALE, where *r*_*ℓ*_ ∈ {*AA*, …, *YY*, *A*∗, …, *Y*∗} denotes a single RVD, and *r*_*ℓ*,12_ and *r*_*ℓ*,13_ denote the 12th and 13th AA of that RVD, respectively. Let ***x*** = *x*_0_*x*_1_ … *x*_*L*_ denote a putative target box of length *L* + 1 of that TALE, where *x*_*ℓ*_ ∈ {*A, C, G, T*} and *x*_0_ denotes the nucleotide bound by the zero-th, cryptic repeat.

The general idea of the model proposed here is to model the total binding score of a putative target box ***x*** given the RVD sequence ***r*** of a TALE as a sum of contributions of i) binding to the zero-th repeat, ii) binding to the first RVD, and iii) binding to the remaining RVDs, where the latter two terms may be weighted by an additional, position-dependent but sequence-independent term.

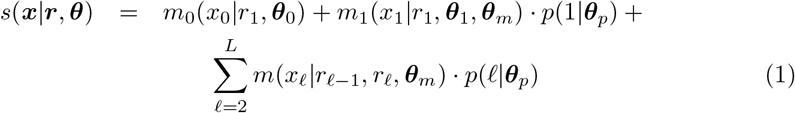

Here, ***θ*** = (***θ***_0_, ***θ***_1_, ***θ***_*m*_, ***θ***_*p*_) denote the sets of real-valued parameters of the term for binding to the zero-th, first, and remaining repeats, and the position-dependent term, respectively.

The term *m*_0_(*x*_0_|*r*_1_, ***θ***_0_) for binding to the zero-th repeat may depend on the first RVD on the TALE, since dependencies between zero-th and first repeat have been observed before [23]. However, our knowledge about such dependencies is limited to the data presently available and, hence, we limit the RVDs for which a dependency is considered to a set 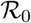. Our data regarding systematic, quantitative analyses of the base preference of the zero-th repeat is limited in general, although it is widely assumed that position 0 in target boxes of natural TALEs is preferentially *T* and less frequently *C*. We include this prior knowledge into *a-priori* parameters 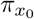.

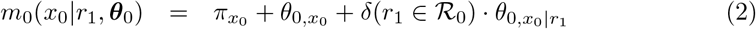

In this paper, we set 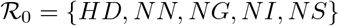 and *π*_*T*_ = log(0.6), *π*_*C*_ = log(0.3), *π*_*A*_ = *π*_*G*_ = log(0.05).

The term *m*_1_(*x*_1_ | *r*_1_, ***θ***_1_, ***θ***_*m*_) for binding to the first repeat depends on the 13th AA *r*_1,13_ of the first RVD *r*_1_, but may be extended by additional terms that either model a general dependency on the complete first RVD (including the 12th AA), and/or a separate base preference for a given 13th AA at the first position. Again, this modularity allows us to adapt the model to the resolution of data available, since a substantial part of RVDs is only covered by the systematic but limited data reported in [24, 25].

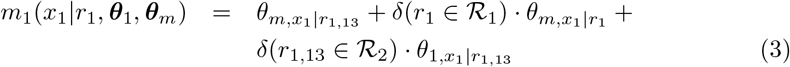

In this paper, we set 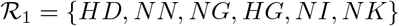 and 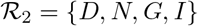.

The term *m*(*x*_*ℓ*_ | *r*_*ℓ*−1_, *r*_*ℓ*_, ***θ***_*m*_) for binding to the remaining repeats again depends on the 13th AA *r*_*ℓ*, 13_ of the current RVD *r_ℓ_*, but may be extened by additional terms that either model a dependency on the complete RVD (with parameters shared with the correponding term used for the first RVD), and/or the complete RVD *r*_*ℓ*_ at the current repeat and the 12th AA *r*_*ℓ*− 1,12_ at the previous repeat:

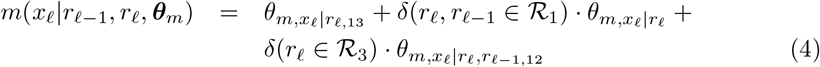

In this paper, we set 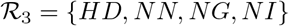.

Finally, we define the position-dependent term as a mixture of two logistic functions and a constant term, where the logistic functions depend on the relative distance of *ℓ* from the start and end of the putative target box, respectively:

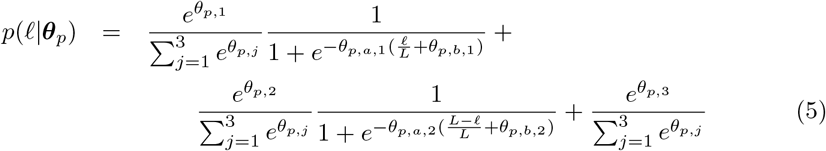

The parameters *θ*_*p,a*,1_ and *θ*_*p,a*,2_ denote the slopes, and *θ*_*p,b*,1_ and *θ*_*p,b*,2_ denote the location parameters of the logistic functions.

### Learning parameters

The training data 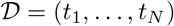 comprise tuples *t*_*i*_ = (***r***_*i*_, ***x***_*i*_, *v*_*i*_, *w*_*i*_, *g*_*i*_) of TALE RVD sequence ***r***_*i*_, target box ***x***_*i*_, target value *v_i_*, global weight *w*_*i*_ and group *g*_*i*_ (cf. sections “Data” and “Model”). Given the current parameter values ***θ***, we may further compute for each pair of TALE and target box, the corresponding model score *s*_*i*_ = *s*(***x*** | ***r***_i_, ***θ***_*i*_). The goal of the learning process is to adapt the parameter values ***θ*** such that the differences between computed scores *s*_*i*_ and target values *v*_*i*_ becomes minimal. However, despite the normalization of target values described in section “Data”, target values from different experimental setups (represented by the groups *g*_*i*_) may live on different scales. Hence, we allow the learning process to linearly transform the computed scores *s*_*i*_ before comparing them to the target values. The total error between target value and prediction score is defined as

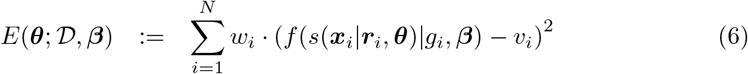

where

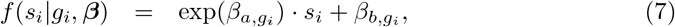

***β*** = (*β*_*a*,1_, *β*_*b*,1_, …, *β*_*a,G*_, *β*_*b,G*_), 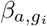 and 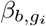 are group-specific scale and shift parameters, respectively, and *G* is the total number of groups in the data set 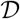.

In addition, we use an *L*_2_ regularization term on the model parameters ***θ*** to avoid overfitting and explosion of parameter values:

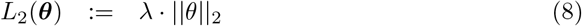

where the regularization parameter *λ* is set to 0.1 in this paper.

The number of model parameters for the different terms varies greatly, depending on the number of conditions (e.g., 12th AA of previous RVD, separate parameters for individual RVDs). This regularization also has the effect that more complex dependency parameters assume values considerably different from 0 only if the modeled specificity cannot be captured by the less complex sets of parameters.

The final objective function is then to minimize sum of the error term 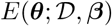 and the regularization term *L*_2_(***θ***) with respect to the parameter values:

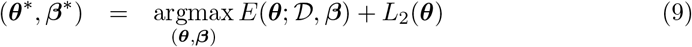

Parameter optimization is performed by a gradient-based quasi-Newton method as implemented in the Jstacs library [20].

The final parameters ***θ***\* of the trained model may then be used to determine prediction scores of previously unseen pairs of TALEs and putative target boxes, whereas the value of ***β***\* is discarded after optimization.

### Prediction of TALE target boxes

For predicting putative TALE target boxes for a given TALE with RVD sequence ***r*** of length *L*, we follow a sliding window approach scanning input sequences ***x***_1_, …, ***x***_*N*_. Input sequences could, for instance, be promoter sequences of annotated genes but also complete chromosomes. Each sub-sequence ***x***_*i,ℓ*_, …, ***x***_*i,ℓ*+*L*_ then serves as input of the model to compute the corresponding score *s*(***x***_*i,ℓ*_, …, ***x***_*i,ℓ*+*L*_ | ***r***, ***θ***\*). To allow for a rough comparison of scores, even between TALEs of different lengths, we normalize this score to the length of the input sequence, i.e., we compute a normalized score as *s*′ (***x***_*i,ℓ*_, …, ***x***_*i,ℓ+L*_ | ***r***, ***θ***\*): = *s*(***x***_*i,ℓ*_, …, ***x***_*i,ℓ+L*_ | ***r***, ***θ***\*)/(*L* + 1)

For scanning promoter sequences, we also provide an option for penalizing predictions of the reverse complementary strand, relative to the orientation of the downstream gene. Specifically, a small constant *c* is subtracted from all prediction scores *s*′ on the reverse complementary strand. Throughout this paper, we use *c* = 0.01.

The scanning process explicitly accounts for aberrant repeats, which may loop out of the repeat array [6]. To this end, we search for putative target boxes with all repeats present in the repeat array, but also all combinations of aberrant repeats removed from the RVD sequence. Due to the normalization of scores by the number of repeats, predictions based on these modified RVD sequences can still be ranked in a common list.

In addition, we provide a box-specific p-value as a statistical measure for the significance of target box predictions. Those p-values may either be computed from a dedicated background set of sequences or from a random sub-sample of the scanned input sequences, where the latter option is used throughout this paper. In either case, scores are computed for the sub-sequences given the current RVD sequences, then a Gaussian distribution is fitted to those score values, and the p-value for a given score is determined from that Gaussian distribution. While the Gaussian distribution does not perfectly fit the true distribution of score values, it allows for computing p-values with high resolution (as opposed to just using percentages of the scores themselves) and even for score values larger than any of the scores in the random sample.

### Genome-wide predictions & filtering

We use PrediTALE for genome-wide prediction in the genome of *Oryza sativa* Nipponbare (MSU7, http://rice.plantbiology.msu.edu/pub/data/Eukaryotic_Projects/o_sativa/annotation_dbs/pseudomolecules/version_7.0/all.dir/all.chrs.con). We make predictions for each TALE of 3 *Xoo* strains and 10 *Xoc* strains. In order to confirm that the predicted target boxes might indeed be bound by the respective TALE, we use the above-mentioned RNA-seq data to determine if there are differentially transcribed regions around a putative target box. For each of the top 100 predictions, we search ± 3000 bp around the predicted site for regions of at least 400 bp that are differentially expressed. Specifically, we count the number of mapped reads for each 400 bp window in replicates of treatment and control. Counts are then normalized relative to the total number of reads within each library, and replicated are averages separately for treatment and control. Here, we consider a region as differentially expressed if the mean normalized number of reads after infection (treatment) is at least 2-fold larger than the mean normalized number of reads in the control experiment. If several, adjacent 400 bp regions meet this criterion, those are joined to a common, longer region.

This procedure is implemented in a tool called DerTALE. As input, DerTALE expects genomic positions, i.e., the position of predicted target boxes, and BAM files of mapped reads for replicates of treatment and control. Region width, thresholds and averaging methods may be adjusted by user parameters.

For each predicted target box, a profile output is generated if there is at least one differential expressed region with a minimum length of 400 bp that does not overlap the target box, or if it overlaps, the differential region starts or ends at most 50 bp upstream or downstream of the target box.

The obtained profiles may be visualized using an auxiliary R script. In addition to the profile data, this R script requires annotations data of already known transcripts in gff3 format. By this means, users may then investigate whether the predicted binding site may activate the transcription of a gene that has not been annotated yet. Here, we use the MSU7 annotation (http://rice.plantbiology.msu.edu/pub/data/Eukaryotic_Projects/o_sativa/annotation_dbs/pseudomolecules/version_7.0/all.dir/all.gff3).

For differentially expressed regions without annotated MSU7 transcript, we searched for similar sequences using blastx of NCBI BLAST+ version 2.7.1 ftp://ftp.ncbi.nlm.nih.gov/blast/executables/blast+/LATEST/ and choose the non-redundant protein sequence (nr) database. In cases, where we did not receive a convincing hit, we additionally compared sequences with blastn against the reference RNA sequences (refseq_rna) database.

### Implementation & scanning speed-up

The model and learning process are implemented using the open-source Jstacs library [20] and will be part of the next Jstacs release.

For scanning large input sequences, e.g., complete genomes of host plant species, an acceptible runtime is essential. Since the parameters at each position of the proposed model depend on the RVD sequence of the TALE of interest but do not include dependencies between different nucleotides of a putative target box, we may convert the model given a fixed TALE RVD sequence into an position weight matrix (PWM) [32, 33]. This allows for a quick computation of prediction scores that may be formulated as the position-wise sum of values stored in the TALE-specific PWM model. We further speed-up the scanning process by pre-computing indexes of overlapping *k*-mers in the same manner as proposed for the TALENoffer application earlier [34].

### Evaluation of prediction results

We compare the performance of the approach presented in this paper to those of established tools for predicting TALE target sites, namely TALESF [14], Talvez [16], and TALgetter [18], based on RNA-seq data after inoculation with different *Xoo* and *Xoc* strains described above.

To this end, we collect the promoter sequences of all transcripts based on the MSU7 assembly and gene models [35] available from http://rice.plantbiology.msu.edu/pub/data/Eukaryotic_Projects/o_sativa/annotation_dbs/pseudomolecules/version_7.0/all.dir/. We consider as promoter the sequence spanning from 300 bp upstream of the transcription start site to 200 bp downstream of the transcription start site or the start codon, whichever comes first, as proposed before [18]. We then run each of the tools using default parameters on the extracted promoter sequence providing the RVD sequences of the TALEs present in the respective *Xanthomonas* strain. Predictions in promoters of different transcripts belonging to the same gene are merged by considering only the prediction yielding the best prediction score.

Assessment of prediction performance based on *in-planta* inoculation experiments with *Xanthomonas* strains harboring multiple TALEs has the inherent complications that i) putative target genes cannot be attributed to one specific TALE based on the RNA-seq data alone and ii) genes showing increased expression after inoculation may either be regulated directly by a TALE binding to their promoter or indirectly via other, regulatory target genes. Hence, we define *true positives* as those genes that have a predicted target box in their promoter and are also up-regulated after inoculation with the respective *Xanthomonas* strain relative to control as derived from RNA-seq data. By contrast, we cannot clearly define *false negatives*, since genes that are up-regulated after inoculation but do not contain a predicted target box in their promoter could be indirect target genes. *False positives*, in turn, would be genes with a predicted target box in their promoter that are *not* up-regulated after *Xanthomonas* inoculation.

A further issue hampering performance assessment by standard methods like receiver operating characteristic (ROC) [36] or precision-recall (PR) curves [37, 38] is that for two of the tools considered (TALESF and Talvez), none of the reported prediction scores is comparable between different TALEs, especially TALEs of different lengths. Hence, we decide to use varying cutoffs on the number of predicted target genes *per TALE* to establish a common ground for comparing all four approaches.

Following these considerations, we collect for each of the four approaches the number of true positive predictions (TPs) for cutoffs on the number of predictions per TALE from 1 (i.e., the top prediction) to 50. We then plot for each approach the number of true positives against this cutoff to obtain a continuous picture of its prediction performance. In addition, we collect for the same cutoffs the number of TALEs with at least one predicted target gene among the true positives.

The area under these curves may serve as a further measure of general prediction performance in analogy to, for instance, the area under the ROC curve.

Finally, we compare the TPs at distinct cutoffs (1, 10, 20, 50) between the four tools. For a specific cutoff, we collect the TPs (or, in analogy, number of TALEs with at least one predicted target) for each of the four tools. Statistical significance of the differences in observed TPs is then assessed by a Quade test [39] using the `quade.test` function in R [40] and pairwise comparisons are performed by the post-hoc test implemented in function `quadeAllPairsTest` of the `PMCMRplus` R-package [41].

### Availability

PrediTALE is available as a web-application based on Galaxy at http://galaxy.informatik.uni-halle.de. Both PrediTALE and DerTALE are available as command line application from http://jstacs.de/index.php/PrediTALE and have also been integrated in AnnoTALE 1.4. Source code is available from https://github.com/Jstacs/Jstacs in packages `projects.tals.linear`, `projects.tals.predictions` and `projects.tals.rnaseq`.

## Results/Discussion

### Benchmarking PrediTALE against previous approaches

In this section, we benchmark the predictions of PrediTALE against those made by one of the previous approaches, namely Target Finder [14], Talvez [16], and TALgetter [18].

To this end, we consider different *Xanthomonas oryzae* pv. *oryzae* (*Xoo*) and *Xanthomonas oryzae* pv. *oryzicola* (*Xoc*) strains for which we have an experimental support of up-regulated genes in *Oryza sativa* after infection based on RNA-seq data. Specifically, we consider the *Xoo* strains ICMP 3125^T^, PXO142 and PXO83 with in-house RNA-seq data available, and the *Xoc* strains B8-12, BLS256, BLS279, BXOR1, CFBP2286, CFBP7331, CFBP7341, CFBP7342, L8 and RS105 based on public RNA-seq data [9]. For the TALEs from the repertoires of these three *Xoo* and ten *Xoc* strains, we determine target gene predictions for each of the previous approaches and for PrediTALE. Predicted target genes are ranked by the corresponding prediction scores of the different approaches per TALE.

First, we study the overlaps between the sets of predicted target genes per approach to investigate how strongly predictions are affected by conceptual differences of these approaches. In Figure 1A, we show Venn diagrams of predicted target genes for the three *Xoo* strains based on the top 20 predictions per TALE, while the corresponding diagrams for the ten *Xoc* strains are available as Supplementary figure S1. In general, we observe a substantial number of unique predictions for each of the four approaches, but especially for Talvez and PrediTALE. By contrast, the overlapping predictions between all four approaches amount to less than a quarter of the total predictions per approach. This demonstrates that prediction results strongly depend on the employed approach. However, prediction accuracy cannot be assessed without an experimental knowledge about genes that are up-regulated *in planta* upon *Xanthomonas* infection. For this reason, we filter predictions based on the corresponding RNA-seq data in the following.

**Fig 1.**
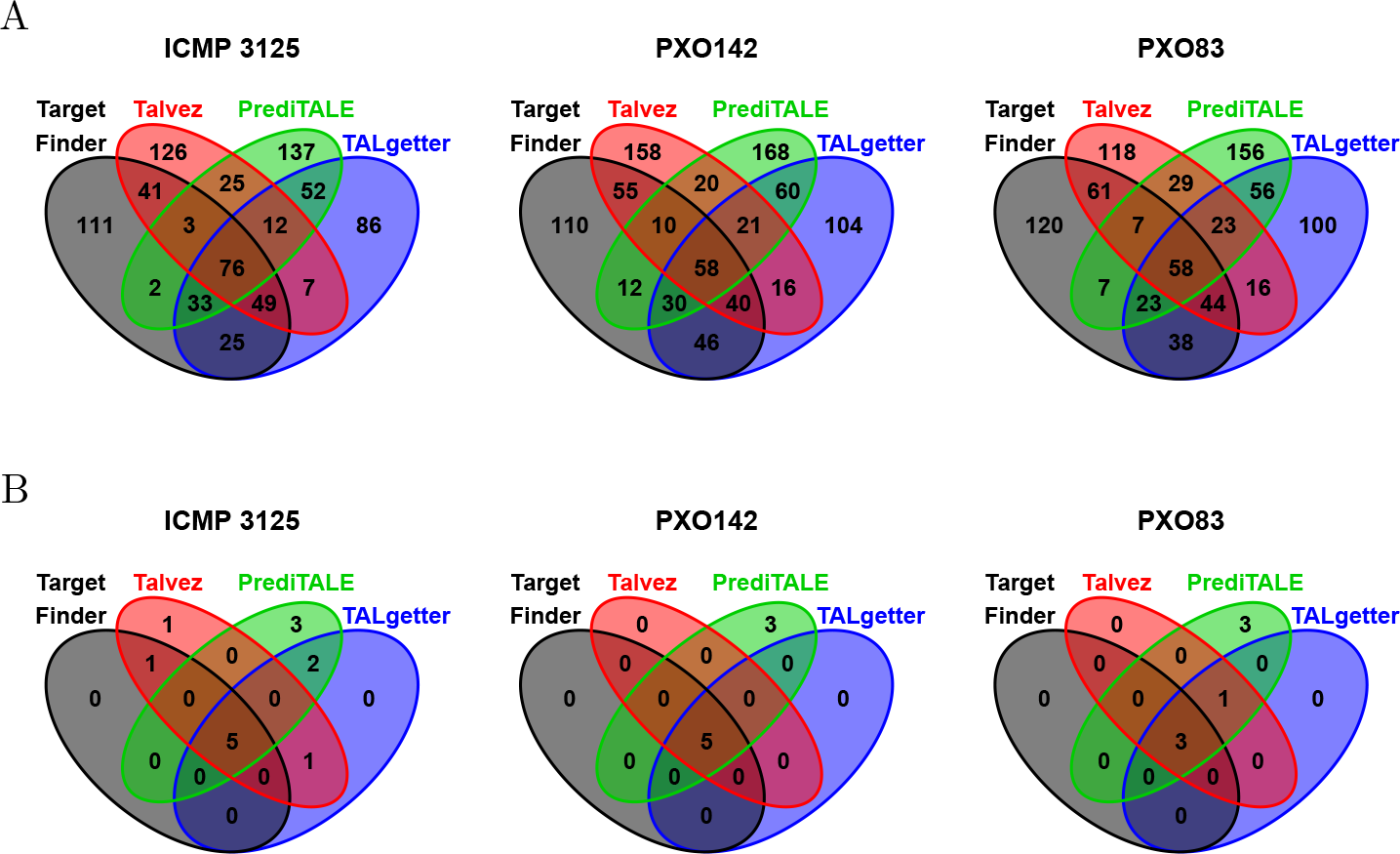
Venn diagrams of predictions of the four approaches considered. (A) For each *Xoo* strain and each approach, we consider the set of target genes obtained as the union of the top 20 predictions per TALE. For *Xoo* ICMP 3125^T^ harboring 17 TALEs, this results in a total number of 340 raw predictions per approach, where the actual number in the diagram may be slightly lower if two TALEs are predicted to target the same gene. For *Xoo* PXO142 (19 TALEs), we obtain 380 raw predictions and for *Xoo* PXO83 (18 TALEs), we obtain 360 raw predictions per approach. (B) Venn diagrams of the subsets of genes from sub-figure A that are also up-regulated according to RNA-seq data.

RNA-seq data for the three *Xoo* strains including previously unpublished data for PXO83, have been collected 24 hours after infection. Collection at this early time point has the advantage that the number of secondary targets, i.e., genes that are up-regulated as a secondary effect of direct TALE targets with regulatory function, should still be low. However, as the infection might not be fully established, yet, the variation between replicates and, hence, the number of significantly differentially expressed genes based on standard FDR-based criteria is rather low (cf. Supplementary table A). As we aim at sensitivity for the benchmark study, i.e., we want to avoid predictions to be erroneously counted as false positives, we consider genes as differentially up-regulated if they obtain an uncorrected p-value below 0.05 and are at least 2-fold up-regulated in this case, which results in 43 (PXO142) to 107 (ICMP 3125^T^) differentially up-regulated genes.

In case of the ten *Xoc* strains, RNA-seq data have been recorded 48 hours after infection. Here, infection should be fully established, but we expect a substantial number of secondary targets to be up-regulated already. Hence, we resort to rather standard thresholds with a FDR-corrected *q − value* < 0.01 and log fold change greater than 2 in this case. Notably, this still results in a larger number of differentially up-regulated genes than for the *Xoo* strains with numbers between 202 (CFBP2286) and 672 (L8).

Given these up-regulated genes as a *ground truth*, we may now count predictions of TALE target boxes in promoters of up-regulated genes as *true positives*, and predictions without observed up-regulation as *false positives*. In Figure 1B, we plot Venn diagrams of the true positives among the top 20 predictions of all four approaches. Notably, we find that the intersection of the predictions of all four approaches constitutes (one of) the largest set(s) in each of the three Venn diagrams. Among the predictions that are unique to one of the four approaches, we consistently find the largest number of true positive predictions for PrediTALE, which indicates the utility of our novel approach. Turning to the ten *Xoc* strains (Supplementary figure S2), we again find the same tendency with regard to the predictions overlapping among all four approaches. However, the number of true positives among the unique predictions shows a less clear picture with a slight advantage towards Talvez, while predictions of PrediTALE often overlap with TALgetter and/or Target Finder. Together, the Venn diagrams for the *Xoo* and *Xoc* strains also illustrate why it is generally beneficial to complement *in silico* TALE target predictions with experimental data about gene regulation.

The results presented so far strongly depend on the thresholds of the ranks of the target predictions but also on the thresholds applied to the RNA-seq data. To address the former problem, we aim at an assessment of target predictions over all rank thresholds, while we will handle the latter by separate evaluations applying different criteria to the RNA-seq data.

As detailed in section “Evaluation of prediction results”, standard performance measures like the area under the ROC curve [36] or the area under the precision-recall curve [37, 38] are inappropriate under this setting. Briefly, we cannot attribute an up-regulated gene to a specific TALE from the TALE repertoire of the strain under study. In addition, genes that are up-regulated in the RNA-seq experiment might also be due to secondary effects of TALE targets, due to general plant response to the bacteria, or due to other classes of effector proteins. Thus, we may not consider up-regulated genes *without* a matching prediction of a TALE target box in their promoter as *false negatives*. Hence, we decide to compare the performance of different approaches by means of the number of *true positive* predictions at different rank cutoffs, i.e., considering the top *N* predicted target genes of each approach.

In Figure 2, we plot the number of true positives for the three *Xoo* strains and each of the four approaches against the total number of predictions per TALE, considering only the highest-ranking prediction up to 50 target predictions per TALE, which we consider a reasonable cutoff under the scenario of manual inspection. In addition, we compute the area under this curve as an overall performance statistic across all rank cutoffs. For ICMP 3125^T^, we find that PrediTALE yields the largest number of rank 1 predictions (4), but also dominates the other three tools for all other rank cutoffs. With regard to the area under the curve (AUC), we find that PrediTALE yields the best overall performance, followed by TALgetter, Talvez, and finally Target Finder. For PXO142, PrediTALE yields a better prediction performance than the other tools starting from rank 4, while Target Finder and TALgetter achieve a larger number of true positive predictions (3) at rank 1. We also observe that the previous approaches do not gain many additional targets above a rank cutoff of 20, whereas PrediTALE still reports additional predictions with up-regulation in the RNA-seq experiment when increasing the rank cutoff. The ranking according to AUC again shows the best performance for PrediTALE, followed by TALgetter, Talvez and Target Finder. The general picture for PXO83 is similar, where PrediTALE yields the largest number of true positive predictions starting from rank 5, but also the largest number of rank 1 predictions (3). This is also reflected by the AUC values, which rank the approaches in the order of PrediTALE, TALgetter, Talvez and Target Finder.

**Fig 2.**
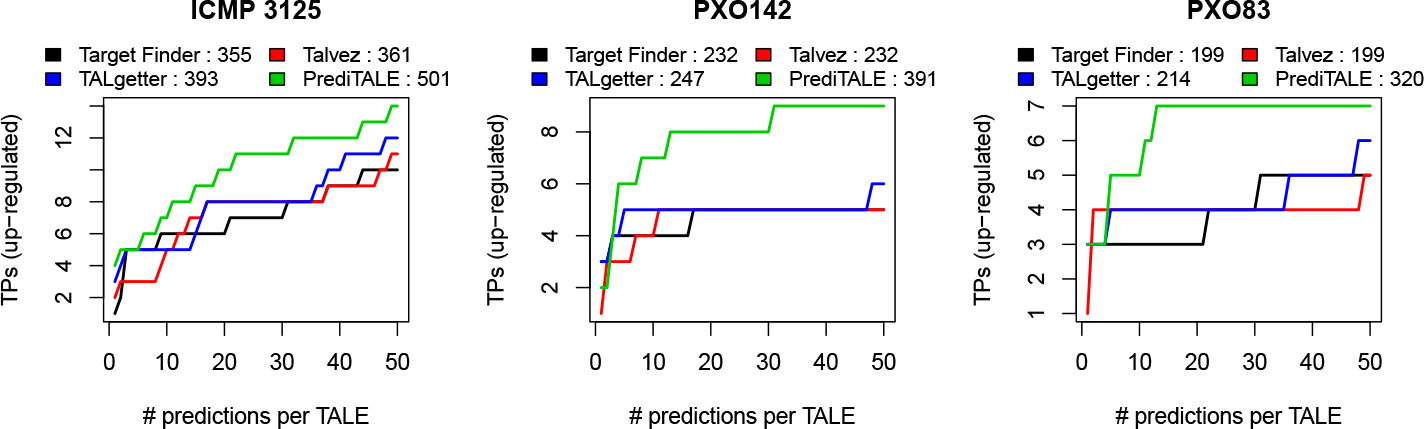
Performance evaluation on the level of target genes for three *Xoo* strains. For each approach, we plot the number of predicted target genes that are also up-regulated in the infection against the number of predicted target sites per TALE. In the legends, we further report the areas under the curves after the name of the individual approaches.

We take a different perspective on prediction results by assessing prediction performance on the level of TALEs. Specifically, we count the number of TALEs with at least one true positive target prediction for the same rank cutoffs as before. Again, PrediTALE identifies targets for a larger number of TALEs than the other approaches for the majority of rank cutoffs (Figure 3). However, we see notable differences between the different *Xoo* strain. For ICMP 3125^T^, PrediTALE is able to identify putative targets for 10 of its 17 TALEs. By contrast, the number of TALEs with a true positive prediction is lower for PXO142, where PrediTALE finds targets for at most 7 out of 19 TALEs, and for PXO83, where PrediTALE find targets for at most 7 out of 18 TALEs. As ICMP 3125^T^ has also been the strain with the largest number of differentially up-regulated genes (cf. Supplementary table A), the lower number of TALEs in PXO142 and PXO83 with a predicted target might be due to a different progression of the *Xanthomonas* infection.

**Fig 3.**
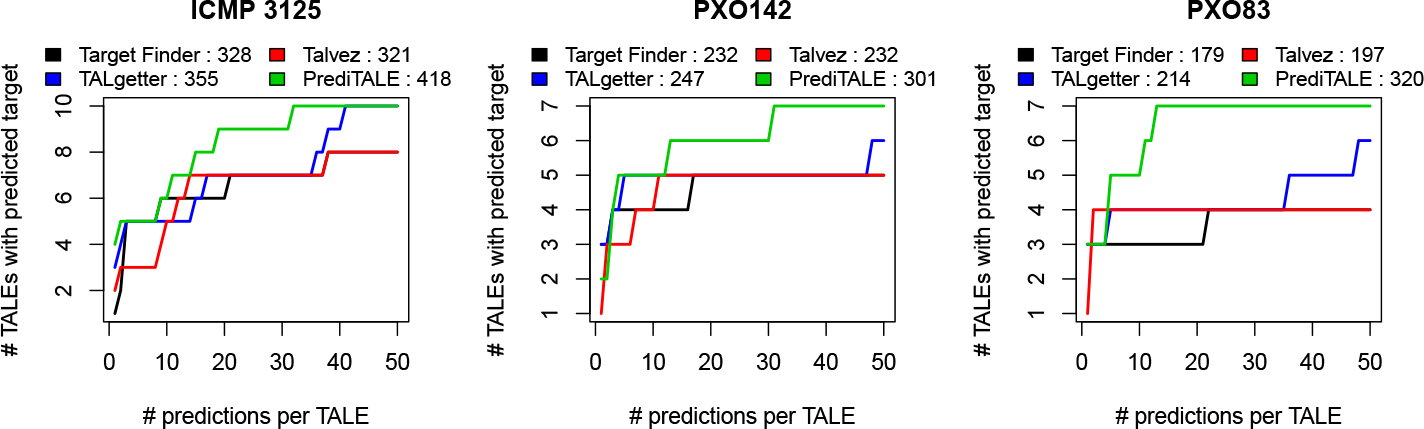
Performance evaluation on the level of TALEs for three *Xoo* strains. For each approach, we plot the number of TALEs with at least one predicted target gene that is also up-regulated in the infection against the number of predicted target sites per TALE.

We further summarize the data behind Figures 4 and 3 in Supplementary tables C and D, where we also report the average ranks of the four approaches across all three *Xoo* strains.

For sake of completeness, we also evaluate the four approaches for differentially up-regulated genes after *Xoo* infection based on the same FDR-based thresholds as for the *Xoc* experiments (Supplementary figure S3 and S4).

Although it has been shown that TALEs may activate transcription in both strand orientations relative to the transcription start site (TSS) of target genes [12, 13], a preference for the forward orientation has been postulated [13]. This is reflected by the strand penalty of PrediTALE, but no similar parameter exists for the previous approaches. Hence, above comparison might be perceived as partially unfair in favor of PrediTALE. For this reason, we repeat the benchmarking after restricting the predictions of all four appraoches to a forward orientation relative to the TSS (Supplementary figure S5 and S6). While the restriction to the forward strand has an effect on the number of target genes and TALEs with at least one true positive target, PrediTALE still yields an improved performance compared with the previous approaches over a wide range of rank cutoffs and, hence, achieves the largest AUC value of the four approaches in all cases.

For the ten *Xoc* strains, we find an improved prediction performance for PrediTALE as well. On the level of true positive target genes (Figure 4), PrediTALE yields the largest number of true positives for a rank cutoff of 1 for seven of the ten *Xoc* strains (cf. Table I). We also find an improved performance for the majority of the remaining rank cutoffs and *Xoc* strains. This improvement is especially pronounced for strains *Xoc* BLS279, CFBP7331, CFBP7341, and L8, whereas PrediTALE performs similar to or slightly worse than at least one of the previous approaches for *Xoc* CFBP7342 and RS105. For the remaining strains (B8-12, BLS256, BXOR1, CFBP2286), the improvement by PrediTALE is either rather small or mostly restricted to rank cutoffs of 20 or larger. This is also reflected by the areas under the curves, where PrediTALE yields the largest areas for B8-12, BLS256, BLS279, BXOR1, CFBP2286, CFBP7331, CFBP7341, L8, and also RS105, but nor for CFBP7342. Results are largely similar on the level of TALEs with at least one true positive predicted target (Supplementary figure S7), where PrediTALE yields the largest area under the curve for the same strains.

**Fig 4.**
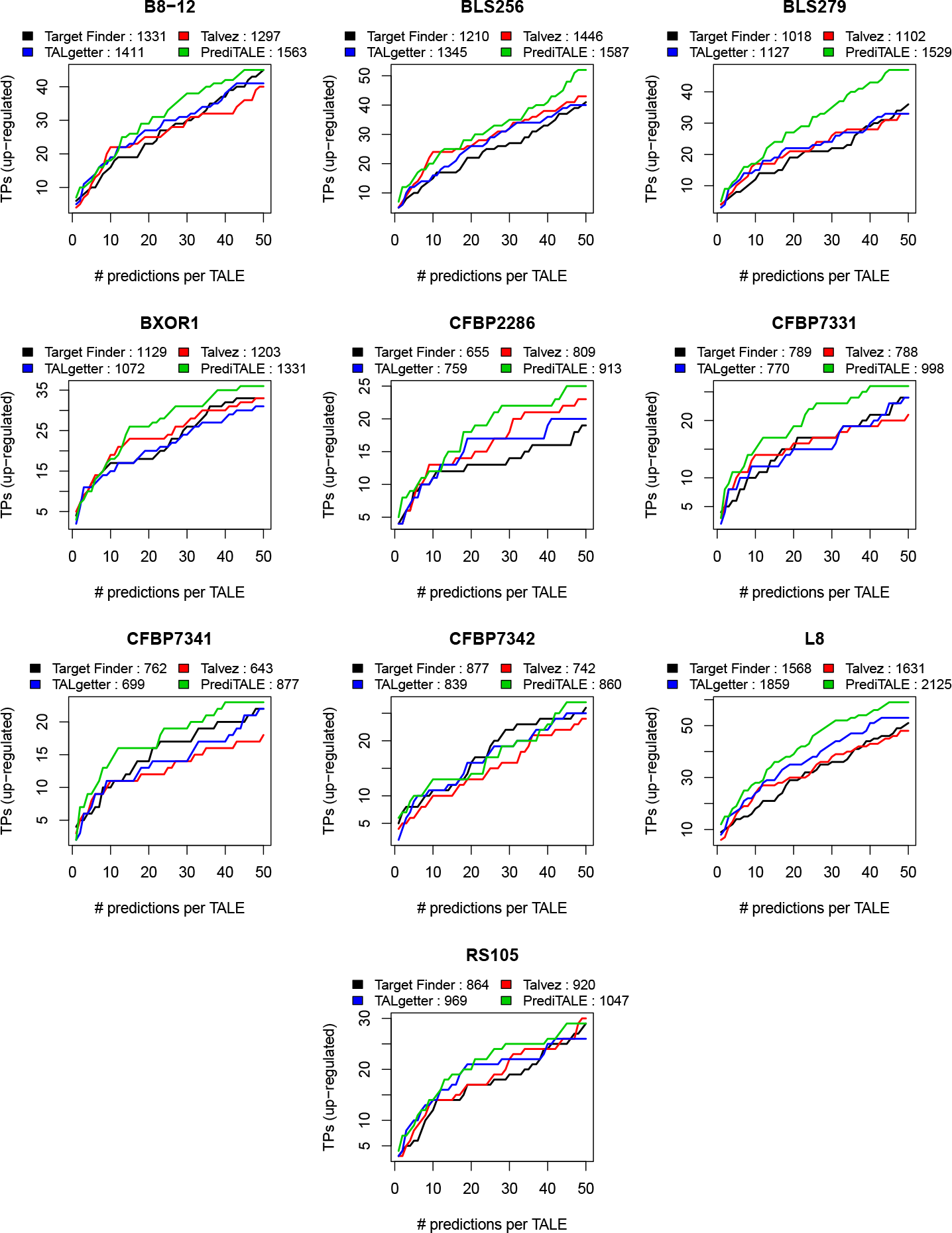
Performance evaluation on the level of target genes for 10 *Xoc* strains. For each approach, we plot the number of predicted target genes that are also up-regulated in the infection against the number of predicted target sites per TALE.

To obtain a more condensed overview on the results for the *Xoc* strains, we finally compute the average performance ranks across all ten *Xoc* strains for each of the four approaches and fixed rank cutoffs of 1, 10, 20, and 50, and for the area under the curve both on the level of target genes and on the level of TALEs (Table 1 and Supplementary tables I and J). For all rank cutoffs and the area under the curve, we observe that PrediTALE yields the best average rank with values betwen 1.1 and 1.5. We further assess the statistical significance of differences between the different tools by a Quade test, and the pairwise differences between tools by the associated post-hoc test (see Methods). This assessment is partly limited by the fact that pairs of *Xoc* strains may have identical TALEs in their TALEomes, which also means that the performance values of those strains are not truly independent. However, we did not find a clear relationship between the similarity of performance values obtained for the different strains and the similarity of the corresponding TALEomes. For this reason, we consider this dependency rather mild and favor this limited statistical assessment over the complete lack of it.

**Table 1.** Testing the significance of differences in prediction performance.

| measure | Target Finder | TALgetter | Talvez | PrediTALE | Quade | TALgetter vs TALESF | Talvez vs Target Finder | Talvez vs TALgetter | PrediTALE vs Target Finder | PrediTALE vs TALgetter | PrediTALE vs Talvez |
| --- | --- | --- | --- | --- | --- | --- | --- | --- | --- | --- | --- |
| Genes R1 | 1.8 | 3.1 | 2.4 | 1.5 | ** | — | - |  |  | +++ | ++ |
| Genes R10 | 3.6 | 2.5 | 1.6 | 1.5 | *** | + | +++ | + | +++ | + |  |
| Genes R20 | 3.3 | 2.1 | 2.9 | 1.3 | *** | +++ | + |  | +++ | ++ | +++ |
| Genes R50 | 2.3 | 3 | 3 | 1.1 | ** |  |  |  | +++ | +++ | +++ |
| Genes AUC | 3.1 | 2.8 | 3 | 1.1 | *** |  |  |  | +++ | +++ | +++ |
| TALEs R1 | 1.8 | 3.1 | 2.4 | 1.5 | ** | — | - |  |  | +++ | ++ |
| TALEs R10 | 3.5 | 2.1 | 1.6 | 1.5 | *** | + | +++ | ++ | +++ | + |  |
| TALEs R20 | 3.5 | 1.8 | 2.6 | 1.2 | *** | +++ | + | - | +++ |  | +++ |
| TALEs R50 | 2.6 | 2.6 | 2.8 | 1.4 | ** |  |  |  | ++ | ++ | +++ |
| TALEs AUC | 3.4 | 2.6 | 2.8 | 1.1 | *** | ++ | + |  | +++ | +++ | +++ |

For each tool and each measure (TALEs/Genes; rank cutoff), we report the average performance rank per tool, the significance of the Quade test (\*:< 0.05; \*\*:< 0.01; \*\*\*:< 0.001), and the significance of the pairwise comparison in a post-hoc test. Here, '+' and '-' indicate that the first tool has gained a significantly better or worse performance than the second one, respectively. The number of symbols encodes the significance level in analogy to the Quade test.

Consistent with the previous observations, we find that PrediTALE never performs significantly worse then any of the three previous approaches, whereas in many cases it performs significantly better, often with p-values below 0.001. Notable exceptions are a rank cutoff of 1, where PrediTALE does not perform significantly different from Target Finder, a rank cutoff of 10, where PrediTALE does not perform significantly different from Talvez, and on the level of TALEs, a rank cutoff of 20, where PrediTALE does not perform significantly different from TALgetter.

Repeating the same analysis for varied q-value threshold (Supplementary figures S8 and S9, Supplementary tables K, L, and M), for varied log fold change threshold (Supplementary figures S10 and S11, Supplementary tables N, O, and P), and for predictions restricted to the forward strand relative to the TSS (Supplementary figures S12 and S13, Supplementary tables Q, R, and S), benchmarking results are essentially similar to our previous findings. One notable exception is the Quade test for rank 1 predictions restricted to the forward strand (Supplementary table S), which is no longer significant. This means that none of the approaches studied yields significantly better rank 1 predictions than any other under this scenario.

In summary, we find i) that PrediTALE produces several unique predictions that might not have been considered based on previous approaches, ii) although low in absolute terms, the number of true positives among these predictions is often larger than for the previous aproaches, and iii) an assessment of the performance of PrediTALE across a wide range of rank cutoffs demonstrates that in most of the cases the application of PrediTALE yields a larger number of true positive target predictions than any of the three previous approaches.

### PrediTALE predicts novel putative target genes

As we have seen from Figure 1B, putative target genes with up-regulation after *Xoo* infection are often found in the intersection of the predictions of all four approaches. In addition, PrediTALE predicts several putative target genes of TALEs from the three *Xoo* strains that might have been neglected using one of the previous tools. In the following, we scrutinize the predictions for the *Xoo* strains with a focus on novel predictions, while we give a complete list of top 20 predictions of all four approaches including the ten *Xoc* strains in Supplementary table V.

In Table 2, we collect further information about those target genes including the corresponding log fold change and prediction ranks for all four approaches.

**Table 2.** Putative TALE target genes that are among the top 20 predictions per TALE for any of the four approaches.

| Gene | lfc | Target Finder | Talvez | Talgetter | PrediTALE | annotation |
| --- | --- | --- | --- | --- | --- | --- |
| **ICMP 3125<sup>T</sup>** |  |  |  |  |  |  |
| Os04g43730 | 5.762 | TalES1 (9) | TalAR13 (19);<br>TalES1 (10) | TalAR13 (564);<br>TalES1 (67) | TalAR13 (472);<br>TalES1 (108) | OsWAK51 |
| Os02g06670 | 3.815 | TalBA8 (1) | TalBA8 (2) | TalBA8 (1) | TalBA8 (1) | retrotransposon protein |
| Os09g29820 | 2.819 | TalAR13 (2) | TalAR13 (1) | TalAR13 (3) | TalAR13 (2) | bZIP transcription factor |
| Os03g51760 | 2.734 | TalAD22 (21) | TalAB16 (407);<br>TalAD22 (209) | TalAD22 (17) | TalAD22 (9) | OsFBX109 - F-box protein |
| Os04g05050 | 2.221 | TalAB16 (490) | NA | TalAB16 (63) | TalAB16 (11);<br>TalAH11 (824) | pectate lyase |
| Os01g40290 | 1.894 | TalAA15 (3) | TalAA15 (12) | TalAA15 (1) | TalAA15 (1) | expressed protein |
| Os05g45070 | 1.704 | NA | NA | TalAO15 (214) | TalAF17 (559);<br>TalAO15 (15) | harpin-induced protein 1 |
| Os11g26790 | 1.695 | TalAH11 (3) | TalAH11 (1) | TalAH11 (1);<br>TalAQ14 (559) | TalAH11 (1) | dehydrin |
| Os06g03710 | 1.591 | TalES1 (44) | TalES1 (81) | TalES1 (41) | TalES1 (19) | DELLA protein SLR1 |
| Os03g03034 | 1.295 | TalAO15 (404);<br>TalAQ14 (125) | TalAO15 (396);<br>TalAQ14 (9) | TalAB16 (600);<br>TalAO15 (566);<br>TalAQ14 (15) | TalAB16 (220);<br>TalAO15 (556);<br>TalAQ14 (32) | flavonol synthase |
| Os01g73890 | 1.079 | TalBM2 (3) | TalBM2 (14) | TalBM2 (2) | TalBM2 (1);<br>TalET1 (477) | transcription initiation factor IIA gamma |
| Os10g28240 | 0.918 | TalAR13 (71) | TalAR13 (47) | TalAR13 (16) | TalAR13 (6) | calcium-transporting ATPase |
| Os09g07460 | 0.746 | TalBA8 (88) | TalBA8 (17) | TalBA8 (48) | TalBA8 (22) | kelch repeat protein |
| **PXO142** |  |  |  |  |  |  |
| Os02g49350 | 5.163 | TalBH2 (1) | TalBH2 (2) | TalBH2 (5) | TalBH2 (8) | plastocyanin-like |
| Os03g09150 | 2.530 | NA | NA | TalBK2 (805) | TalBH2 (4);<br>TalBK2 (239) | pumilio-family RNA binding |
| Os11g31190 | 2.514 | TalAN15 (681) | TalAE16 (530);<br>TalBH2 (848) | TalAQ15 (660);<br>TalBH2 (144) | TalBH2 (3) | SWEET14 (nodulin MtN3) |
| Os09g29820 | 2.272 | TalAR14 (1) | TalAR14 (2) | TalAR14 (1) | TalAR14 (3) | bZIP transcription factor |
| Os03g51760 | 1.368 | TalAD23 (77) | TalAD23 (288) | TalAD23 (48) | TalAD23 (13);<br>TalAS12 (421) | OsFBX109 - F-box protein |
| Os01g40290 | 0.887 | TalAA16 (3) | TalAA16 (7) | TalAA16 (1) | TalAA16 (1) | expressed protein |
| Os06g29790 | 0.833 | TalAO16 (17) | TalAO16 (11) | TalAO16 (3) | TalAO16 (4);<br>TalAP15 (799) | phosphate transporter 1 |
| Os07g06970 | 0.824 | TalAP15 (1);<br>TalAQ15 (521) | TalAP15 (1);<br>TalAQ15 (319) | TalAP15 (1);<br>TalAR14 (563) | TalAI17 (889);<br>TalAP15 (1) | HEN1 |
| **PXO83** |  |  |  |  |  |  |
| Os09g29820 | 2.82 | TalAR3 (1) | TalAR3 (2) | TalAR3 (1) | TalAR3 (5) | bZIP transcription factor |
| Os02g06670 | 2.74 | TalBA2 (1) | TalBA2 (2) | TalAR3 (996);<br>TalBA2 (1) | TalAR3 (83);<br>TalBA2 (1) | retrotransposon protein |
| Os03g51760 | 1.91 | TalAD5 (77) | TalAB5 (407);<br>TalAD5 (288) | TalAD5 (48) | TalAD5 (13) | OsFBX109 - F-box protein |
| Os04g19960 | 1.70 | NA | TalAN3 (668);<br>TalAP3 (365) | TalAP3 (588) | TalAC5 (1);<br>TalAN3 (846) | retrotransposon protein |
| Os04g05050 | 1.62 | TalAB5 (490) | TalAP3 (931) | TalAB5 (63) | TalAB5 (11) | pectate lyase |
| Os07g06970 | 1.40 | TalAP3 (1) | TalAP3 (1);<br>TalAQ3 (512) | TalAP3 (1);<br>TalAR3 (988) | TalAP3 (1) | HEN1 |
| Os03g03034 | 1.18 | TalAO3 (404);<br>TalAQ3 (70) | TalAO3 (396);<br>TalAQ3 (2) | TalAB5 (600);<br>TalAO3 (566);<br>TalAQ3 (5) | TalAB5 (220);<br>TalAO3 (556);<br>TalAQ3 (5) | flavonol synthase |

For each *Xoo* strain, we list the gene ID (MSU7) and the log fold change (lfc) in the corresponding RNA-seq experiment. For each of the four approaches, we further list the TALE(s), for which a gene has been predicted as a target and in parentheses the corresponding prediction rank. An “NA” entry for a combination of gene and prediction approach indicates that this gene has not been among the top 1000 predictions for any TALE.

The target genes in the intersections of the predictions of all four approaches comprise several well known targets: Os09g29820 (OsTFX1), a bZIP transcription factor, is targeted by TALEs from class TalAR with members in all three *Xoo* strains (Supplementary figure S14) and has been proposed as a TALE target early [5, 42]. Os01g40290 [5], an expressed protein without annotated function, is targeted by TalAA members, which are also present in all three *Xoo* strains. However, this gene is not in the list of predictions for *Xoo* PXO83, because the corresponding p-value was larger than the threshold of 0.05 (Supplementary table T). Os01g73890 (TFIIA*γ*) [5], that has been shown to promote TALE function [43], is targeted by TalBM2 in ICMP 3125^T^. In concordance to TalBM class members missing in PXO142 and PXO83, Os01g73890 shows no up-regulation in these two strains. Os07g06970 (HEN1) has also been among the first TALE target genes proposed [5] and is targeted by TalAP members present in all three *Xoo* strains, but falls below the threshold on the log fold change by a small margin in ICMP 3125^T^ (Supplementary figure S15). Os06g29790 [18], a phosphate transporter, has been predicted as a target of TalAO16 from PXO142 by all four approaches, but not for the TalAO members from PXO83 and ICMP 3125^T^, which have a slightly different and longer RVD sequence. In the RNA-seq data, however, we find the strongest up-regulation for ICMP 3125^T^, although Os06g29790 appears only on rank 49 of the PrediTALE predictions for TalAO15 from that strain. Hence, experimental data and computational predictions are partly contradictory in this case.

In addition, we find several putative target genes in the intersection that have not been reported before: Os02g06670, a retrotransposon protein, is predicted as a target of TalBA8 and TalBA2 in ICMP 3125^T^ and PXO83, respectively, whereas PXO142 lacks a TalBA member. Nonetheless, Os02g06670 is up-regulated after PXO142 infection, although to a lesser degree than in the other two strains (cf. Supplementary figure S15). Os11g26790 (RAB21), a dehydrin that has been shown to play a role in drought tolerance related to pathogen infection [44], is predicted to be targeted by TalAH11 from ICMP 3125^T^. Os11g26790 is up-regulated for ICMP 3125^T^ but also for PXO142 (Supplementary figure S15), although in the latter case, the corresponding p-value is again not significant. Os02g49350, a plastocyanin-like protein, is strongly up-regulated only in PXO142 and predicted as a target of TalBH2, where class TalBH is exclusive to PXO142 among the strains studied.

Finally, we find several putative target genes that have been predicted only by a subset of approaches: For ICMP 3125^T^, Os04g43730 (OsWAK51) is among the top 20 predictions for TalES1 only for Target Finder and Talvez. In turn, PrediTALE predicts Os06g03710 (DELLA protein SLR1) as a TalES1 target on rank 19, which appears on later ranks for the other approaches. Os04g43730 is induced more strongly than Os06g03710 and exclusively in ICMP 3125^T^, which renders this the more likely target. Os03g51760 (OsFBX109) is among the top 20 predictions for TalAD members only for PrediTALE. Due to variations in their RVD sequence, TALgetter has this in the top 20 predictions only for TalAD22 in ICMP 3125^T^, but not for the other strains. As Os03g51760 is clearly up-regulated after infection with any of the three *Xoo* strains (Supplementary figure S15), this is likely a true TalAD target.

Talvez and TALgetter have Os03g03034, annotated as a flavonol synthase, among their top 20 predictions for TalAQ members in ICMP 3125^T^ and PXO83, while this gene is among the top 20 predictions of PrediTALE only for TalAQ3 in PXO83 due to differences in RVD sequence. In PXO142, TalAQ15 is annotated as a pseudo gene and this pattern is also reflected by the RNA-seq data. Os03g03034 has been proposed to be a TALE target before [5].

Os04g05050, annotated as a pectate lyase, is only among the top 20 predictions of PrediTALE in ICMP 3125^T^ (TalAB16) and PXO83 (TalAB5), whereas this gene is ranked substantially lower (rank 83) for TalAB8 from PXO142 by PrediTALE as well. From the RNA-seq data, we find that Os04g05050 is up-regulated in all three *Xoo* strains, although the level of up-regulation is lower for PXO142 than for the other two strains.

Os05g45070, annotated as hairpin-induced protein 1, is predicted only by PrediTALE as an alternative target of TalAO15 in ICMP 3125^T^ and shows clear up-regulation only after infection with this *Xoo* strain. Os10g28240, a calcium transporting ATPase, is predicted by TALgetter and PrediTALE as target of TalAR13 of ICMP 3125^T^ but, on later ranks, also by the other two approaches, and is up-regulated exclusively after ICMP 3125^T^ infection. Os09g07460, a kelch repeat protein, is only among the top 20 predictions of Talvez for TalBA and on later ranks for the other approaches. This gene is up-regulated only in ICMP 3125^T^, although not strongly.

For PXO142, we find two further putative targets of TalBH2 that are predicted exclusively by PrediTALE: Os03g09150 (pumilio-family RNA binding) is up-regulated in PXO142 but also in PXO83, for which it does not appear among the top 20 predictions of any approach. Os11g31190 (Os11N3, OsSWEET14) is a well known target [45, 46], which is predicted here also for TalBH exclusively by PrediTALE due to its ability to adequately handle the aberrant repeat [6] of TalBH2. Os11g31190 is also known to be targeted by TalAC members (previously termed AvrXa7) [42] including TalAC5 in PXO83 and, hence, is strongly up-regulated after PXO83 infection as well. However, in this case all approaches fail to predict this target due to the large number of mis-matches in the target box [6], even accounting for the aberrant repeat in TalAC5.

Instead, another retrotransposon protein (Os04g19960) is the top prediction of PrediTALE for TalAC5 from PXO83, which is confirmed by RNA-seq data as this gene is strongly up-regulated after PXO83 infection but not after infection with one of the other strains.

In summary, we find several novel putative target genes of which 6 are highly promising (Os11g26790, Os02g49350, Os03g51760, Os04g05050, Os05g45070, Os04g19960), where 4 of these (Os03g51760, Os04g05050, Os05g45070, Os04g19960) are predicted on high ranks exclusively by PrediTALE. Recently, we could experimentally validate the targets Os04g43730 (OsWAK51), Os06g29790 (phosphate transporter), Os03g51760 (OsFBX109), Os03g03034 (flavonol synthase), and Os04g05050 (pectate lyase) by qRT-PCR using a TALE-less strain (Roth X1-8) complemented with individual TALEs [47].:

### Orphan TALEs

We also observe from Figure 3 and Supplementary figure S7 that for many strains, neither of the approaches considered is able to identify a putative target genes for all TALEs present in their TALEome. We term such TALEs without reasonable target prediction *orphan TALEs*, and we will discuss these in more detail in the following.

More precisely, we call a TALE or a TALE class *orphan* if there is no up-regulated gene among the top 50 predictions of any of the four approaches. Furthermore, we check if this pattern is consistent for the TALEs from a common TALE class across almost all *Xoo* and *Xoc* strains studied.

We find as orphan the TALE classes present in all three *Xoo* strains TalAF, TalAI and TalAN. In addition, TalAG (PXO142, PXO83), TalAL (PXO142), TalAS (PXO142, PXO83), TalBJ (PXO83), TalCA (PXO83), TalET (ICMP 3125^T^), and TalDR (PXO142) are orphan TALE classes in individual *Xoo* strains. The TALEs from class TalAI and TalDR are truncTALEs that are lacking large parts of the C-terminus including the activation domain and, for this reason, do not act as transcriptional activators. TruncTALEs have been found to function as suppressors of resistance mediated by an immune receptor [48].

In the *Xoc* strains, however, TalAF is not orphan as we find putative target genes among the top 50 predictions for the class members present in B8-12 and L8. For TalAZ, we find a target for TalAZ7 from *Xoc* L8, but not for the other 7 *Xoc* strains harboring TalAZ TALEs. In addition, we consider TalCQ1 from BXOR1 and TalCR1 (CFBP7331) and TalCR2 (CFBP7341) as orphan.

Reasons for orphan TALEs could be manifold. First of all, we cannot be sure that these TALEs are indeed expressed by the bacteria and are secreted into the host plant cells. Second, some TALEs might activate target genes slower or to a lesser degree than others and, for this reason, target gene activation might not be detectable, yet, in the RNA-seq experiments, especially at the 24h timepoint chosen for *Xoo*. Third, these TALEs might target specific variants of boxes in promoters of rice lines that are not represented by the *O. sativa* Nipponbare reference genome, or might even target genes in alternative host plants, e.g., grasses in the vicinity of fields where rice is grown. Fourth, these TALEs might target genes that are missing from the current gene annotations of rice. Such targets would be neglected by the current approach to specifically scan promoter sequences of annotated genes for putative TALE boxes. To address the latter issue, we switch to an alternative approach in the following. Here, we perform *genome-wide* scans for putative target boxes instead, and search for differentially expressed regions in the vicinity of putative target boxes predicted anywhere in the reference genome.

### Genome-wide prediction profiles discover potential novel target genes

We perform genome-wide predictions of TALE target boxes in *Oryza sativa* Nipponbare (MSU7) for the 256 *Xoc* TALEs from 10 strains and 54 *Xoo* TALEs from 3 strains and check for differentially expressed regions near the predicted target boxes. Differential expression is based on the mapped RNA-seq data after infection with the respective *Xoo* and *Xoc* strains.

After infection with *Xoo* strains, 14 TALEs are found to have differentially expressed regions near at least one predicted target box. Table 3 lists the total number of 19 TALE target boxes together with MSU7 gene annotations overlapping the differentially expressed regions. Notably, 15 of these targets have already been reported in subsection “PrediTALE predicts novel putative target genes” when restricting the search to promoter regions of annotated genes. However, for two genes, target boxes from other TALs were predicted in case of genome-wide scan. The expression of the pectate lyase precursor (Os04g05050) was up-regulated by TalAB5 according to promotor prediction, but the genome-wide prediction contains the same gene up-regulated by TalAD22. The same scenario for the phosphate transporter 1 (Os06g29790), which according to promotor predictions is up-regulated by TalAO16 and TalAP15. However, in the genome-wide scans, a target box of TalAH11 was predicted. The genome-wide scan i) does not make use of gene annotations, and ii) could be expected to be more prone to false positive predictions than the restricted search in promoters. Hence, the fact that many predictions re-occur in the genome-wide scan demonstrates the general utility of this approach.

**Table 3.** Genome-wide prediction of *Xoo* TALE targets with PrediTALE.

| TALE | Chr | Pos. box | Gene | Annotation | PiP |
| --- | --- | --- | --- | --- | --- |
| **ICMP 3125<sup>T</sup>** |  |  |  |  |  |
| TalAA15 | Chr1 | 22747303 | Os01g40290 | expressed protein | yes |
| TalAD22 | Chr3 | 29685233 | Os03g51760 | OsFBX109 - F-box protein | yes |
| TalAD22 | Chr4 | 2486797 | Os04g05050 | pectate lyase precursor | TalAB5 |
| TalAH11 | Chr6 | 17129738 | Os06g29790 | phosphate transporter 1 | TalAO16,<br>TalAP15 |
| TalAN14 | Chr2 | 31931460 | Os02g52170 | expressed protein | no |
| TalAN14 | Chr8 | 19950534 | Os08g32160 | oxidoreductase, 2OG-FeII oxygenase | no |
| TalAR13 | Chr10 | 14685398 | Os10g28240 | calcium-transporting ATPase | yes |
| TalAR13 | Chr9 | 18123472 | Os09g29820 | bZIP transcription factor | yes |
| TalBA8 | Chr2 | 3353526 | Os02g06670 | retrotransposon protein | yes |
| TalBM2 | Chr1 | 42819000 | Os01g73890 | transcription initiation factor IIA gamma | yes |
| **PXO142** |  |  |  |  |  |
| TalAO16 | Chr7 | 22546154 | – | – | NA |
| TalAR14 | Chr5 | 16047774 | Os05g27580 | wound-induced protein WI12 | no |
| TalAR14 | Chr9 | 18123472 | Os09g29820 | bZIP transcription factor | yes |
| TalBH2 | Chr11 | 18174482 | Os11g31190 | SWEET14 (nodulin MtN3 family) | yes |
| TalBH2 | Chr2 | 30158664 | Os02g49350 | plastocyanin-like | yes |
| **PXO83** |  |  |  |  |  |
| TalAC5 | Chr4 | 11130506 | Os04g19960 | retrotransposon protein | yes |
| TalAP3 | Chr7 | 3434725 | Os07g06970 | HEN1 | yes |
| TalAQ3 | Chr3 | 1245017 | Os03g03034 | flavonol synthase/flavanone 3-hydroxylase | yes |
| TalAR3 | Chr9 | 18123472 | Os09g29820 | bZIP transcription factor | yes |

Genome-wide prediction of *Xoo* TALE targets using PrediTALE filtered for differentially expressed regions within 3000 bp surrounding the target box. For each *Xoo* strain, we list the TALE name, Chromosome number and position of the target box (Pos. box) in *Oryza sativa* Nipponbare genome, and the annotated MSU7 Gene ID and description (if present). In addition, the last column contains the information, whether predictions in promoters (PiP) also report this target.

In addition to those targets reported previously, we find three novel target boxes in the vicinity of differentially expressed regions that overlap annotated genes, including a wound-induced protein and an oxidoreductase. For TalAO16 from PXO142, we find a differentially expressed region next to a predicted target box on chromosome 7 with no annotation in MSU7 (Supplementary Figure S16; complete list in Supplementary Table W). For this reason, we extracted the sequence under the differentially expressed region, and first compared it against the NCBI protein database ‘nr’ using blastx but received no matching result. We additionally compared this sequence against the NCBI reference RNA sequences (refseq rna) using blastn, which resulted in a highly significant hit for XR 001547425.2, a predicted long non-coding RNA.

Upon infection of rice with *Xoc* strains, differentially expressed regions near at least one predicted target box were found for 26 of 28 (B8-12), 28 of 28 (BLS256), 25 of 26 (BLS279), 26 of 27 (BXOR1), 22 of 28 (CFBP2286), 19 of 22 (CFBP7331), 19 of 21 (CFBP7341), 18 of 23 (CFBP7342), 27 of 29 (L8) and 19 of 24 (RS105) TALEs. Supplementary Table X lists all genome-wide predicted targets in the vicinity of differentially expressed regions of these *Xoc* strains.

In the following, we will discuss two example regions in detail. As discussed in the previous section, TalAZ appears to be an orphan TALE based on the promoterome-wide scans for target boxes. However, based on genome-wide scans, we find a differentially expressed region, which could constitute a target gene of TalAZ, on Chr4 (Figure 5). Only 8 of the 10 *Xoc* strains studied have a TalAZ member in their TALEome. The profile plots clearly show that the region of interest is only differentially expressed after infection with these 8 strains harbouring TalAZ members. Performing blast searches of the differentially expressed sequences, we received a hit for XP 015634381.1, a sulfated surface glycoprotein 185 [Oryza sativa Japonica Group], which has been added to the IRGSP-1.0 annotation at NCBI but was not present in MSU7.

**Fig 5.**
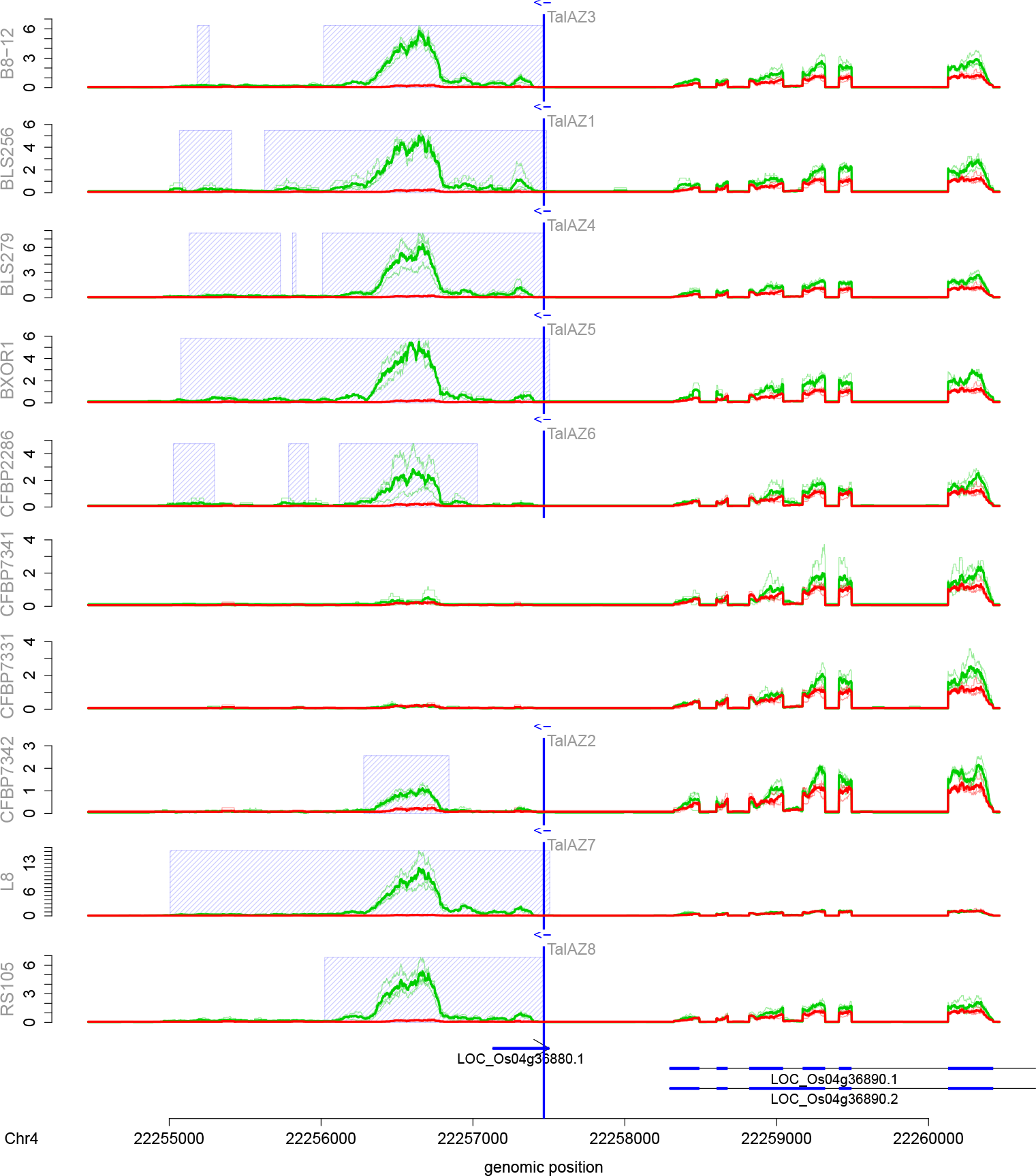
Genome-wide predictions of TalAZ in *Oryza sativa* Nipponbare profile for 10 *Xoc* strains in the area of the TalAZ target box. RNA-seq coverage after inoculation (green line) is compared with mock control (red line). In addition, we show the average of individual replicates of control and treatment are summarized as thick lines. The blue shaded boxes mark the differentially expressed regions. The arrows under the profiles reflect the MSU7 annotation within the genomic region. The genomic position of the TALE target box is marked by a vertical blue line.

As a second example, we consider a putative TalBD target on Chr6. The profile plots (Figure 6) show differentially expressed regions in all 10 strains. However, a blastx search of the respective sequences, spanning two larger differentially expression regions, provides no clear result. Matches include an Auxin-responsive protein IAA22 (Q69TU6.1) and different bromodomain-containing factors (XP 006659043.1, XP 025882131.1 XP 015650662.1). As drops in the coverage profiles and split reads in the mapping indicate the existence of introns within the differentially expressed regions, we additionally compare the spliced sequence using blastn against the NCBI reference RNA sequences. The result contains a predicted non-conding RNA (XR 003242961.1) and different transcript variants of a predicted mRNA, coding for bromodomain-containing factors (XM 015840709.1, XM 015840708.1, XM 006658980.2, XM 026026346.1, XM 015795177.2, XM 015795176.2).

In summary, our results demonstrate that genome-wide prediction of target boxes using PrediTALE enables us to identify novel targets independently of existing gene annotations including previously missing non-coding RNAs.

**Fig 6.**
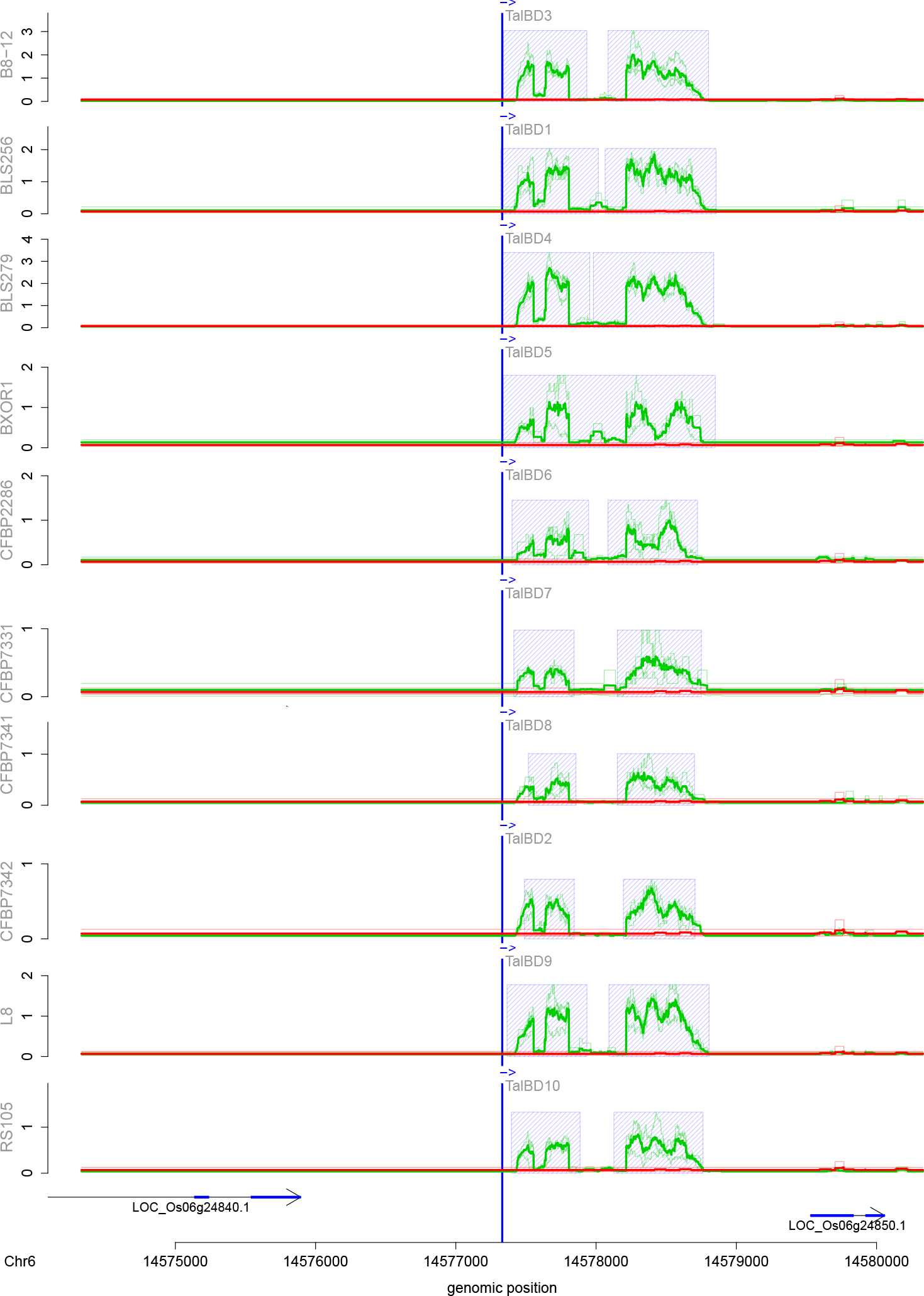
Genome-wide predictions of TalBD in *Oryza sativa* Nipponbare profile for 10 *Xoc* strains in the area of the TalBD target box. RNA-seq coverage after inoculation (green line) is compared with mock control (red line). In addition, we show the average of individual replicates of control and treatment are summarized as thick lines. The blue shaded boxes mark the differentially expressed regions. The arrows under the profiles reflect the MSU7 annotation within the genomic region. The genomic position of the TALE target box is marked by a vertical blue line.

## Conclusion

Accurate computational predictions of TALE target boxes are required for elucidating virulence targets of TALEs that support bacterial infection of host plants. In this paper, we present PrediTALE, a novel approach for predicting target boxes based on a TALE’s RVD sequence. Since the publication of all previous approaches [14, 16, 18], our understanding of mechanisms and principles of TALE targeting has increased substantially. Specifically, it has been shown that repeats of aberrant lengths may compensate for frame shifts in target boxes [6], that activation of gene expression by TALEs binding to the reverse strand is possible, but rare [13]. In addition, quantitative data about virtually all combinations of AAs at RVD positions have been collected [19, 21–25]. All these insights have been integrated into PrediTALE either as part of the model or as training data that are used to adapt model parameters. Here, we demonstrate that PrediTALE predicts TALE targets with improved accuracy compared with previous approaches, where ground truth is derived from in-house and public RNA-seq data after *Xoo* and *Xoc* infection. However, our results also confirm that any of the current computational approaches suffers from false positive predictions and, hence, experimental support of predicted targets is inevitable.

PrediTALE predicts several unique target genes, several of which are highly promising for further experimental validation. While RNA-seq data supports that these are activated by TALEs *in planta*, their importance for the infection process still needs to be investigated.

Given the improved accuracy and acceptable runtime of PrediTALE, we broaden the scope of computational predictions. Previously, predictions have been mostly limited to putative promoter regions of annotated genes. Here, we consider genome-wide predictions instead. We demonstrate that targets reported from promoterome-wide predictions are also recovered in genome-wide scans, but we also find differentially expressed regions at loci that do not overlap with annotated genes. These could be either protein-coding genes that are missing from the current annotation, but also include putative non-coding RNAs, which might have regulatory activity or other functions that foster bacterial infection.

To promote future research in plant-pathogen interactions related to TALEs, we make our methods available to the scientific community as open-source software tools.

## Acknowledgments

We thank Sebastian Becker for valuable discussions.

## Funding

This work was supported by grants from the Deutsche Forschungsgemeinschaft (BO 1496/8-1 to JB and GR 4587/1-1 to JG) and a grant from the European Regional Development Fund of the European Commission to JB and by the COST action FA1208 “SUSTAIN”.

