## Supplementary Figures S1 - S16; Supplementary Tables A - S for "PrediTALE: A novel model learned from quantitative data allows for new perspectives on TALE targeting"

### Supplementary Text S1 – Preprocessing of training data

#### Cong et al.

Pairs of TALE RVD sequence and tested target boxes were obtained from Fig. 1 and Fig. 2a of [1]. Data were grouped by TALE, and the global weight was computed as the maximum "Normalized reporter activation" for the current TALE divided by the maximum "Normalized reporter activation" reported for all TALEs with the same 13th AA at the varied positions. Target values were computed as the "Normalized reporter activation" of the current pair of TALE and target box divided by maximum "Normalized reporter activation" over all tested target boxes for the current TALE.

#### Streubel et al.

Pairs of TALE RVD sequence and tested target boxes were obtained from [2]. Data were grouped by TALE, and the global weight was computed as the maximum GUS activity for the current TALE divided by the maximum GUS activity reported for all TALEs with the same 13th AA at the varied positions stemming from the same experiment. Target values were computed as the GUS activity of the current pair of TALE and target box divided by maximum GUS activity over all tested target boxes for the current TALE.

#### Schreiber et al.

Pairs of TALE RVD sequence and tested target boxes were obtained from [3]. Data were grouped by TALE, and the global weight was computed as the maximum GUS activity for the current TALE divided by the maximum GUS activity reported for all TALEs stemming from the same experiment (corresponding to the same (sub-)figure in [3]). Target values were computed as the GUS activity of the current pair of TALE and target box divided by maximum GUS activity over all tested target boxes for the current TALE.

#### Yang et al.

Pairs of TALE RVD sequence and tested target boxes from [4] were provided by Wensheng Wei. Data were grouped by TALE, and the global weight was computed as the maximum EGFP activity for the current TALE divided by the maximum EGFP activity reported for all TALEs with the same 13th AA at the varied positions. Target values were computed as the EGFP activity of the current pair of TALE and target box divided by maximum EGFP activity over all tested target boxes for the current TALE.

#### Miller et al.

Pairs of TALE RVD sequence and tested target boxes were obtained from Supplementary Table 2 of [5]. Data were grouped by TALE, and the global weight was computed as the maximum "Normalized ELISA score" for the current TALE divided by the maximum "Normalized ELISA score" reported for all TALEs with the same 13th AA at the varied positions. Target values were computed as the "Normalized ELISA score" of the current pair of TALE and target box divided by maximum "Normalized ELISA score" over all tested target boxes for the current TALE.

#### Rogers et al.

Probe sequences and binding intensities were obtained from [6] for 21 TALEs measured at different concentrations yielding a total of 55 PBM experiments. Each PBM experiment is accompanied by the RVD sequence of the corresponding TALE, where the number of RVDs ranges from 9 to 19. These 55 experiments were further filtered by data quality measured for each experiment individually.

Specifically, for probe sequence $i$, a score profile was computed using the model of TALgetter [7] based on the respective TALE and the log-probability of the best-matching sub-sequence on either the forward or the reverse complement strand of the probe sequence was stored as $s_i$. In addition, the number of sub-sequences with a log-probability larger than $s_i - \log(10)$ (i.e., those with a probability that is at most 10-fold lower than that of the best-matching sub-sequence) was stored as $n_i$. Using these measures, the Pearson correlation coefficient $\rho$ was computed based on scores and mean normalized log-intensities for each best-matching sub-sequence. To this end, the unique set of all best-matching sub-sequences $m_j$ was constructed and for each $m_j$ all probe sequences containing $m_j$ as the best-matching sub-sequence were collected, i.e., the $m_j$ define a partitioning on the probe sequences. For each partition belonging to one specific $m_j$, log-intensities $I_i$ were collected, divided by the corresponding $n_i$ and averaged over all probe sequences in the current partition, yielding mean log-intensity values $\bar{I}_{m_j}$. In addition, each $m_j$ has also been assigned a score by the TALgetter model denoted as $s_{m_j}$.

Pearson correlation $\rho$ was finally computed between the $\bar{I}_{m_j}$ and $s_{m_j}$ values. Only PBM data sets with $\rho > 0.6$ were retained.

In addition, probe sequences were partitioned into "positives" and "negatives" based on the PBM intensity values. Probe sequences with a log-intensity more than two standard deviations above the mean log-intensity of the current PBM experiment were assigned to the "positive" class (or the top 50 probe sequences if this rule yielded less than 50 sequences) and all remaining probe sequences were assigned to the "negative" class. Probe sequences were also scored by their log-probability according to the TALgetter model, i.e., by the log of the mean probability of all corresponding sub-sequences. These scores were then used as classification scores to compute the area under the precision-recall curve [8, 7] (AUC-PR) for the given binary classification problem. Only PBM experiments with an AUC-PR above 0.5 were retained for further analyes.

Probe sequences and PBM intensities of those experiments meeting both selection criteria were further processed to yield the final training data. Specifically, the unique best-matching sub-sequences $m_j$ and corresponding average log-intenties $I_{m_j}$ were collected, and the target values were set to the average log-intenties $I_{m_j}$ normalized to the maximum $I_{m_j}$ per PBM experiment. Global weights were defined identically for all $m_j$ from a common PBM experiment and set such that all PBM experiments yield the same total (i.e., summed) global weight and all PBM experiments together obtain a total global weight of 200, i.e., to a total global weight that is similar to that of 200 groups from one of the other experiments.

### Supplementary Figures

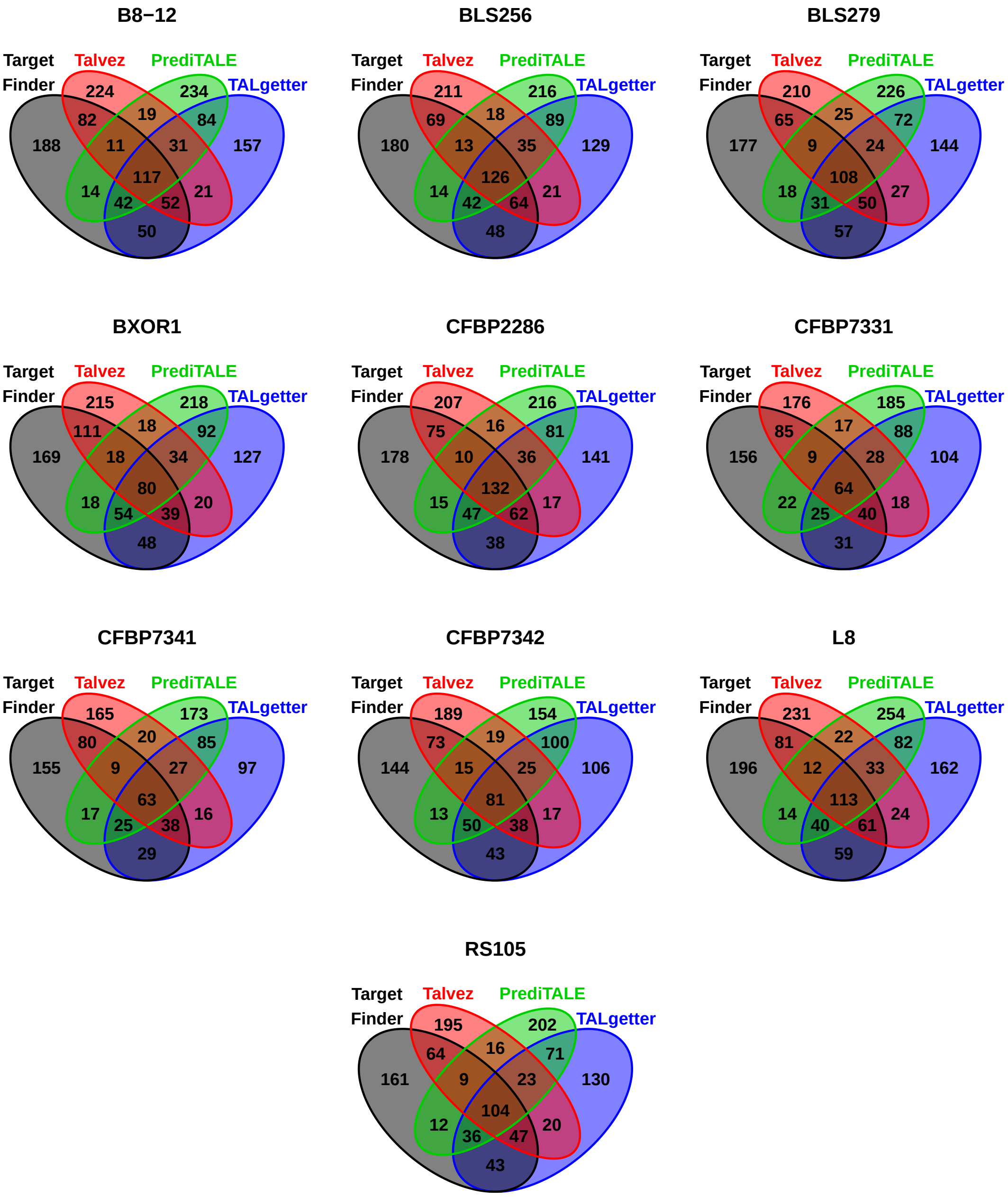


**Supplementary Figure S1.** Venn diagrams of predictions of the four approaches considered. For each *Xoc* strain and each approach, we consider the set of target genes obtained as the union of the top 20 predictions per TALE.

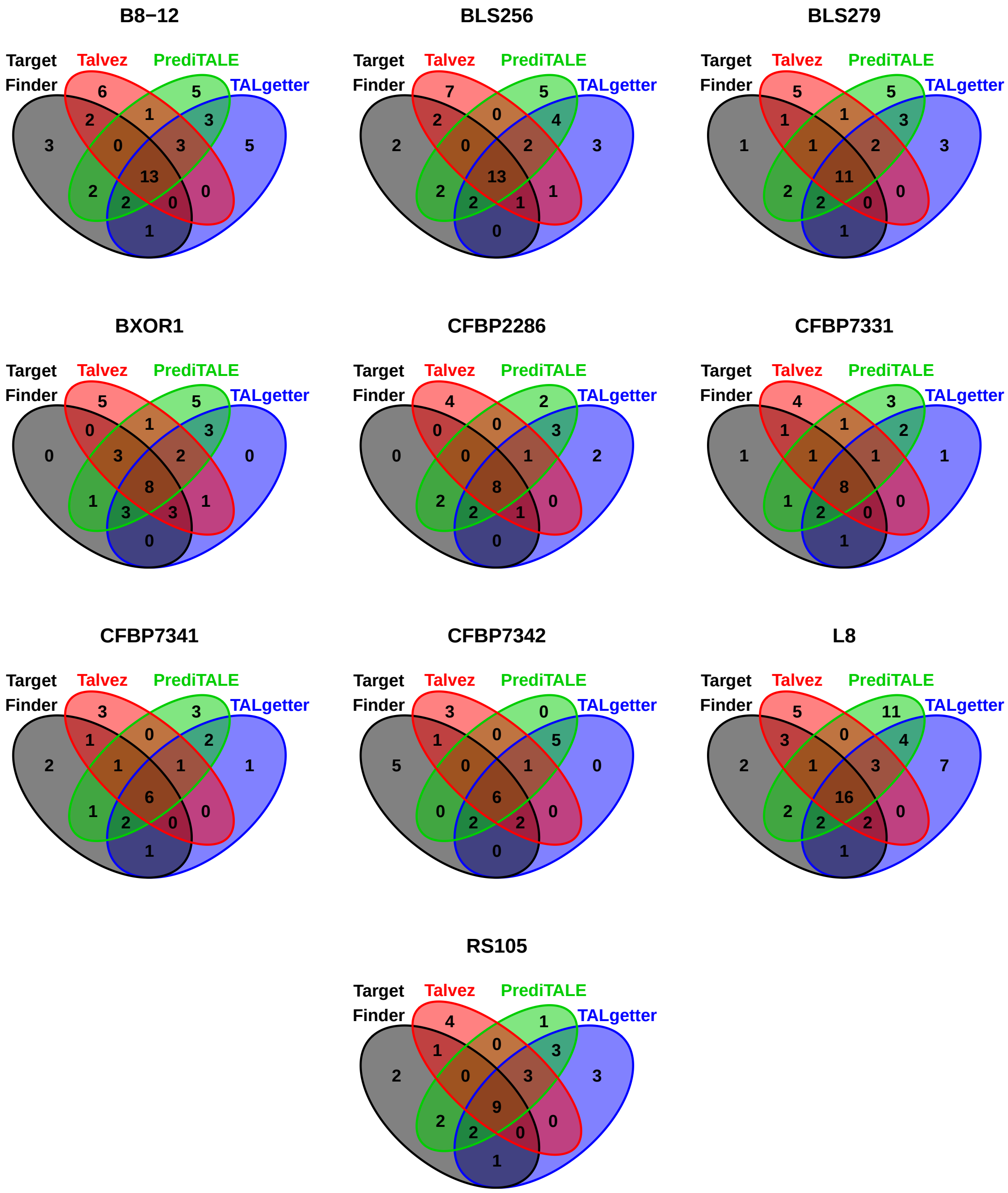


**Supplementary Figure S2.** Venn diagrams of true positive predictions of the four approaches considered. For each *Xoc* strain and each approach, we consider the set of target genes obtained as the union of the top 20 predictions per TALE. These sets are filtered by up-regulation of the corresponding genes according to RNA-seq data, and the resulting subsets are displayed.

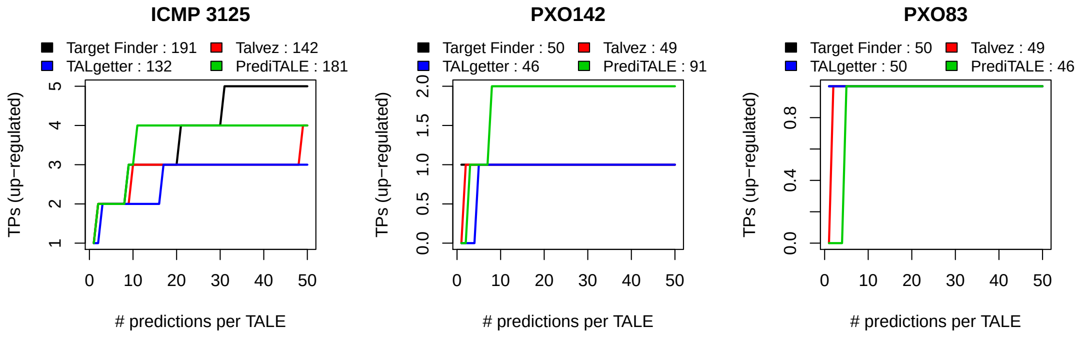


**Supplementary Figure S3.** Performance evaluation on the level of target genes for three *Xoo* strains. For each approach, we plot the number of predicted target genes that are also up-regulated in the infection (q-value $< 0.01$, log fold change $> 2$) against the number of predicted target sites per TALE.

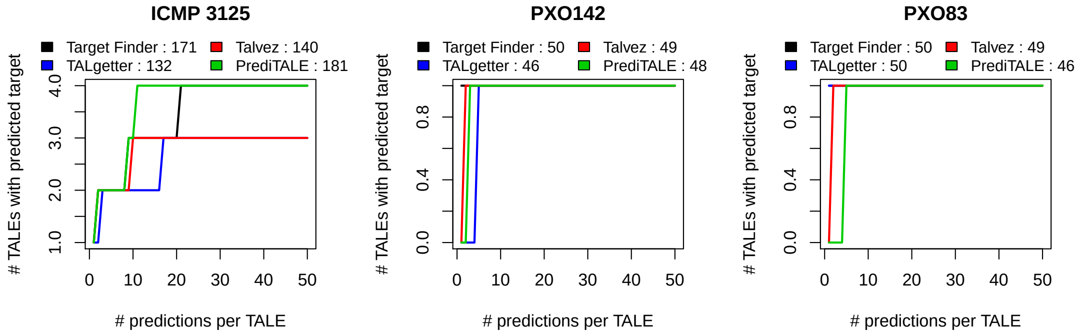


**Supplementary Figure S4.** Performance evaluation on the level of TALEs for three *Xoo* strains. For each approach, we plot the number of TALEs with at least one predicted target gene that is also up-regulated in the infection (q-value $< 0.01$, log fold change $> 2$) against the number of predicted target sites per TALE.

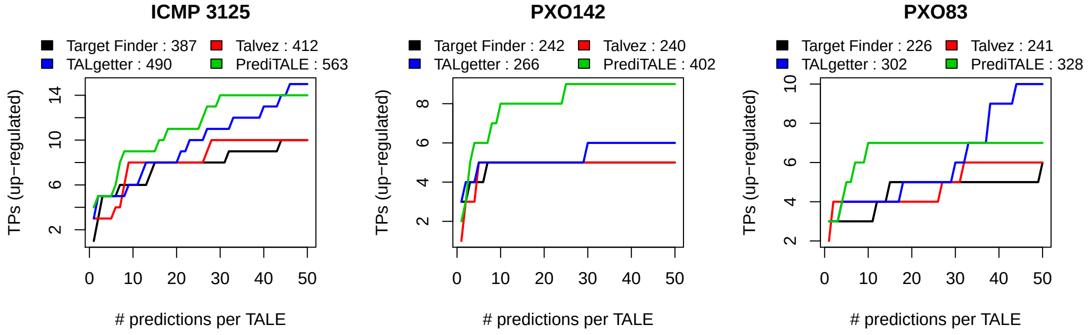


**Supplementary Figure S5.** Performance evaluation on the level of target genes for three *Xoo* strains when filtering for predictions of TALE boxes on the same strand as the downstream gene. For each approach, we plot the number of predicted target genes that are also up-regulated in the infection against the number of predicted target sites per TALE.

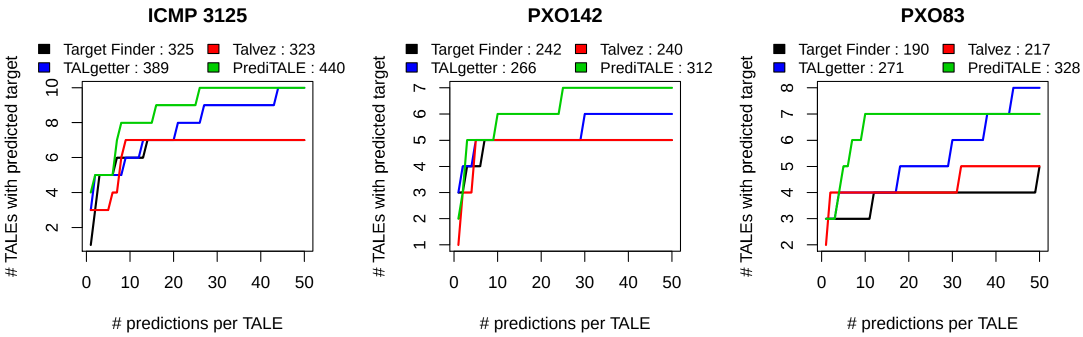


**Supplementary Figure S6.** Performance evaluation on the level of TALEs for three *Xoo* strains when filtering for predictions of TALE boxes on the same strand as the downstream gene. For each approach, we plot the number of TALEs with at least one predicted target gene that is also up-regulated in the infection against the number of predicted target sites per TALE.

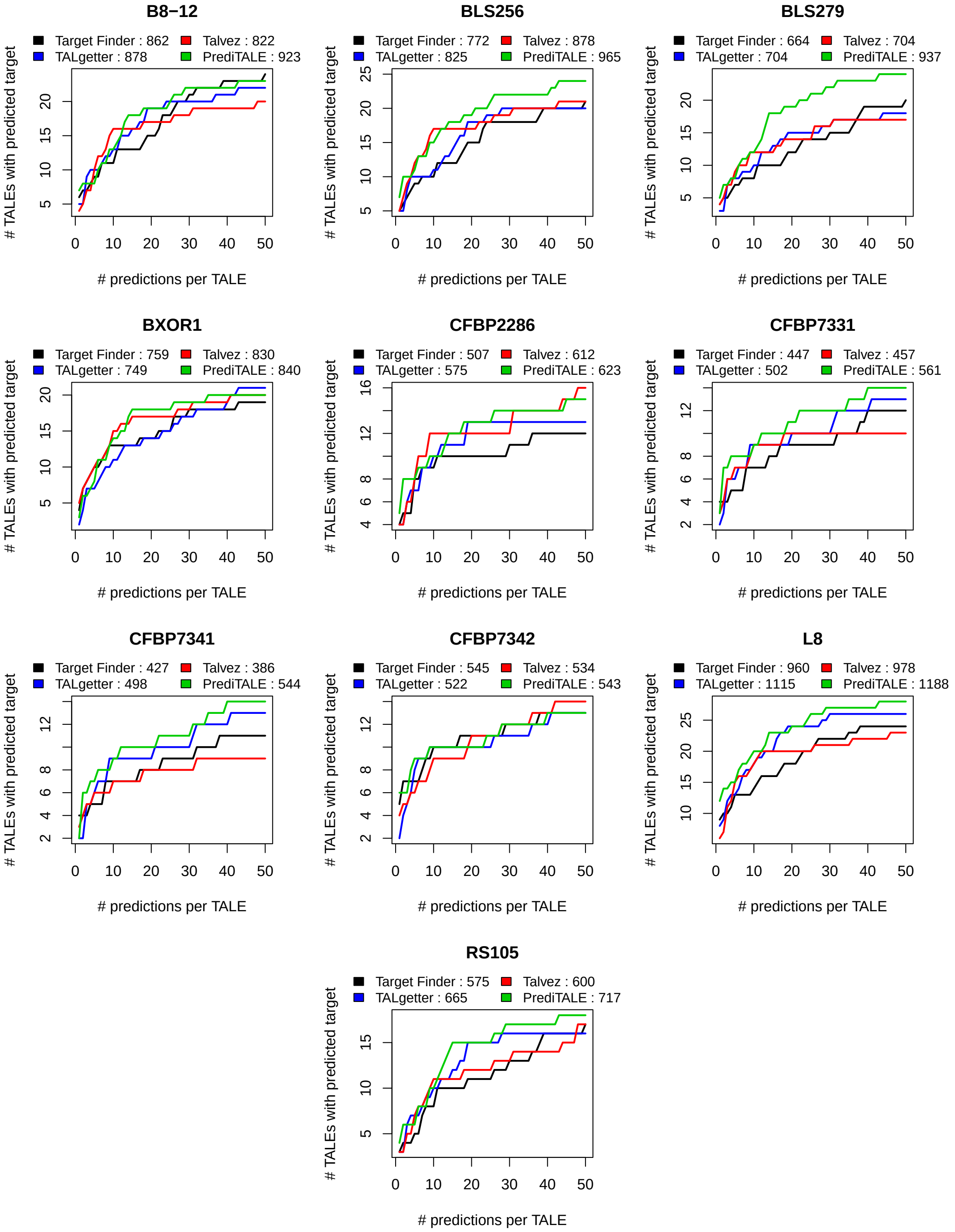


**Supplementary Figure S7.** Performance evaluation on the level of TALEs for 10 *Xoc* strains. For each approach, we plot the number of TALEs with at least one predicted target gene that is also up-regulated in the infection against the number of predicted target sites per TALE.

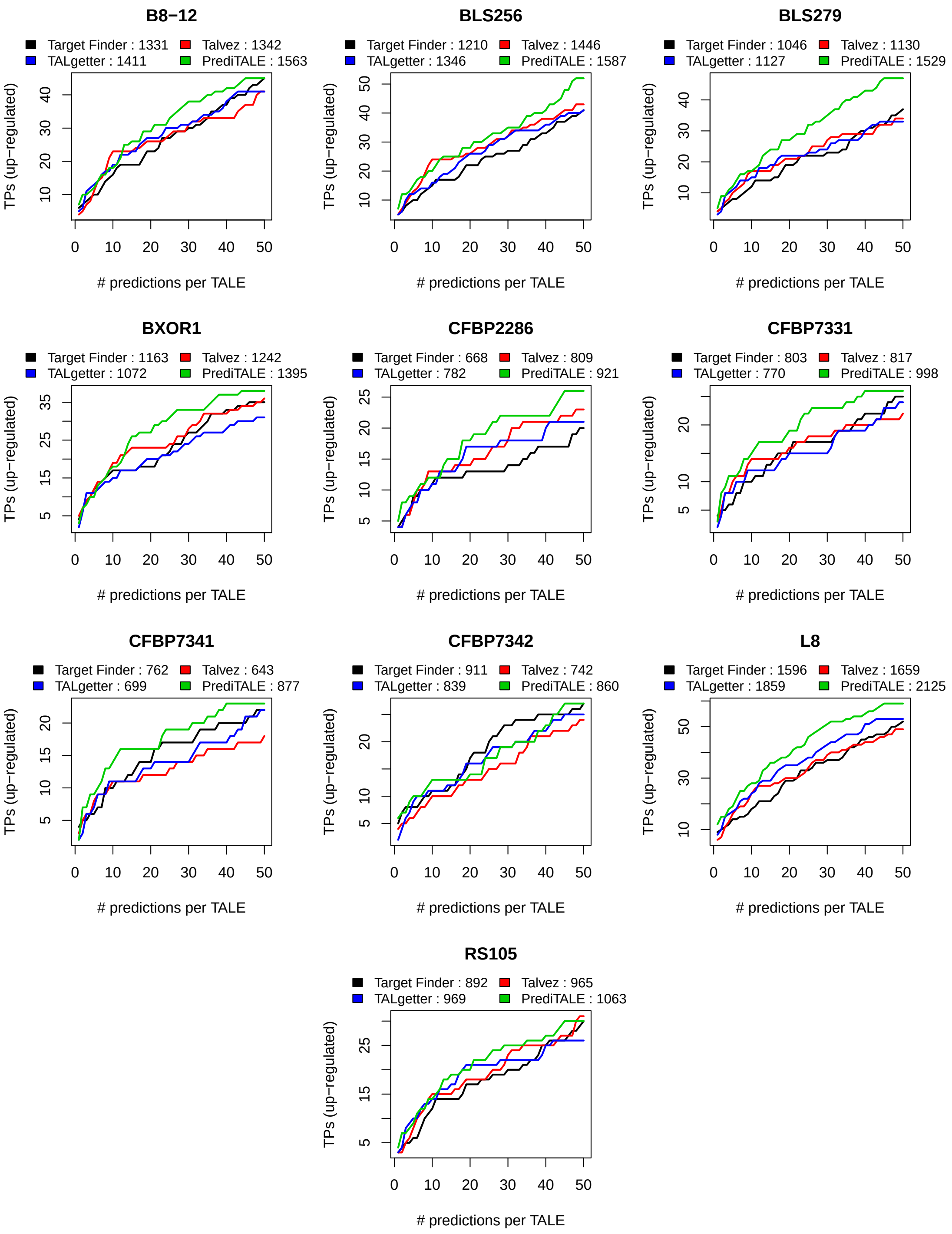


**Supplementary Figure S8.** Performance evaluation on the level of target genes for 10 *Xoc* strains. For each approach, we plot the number of predicted target genes that are also up-regulated in the infection (q-value $< 0.05$, log fold change $> 2$) against the number of predicted target sites per TALE.

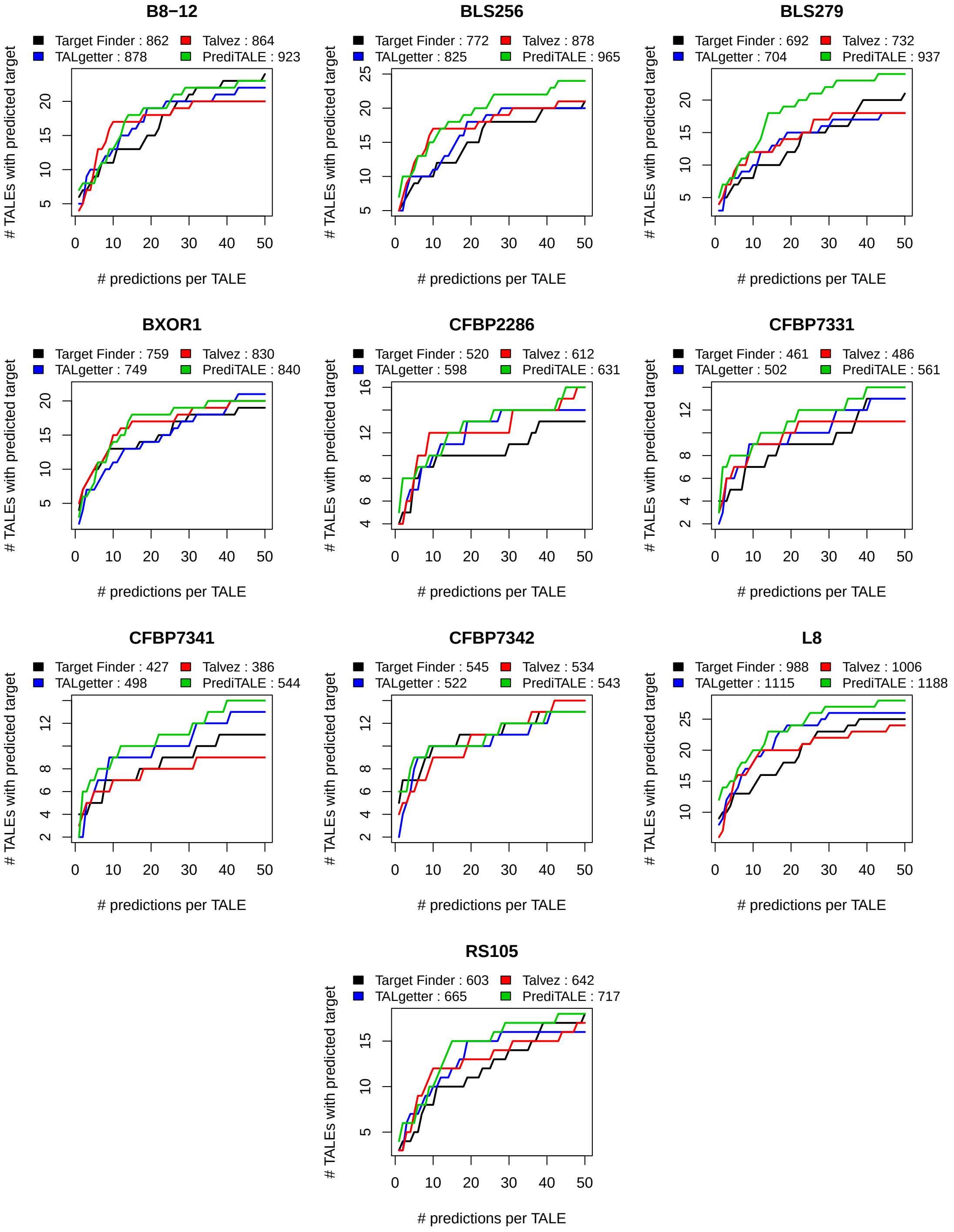


**Supplementary Figure S9.** Performance evaluation on the level of TALEs for 10 *Xoc* strains. For each approach, we plot the number of TALEs with at least one predicted target gene that is also up-regulated in the infection (q-value $< 0.05$, log fold change $> 2$) against the number of predicted target sites per TALE.

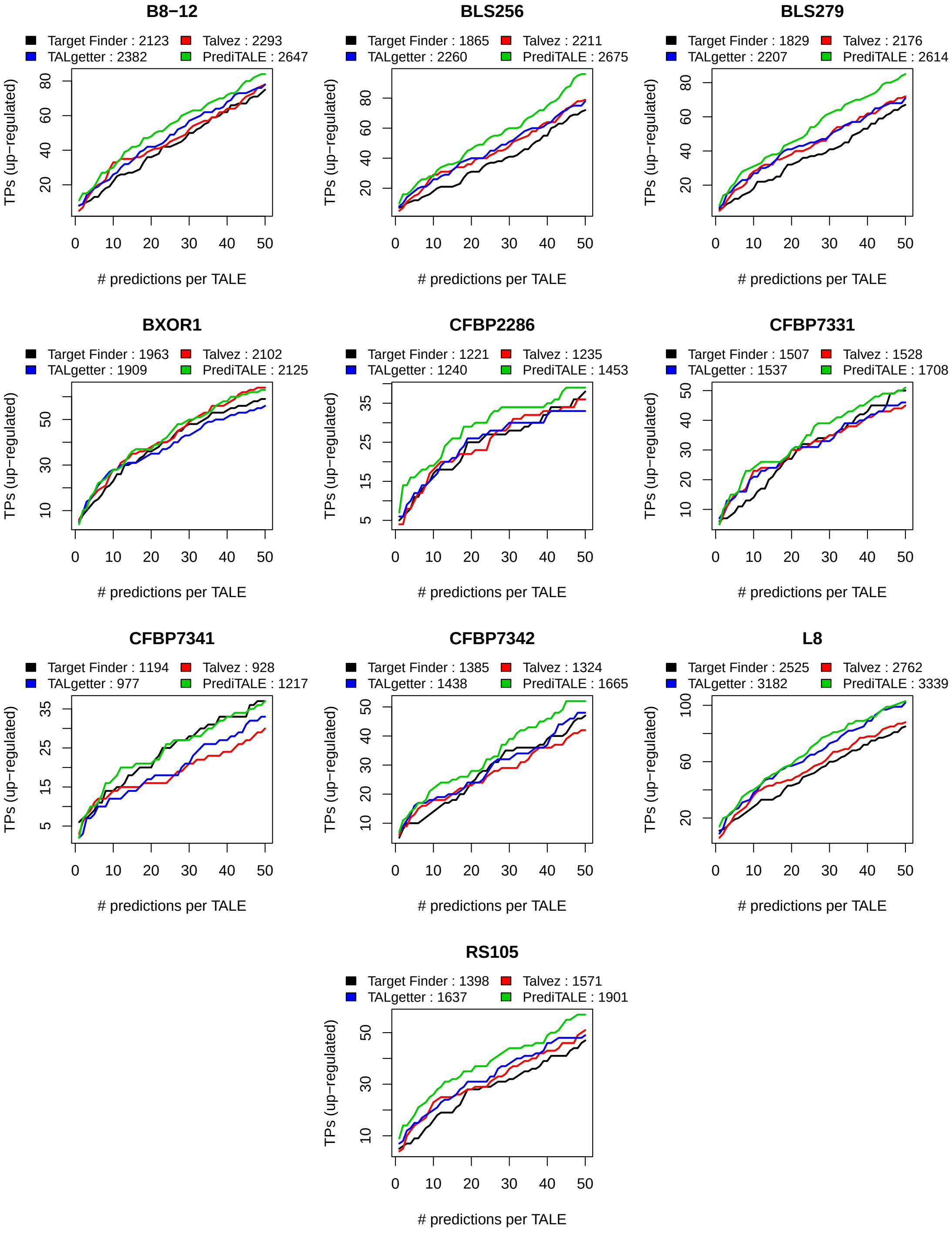


**Supplementary Figure S10.** Performance evaluation on the level of target genes for 10 *Xoc* strains. For each approach, we plot the number of predicted target genes that are also up-regulated in the infection (q-value $< 0.01$, log fold change $> 1$) against the number of predicted target sites per TALE.

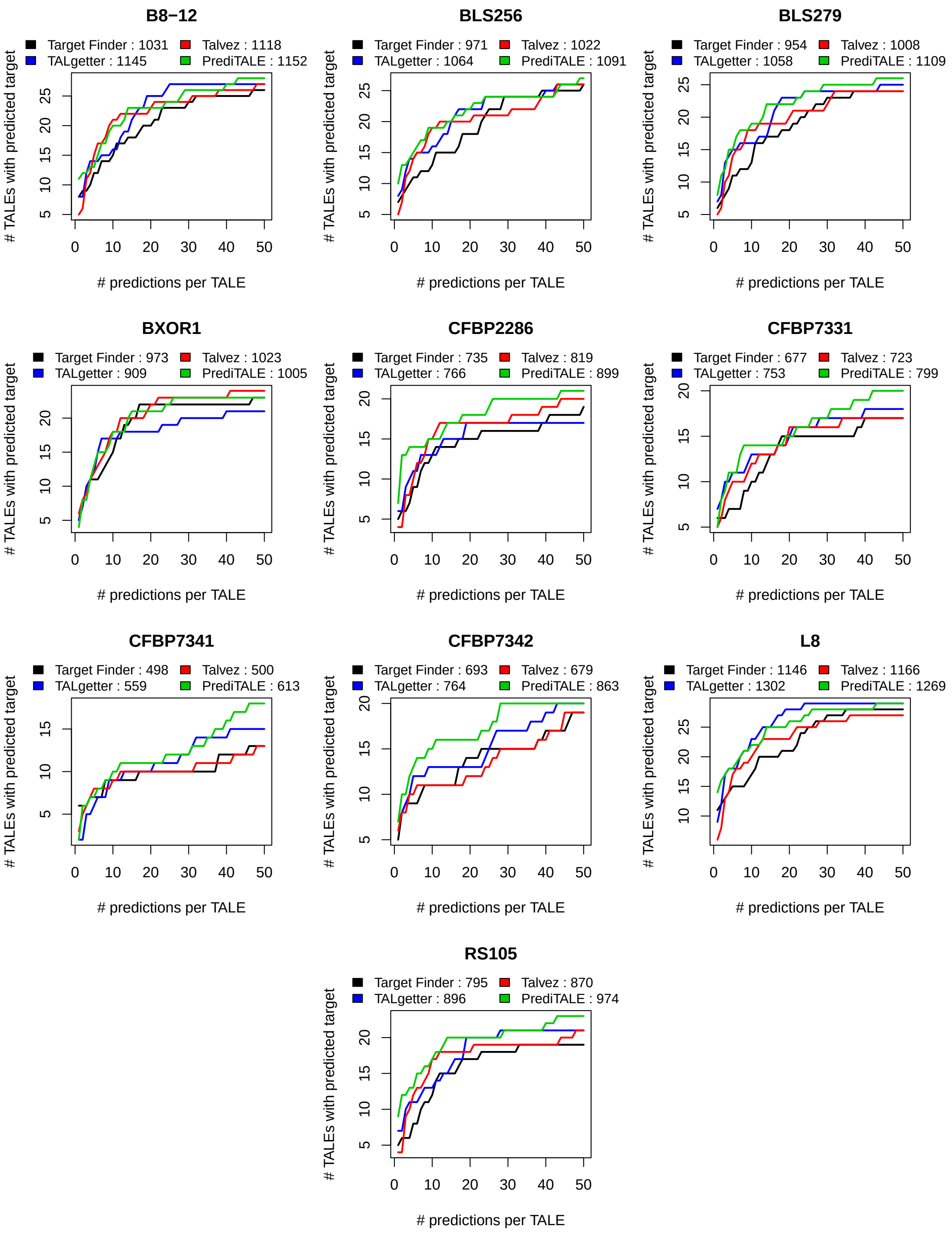


**Supplementary Figure S11.** Performance evaluation on the level of TALEs for 10 *Xoc* strains. For each approach, we plot the number of TALEs with at least one predicted target gene that is also up-regulated in the infection (q-value $< 0.01$, log fold change $> 1$) against the number of predicted target sites per TALE.

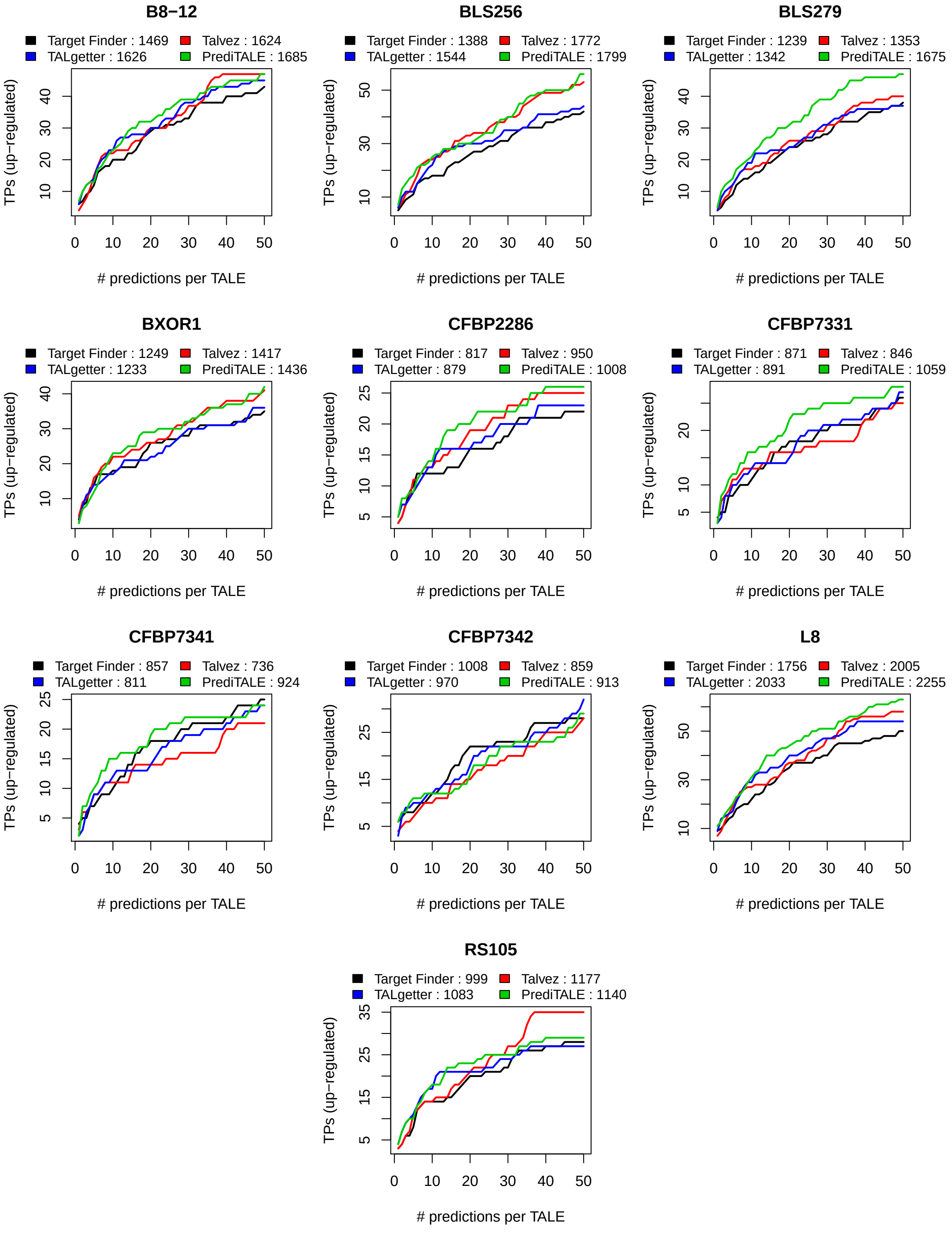


**Supplementary Figure S12.** Performance evaluation on the level of target genes for 10 *Xoc* strains when filtering for predictions of TALE boxes on the same strand as the downstream gene. For each approach, we plot the number of predicted target genes that are also up-regulated in the infection against the number of predicted target sites per TALE.

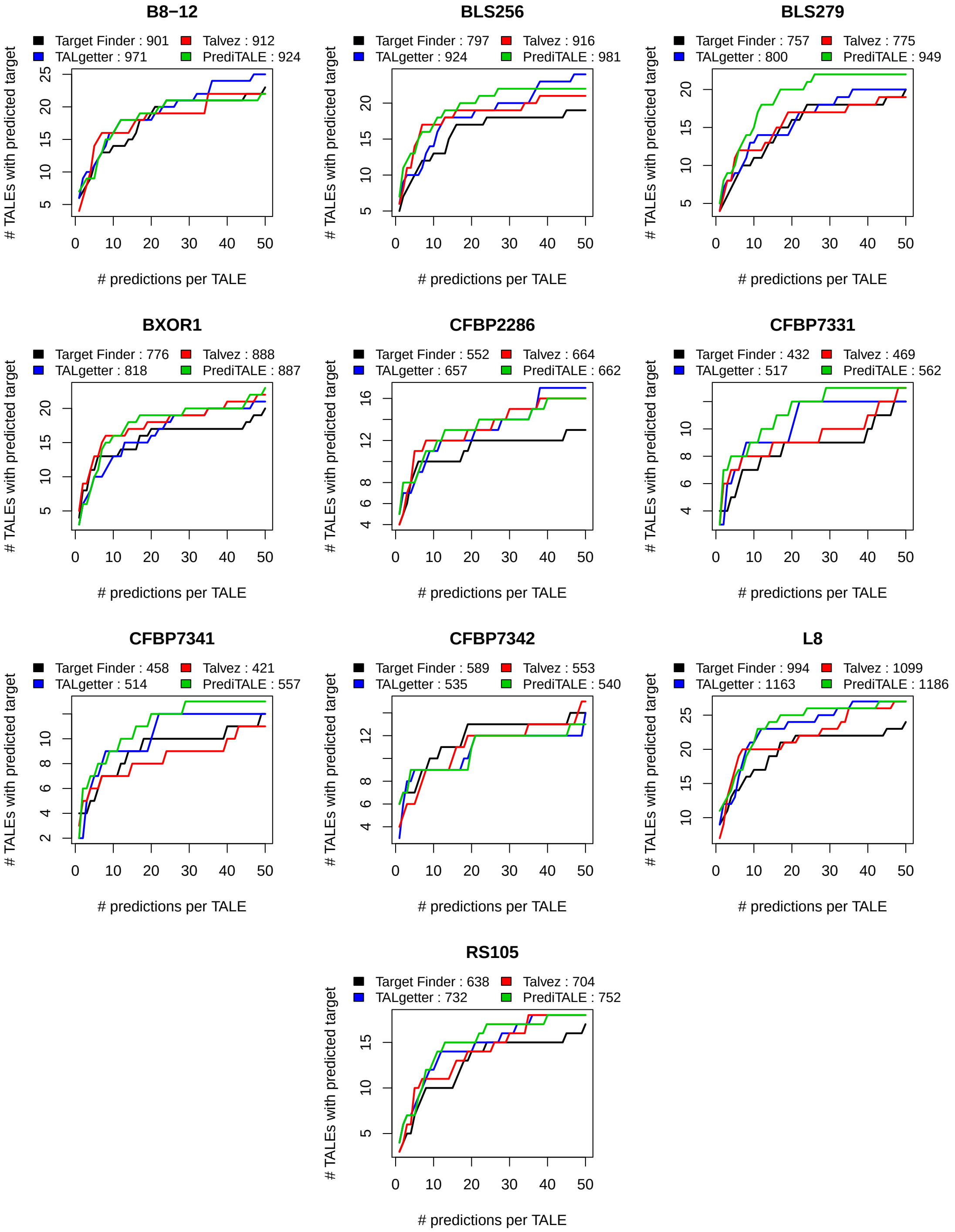


**Supplementary Figure S13.** Performance evaluation on the level of TALEs for 10 *Xoc* strains when filtering for predictions of TALE boxes on the same strand as the downstream gene. For each approach, we plot the number of TALEs with at least one predicted target gene that is also up-regulated in the infection against the number of predicted target sites per TALE.

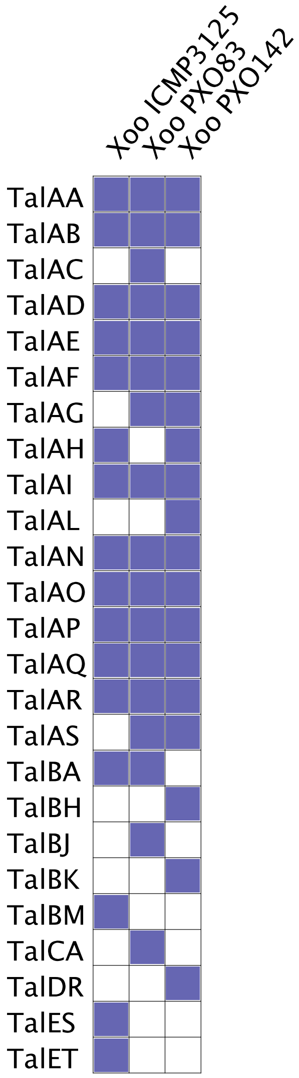


**Supplementary Figure S14.** Presence of TALE classes in the three *Xoo* strains studied according to AnnoTALE.

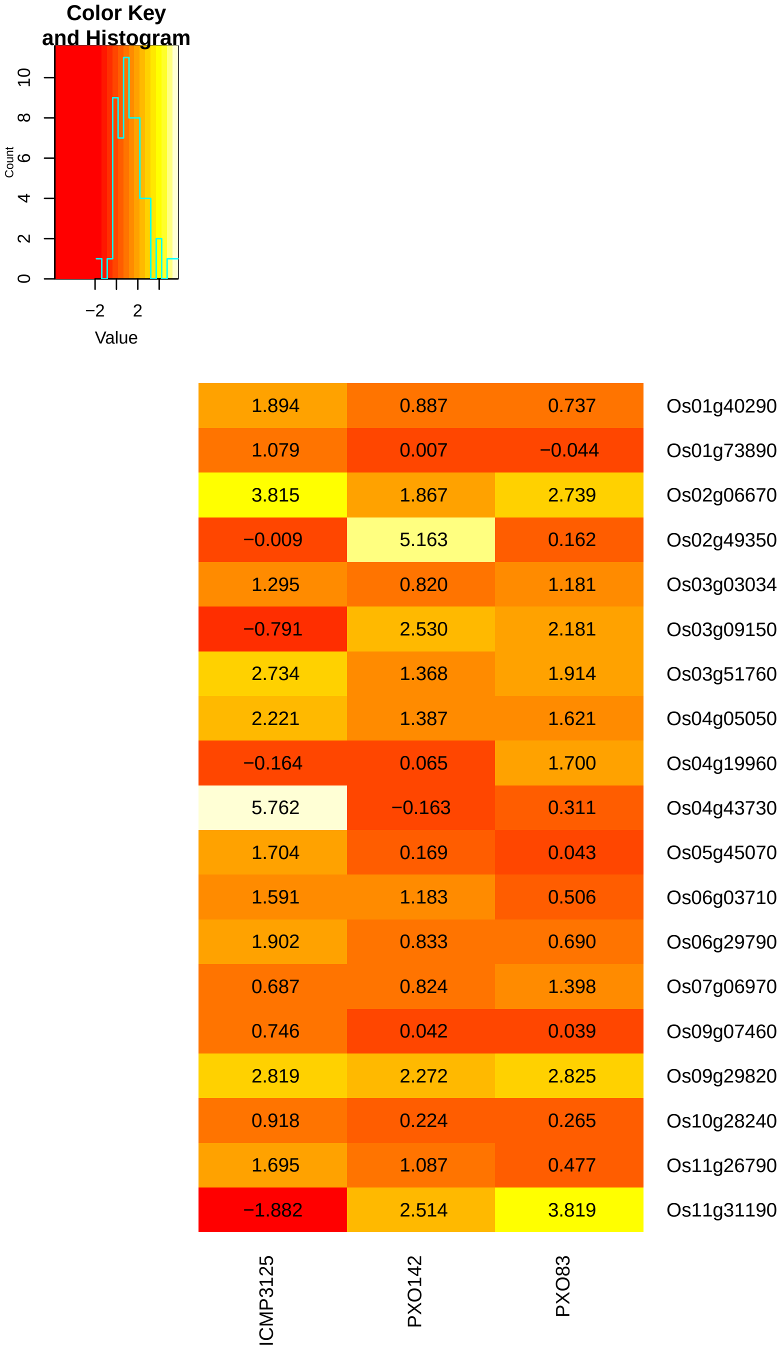


**Supplementary Figure S15.** Log fold changes of the genes that are present among the top 20 predicted target genes of any of the four approaches and that are up-regulated in at least one of the *Xoo* strains.

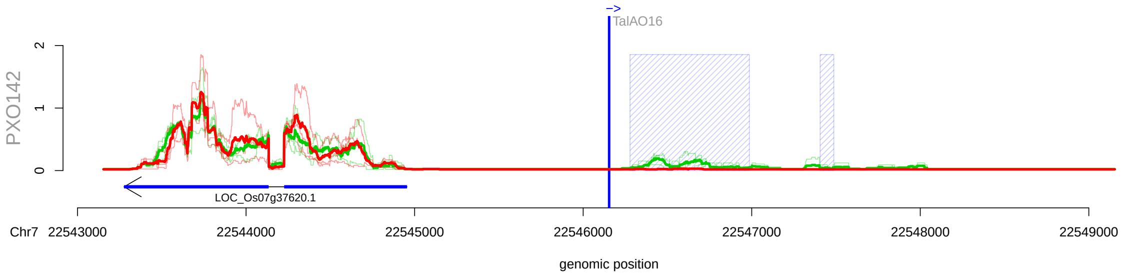


**Supplementary Figure S16.** Genome-wide prediction of TalAO16 in *Oryza sativa* Nipponbare with corresponding RNA-seq data. RNA-seq coverage after inoculation (green line) is compared with mock control (red line). In addition, we show the average of individual replicates of control and treatment are summarized as thick lines. The blue shaded boxes mark the differentially expressed regions. The arrows under the profiles reflect the MSU7 annotation within the genomic region. The genomic position of the TALE target box is marked by a vertical blue line.

### Supplementary Tables

| Strain | #DEGs ($q < 0.01$, $lfc > 2$) | #DEGs ($p < 0.05$, $lfc > \log(2)$) |
|---|---|---|
| ICMP3125 | 7 | 107 |
| PXO142 | 2 | 43 |
| PXO83 | 2 | 49 |

**Supplementary Table A.** Number of differentially expressed genes (DEGs) using the specified thresholds on p/q-values and log fold changes (lfc), respectively, considering RNA-seq data for three *Xoo* strains compared with mock inoculation.

| Strain | #DEGs |
|---|---|
| B8-12 | 628 |
| BLS256 | 652 |
| BLS279 | 567 |
| BXOR1 | 443 |
| CFBP2286 | 202 |
| CFBP7331 | 368 |
| CFBP7341 | 328 |
| CFBP7342 | 494 |
| L8 | 672 |
| RS105 | 335 |

**Supplementary Table B.** Number of differentially expressed genes (DEGs) using a threshold of 0.01 on the FDR-corrected q-values and a threshold of 2 on the log fold change considering RNA-seq data for ten *Xoc* strains compared with mock inoculation.

| strain | # predictions | Target.Finder | TALgetter | Talvez | PrediTALE |
|---|---|---|---|---|---|
| ICMP3125 | 1 | 1 (4) | 3 (2) | 2 (3) | 4 (1) |
| PXO142 | 1 | 3 (1) | 3 (1) | 1 (4) | 2 (3) |
| PXO83 | 1 | 3 (1) | 3 (1) | 1 (4) | 3 (1) |
| avg. rank | 1 | 2.00 | 1.33 | 3.67 | 1.67 |
| ICMP3125 | 10 | 6 (2) | 5 (3) | 5 (3) | 7 (1) |
| PXO142 | 10 | 4 (3) | 5 (2) | 4 (3) | 7 (1) |
| PXO83 | 10 | 3 (4) | 4 (2) | 4 (2) | 5 (1) |
| avg. rank | 10 | 3.00 | 2.33 | 2.67 | 1.00 |
| ICMP3125 | 20 | 6 (4) | 8 (2) | 8 (2) | 10 (1) |
| PXO142 | 20 | 5 (2) | 5 (2) | 5 (2) | 8 (1) |
| PXO83 | 20 | 3 (4) | 4 (2) | 4 (2) | 7 (1) |
| avg. rank | 20 | 3.33 | 2.00 | 2.00 | 1.00 |
| ICMP3125 | 50 | 10 (4) | 12 (2) | 11 (3) | 14 (1) |
| PXO142 | 50 | 5 (3) | 6 (2) | 5 (3) | 9 (1) |
| PXO83 | 50 | 5 (3) | 6 (2) | 5 (3) | 7 (1) |
| avg. rank | 50 | 3.33 | 2.00 | 3.00 | 1.00 |
| ICMP3125 | Genes AUC | 355 (4) | 393 (2) | 361 (3) | 501 (1) |
| PXO142 | Genes AUC | 232 (3) | 247 (2) | 232 (3) | 391 (1) |
| PXO83 | Genes AUC | 199 (3) | 214 (2) | 199 (3) | 320 (1) |
| avg. rank | Genes AUC | 3.33 | 2.00 | 3.00 | 1.00 |

**Supplementary Table C.** Performance evaluation on the level of target genes for three *Xoo* strains. For each strain and each approach, we list the number of predicted target genes that are also up-regulated in the infection for different thresholds on the number of predictions per TALE, i.e., prediction cutoffs. For each threshold, we additionally report the average rank of each tool.

| strain | # predictions | Target.Finder | TALgetter | Talvez | PrediTALE |
|---|---|---|---|---|---|
| ICMP3125 | 1 | 1 (4) | 3 (2) | 2 (3) | 4 (1) |
| PXO142 | 1 | 3 (1) | 3 (1) | 1 (4) | 2 (3) |
| PXO83 | 1 | 3 (1) | 3 (1) | 1 (4) | 3 (1) |
| avg. rank | 1 | 2.00 | 1.33 | 3.67 | 1.67 |
| ICMP3125 | 10 | 6 (1) | 5 (3) | 5 (3) | 6 (1) |
| PXO142 | 10 | 4 (3) | 5 (1) | 4 (3) | 5 (1) |
| PXO83 | 10 | 3 (4) | 4 (2) | 4 (2) | 5 (1) |
| avg. rank | 10 | 2.67 | 2.00 | 2.67 | 1.00 |
| ICMP3125 | 20 | 6 (4) | 7 (2) | 7 (2) | 9 (1) |
| PXO142 | 20 | 5 (2) | 5 (2) | 5 (2) | 6 (1) |
| PXO83 | 20 | 3 (4) | 4 (2) | 4 (2) | 7 (1) |
| avg. rank | 20 | 3.33 | 2.00 | 2.00 | 1.00 |
| ICMP3125 | 50 | 8 (3) | 10 (1) | 8 (3) | 10 (1) |
| PXO142 | 50 | 5 (3) | 6 (2) | 5 (3) | 7 (1) |
| PXO83 | 50 | 4 (3) | 6 (2) | 4 (3) | 7 (1) |
| avg. rank | 50 | 3.00 | 1.67 | 3.00 | 1.00 |
| ICMP3125 | TALEs AUC | 328 (3) | 355 (2) | 321 (4) | 418 (1) |
| PXO142 | TALEs AUC | 232 (3) | 247 (2) | 232 (3) | 301 (1) |
| PXO83 | TALEs AUC | 179 (4) | 214 (2) | 197 (3) | 320 (1) |
| avg. rank | TALEs AUC | 3.33 | 2.00 | 3.33 | 1.00 |

**Supplementary Table D.** Performance evaluation on the level of TALEs for three *Xoo* strains. For each strain and each approach, we list the number of TALEs with at least one predicted target gene that is also up-regulated in the infection for different thresholds on the number of predictions per TALE, i.e., prediction cutoffs. For each threshold, we additionally report the average rank of each tool.

| strain | # predictions | Target.Finder | TALgetter | Talvez | PrediTALE |
|---|---|---|---|---|---|
| ICMP3125 | 1 | 1 (1) | 1 (1) | 1 (1) | 1 (1) |
| PXO142 | 1 | 1 (1) | 0 (2) | 0 (2) | 0 (2) |
| PXO83 | 1 | 1 (1) | 1 (1) | 0 (3) | 0 (3) |
| avg. rank | 1 | 1.00 | 1.33 | 2.00 | 2.00 |
| ICMP3125 | 10 | 3 (1) | 2 (4) | 3 (1) | 3 (1) |
| PXO142 | 10 | 1 (2) | 1 (2) | 1 (2) | 2 (1) |
| PXO83 | 10 | 1 (1) | 1 (1) | 1 (1) | 1 (1) |
| avg. rank | 10 | 1.33 | 2.33 | 1.33 | 1.00 |
| ICMP3125 | 20 | 3 (2) | 3 (2) | 3 (2) | 4 (1) |
| PXO142 | 20 | 1 (2) | 1 (2) | 1 (2) | 2 (1) |
| PXO83 | 20 | 1 (1) | 1 (1) | 1 (1) | 1 (1) |
| avg. rank | 20 | 1.67 | 1.67 | 1.67 | 1.00 |
| ICMP3125 | 50 | 5 (1) | 3 (4) | 4 (2) | 4 (2) |
| PXO142 | 50 | 1 (2) | 1 (2) | 1 (2) | 2 (1) |
| PXO83 | 50 | 1 (1) | 1 (1) | 1 (1) | 1 (1) |
| avg. rank | 50 | 1.33 | 2.33 | 1.67 | 1.33 |
| ICMP3125 | Genes AUC | 191 (1) | 132 (4) | 142 (3) | 181 (2) |
| PXO142 | Genes AUC | 50 (2) | 46 (4) | 49 (3) | 91 (1) |
| PXO83 | Genes AUC | 50 (1) | 50 (1) | 49 (3) | 46 (4) |
| avg. rank | Genes AUC | 1.33 | 3.00 | 3.00 | 2.33 |

**Supplementary Table E.** Performance evaluation on the level of target genes for three *Xoo* strains. For each strain and each approach, we list the number of predicted target genes that are also up-regulated in the infection (q-value $< 0.01$, log fold change $> 2$) for different thresholds on the number of predictions per TALE, i.e., prediction cutoffs. For each threshold, we additionally report the average rank of each tool.

| strain | # predictions | Target.Finder | TALgetter | Talvez | PrediTALE |
|---|---|---|---|---|---|
| ICMP3125 | 1 | 1 (1) | 1 (1) | 1 (1) | 1 (1) |
| PXO142 | 1 | 1 (1) | 0 (2) | 0 (2) | 0 (2) |
| PXO83 | 1 | 1 (1) | 1 (1) | 0 (3) | 0 (3) |
| avg. rank | 1 | 1.00 | 1.33 | 2.00 | 2.00 |
| ICMP3125 | 10 | 3 (1) | 2 (4) | 3 (1) | 3 (1) |
| PXO142 | 10 | 1 (1) | 1 (1) | 1 (1) | 1 (1) |
| PXO83 | 10 | 1 (1) | 1 (1) | 1 (1) | 1 (1) |
| avg. rank | 10 | 1.00 | 2.00 | 1.00 | 1.00 |
| ICMP3125 | 20 | 3 (2) | 3 (2) | 3 (2) | 4 (1) |
| PXO142 | 20 | 1 (1) | 1 (1) | 1 (1) | 1 (1) |
| PXO83 | 20 | 1 (1) | 1 (1) | 1 (1) | 1 (1) |
| avg. rank | 20 | 1.33 | 1.33 | 1.33 | 1.00 |
| ICMP3125 | 50 | 4 (1) | 3 (3) | 3 (3) | 4 (1) |
| PXO142 | 50 | 1 (1) | 1 (1) | 1 (1) | 1 (1) |
| PXO83 | 50 | 1 (1) | 1 (1) | 1 (1) | 1 (1) |
| avg. rank | 50 | 1.00 | 1.67 | 1.67 | 1.00 |
| ICMP3125 | TALEs AUC | 171 (2) | 132 (4) | 140 (3) | 181 (1) |
| PXO142 | TALEs AUC | 50 (1) | 46 (4) | 49 (2) | 48 (3) |
| PXO83 | TALEs AUC | 50 (1) | 50 (1) | 49 (3) | 46 (4) |
| avg. rank | TALEs AUC | 1.33 | 3.00 | 2.67 | 2.67 |

**Supplementary Table F.** Performance evaluation on the level of TALEs for three *Xoo* strains. For each strain and each approach, we list the number of TALEs with at least one predicted target gene that is also up-regulated in the infection (q-value $< 0.01$, log fold change $> 2$) for different thresholds on the number of predictions per TALE, i.e., prediction cutoffs. For each threshold, we additionally report the average rank of each tool.

| strain | # predictions | Target.Finder | TALgetter | Talvez | PrediTALE |
|---|---|---|---|---|---|
| ICMP3125 | 1 | 1 (4) | 3 (2) | 3 (2) | 4 (1) |
| PXO142 | 1 | 3 (1) | 3 (1) | 1 (4) | 2 (3) |
| PXO83 | 1 | 3 (1) | 3 (1) | 2 (4) | 3 (1) |
| avg. rank | 1 | 2.00 | 1.33 | 3.33 | 1.67 |
| ICMP3125 | 10 | 6 (3) | 6 (3) | 8 (2) | 9 (1) |
| PXO142 | 10 | 5 (2) | 5 (2) | 5 (2) | 8 (1) |
| PXO83 | 10 | 3 (4) | 4 (2) | 4 (2) | 7 (1) |
| avg. rank | 10 | 3.00 | 2.33 | 2.00 | 1.00 |
| ICMP3125 | 20 | 8 (2) | 8 (2) | 8 (2) | 11 (1) |
| PXO142 | 20 | 5 (2) | 5 (2) | 5 (2) | 8 (1) |
| PXO83 | 20 | 5 (2) | 5 (2) | 4 (4) | 7 (1) |
| avg. rank | 20 | 2.00 | 2.00 | 2.67 | 1.00 |
| ICMP3125 | 50 | 10 (3) | 15 (1) | 10 (3) | 14 (2) |
| PXO142 | 50 | 5 (3) | 6 (2) | 5 (3) | 9 (1) |
| PXO83 | 50 | 6 (3) | 10 (1) | 6 (3) | 7 (2) |
| avg. rank | 50 | 3.00 | 1.33 | 3.00 | 1.67 |
| ICMP3125 | Genes AUC | 387 (4) | 490 (2) | 412 (3) | 563 (1) |
| PXO142 | Genes AUC | 242 (3) | 266 (2) | 240 (4) | 402 (1) |
| PXO83 | Genes AUC | 226 (4) | 302 (2) | 241 (3) | 328 (1) |
| avg. rank | Genes AUC | 3.67 | 2.00 | 3.33 | 1.00 |

**Supplementary Table G.** Performance evaluation on the level of target genes for three *Xoo* strains when filtering for predictions of TALE boxes on the same strand as the downstream gene. For each strain and each approach, we list the number of predicted target genes that are also up-regulated in the infection for different thresholds on the number of predictions per TALE, i.e., prediction cutoffs. For each threshold, we additionally report the average rank of each tool.

| strain | # predictions | Target.Finder | TALgetter | Talvez | PrediTALE |
|---|---|---|---|---|---|
| ICMP3125 | 1 | 1 (4) | 3 (2) | 3 (2) | 4 (1) |
| PXO142 | 1 | 3 (1) | 3 (1) | 1 (4) | 2 (3) |
| PXO83 | 1 | 3 (1) | 3 (1) | 2 (4) | 3 (1) |
| avg. rank | 1 | 2.00 | 1.33 | 3.33 | 1.67 |
| ICMP3125 | 10 | 6 (3) | 6 (3) | 7 (2) | 8 (1) |
| PXO142 | 10 | 5 (2) | 5 (2) | 5 (2) | 6 (1) |
| PXO83 | 10 | 3 (4) | 4 (2) | 4 (2) | 7 (1) |
| avg. rank | 10 | 3.00 | 2.33 | 2.00 | 1.00 |
| ICMP3125 | 20 | 7 (2) | 7 (2) | 7 (2) | 9 (1) |
| PXO142 | 20 | 5 (2) | 5 (2) | 5 (2) | 6 (1) |
| PXO83 | 20 | 4 (3) | 5 (2) | 4 (3) | 7 (1) |
| avg. rank | 20 | 2.33 | 2.00 | 2.33 | 1.00 |
| ICMP3125 | 50 | 7 (3) | 10 (1) | 7 (3) | 10 (1) |
| PXO142 | 50 | 5 (3) | 6 (2) | 5 (3) | 7 (1) |
| PXO83 | 50 | 5 (3) | 8 (1) | 5 (3) | 7 (2) |
| avg. rank | 50 | 3.00 | 1.33 | 3.00 | 1.33 |
| ICMP3125 | TALEs AUC | 325 (3) | 389 (2) | 323 (4) | 440 (1) |
| PXO142 | TALEs AUC | 242 (3) | 266 (2) | 240 (4) | 312 (1) |
| PXO83 | TALEs AUC | 190 (4) | 271 (2) | 217 (3) | 328 (1) |
| avg. rank | TALEs AUC | 3.33 | 2.00 | 3.67 | 1.00 |

**Supplementary Table H.** Performance evaluation on the level of TALEs for three *Xoo* strains when filtering for predictions of TALE boxes on the same strand as the downstream gene. For each strain and each approach, we list the number of TALEs with at least one predicted target gene that is also up-regulated in the infection for different thresholds on the number of predictions per TALE, i.e., prediction cutoffs. For each threshold, we additionally report the average rank of each tool.

| strain | # predictions | Target.Finder | TALgetter | Talvez | PrediTALE |
|---|---|---|---|---|---|
| B8-12 | 1 | 6 (2) | 5 (3) | 4 (4) | 7 (1) |
| BLS256 | 1 | 5 (2) | 5 (2) | 5 (2) | 7 (1) |
| BLS279 | 1 | 4 (2) | 3 (4) | 4 (2) | 5 (1) |
| BXOR1 | 1 | 4 (2) | 2 (4) | 5 (1) | 3 (3) |
| CFBP2286 | 1 | 4 (2) | 4 (2) | 4 (2) | 5 (1) |
| CFBP7331 | 1 | 4 (1) | 2 (4) | 3 (2) | 3 (2) |
| CFBP7341 | 1 | 4 (1) | 2 (3) | 3 (2) | 2 (3) |
| CFBP7342 | 1 | 5 (2) | 2 (4) | 4 (3) | 6 (1) |
| L8 | 1 | 9 (2) | 8 (3) | 6 (4) | 12 (1) |
| RS105 | 1 | 3 (2) | 3 (2) | 3 (2) | 4 (1) |
| avg. rank | 1 | 1.8 | 3.1 | 2.4 | 1.5 |
| B8-12 | 10 | 16 (4) | 19 (2) | 22 (1) | 18 (3) |
| BLS256 | 10 | 15 (4) | 16 (3) | 24 (1) | 20 (2) |
| BLS279 | 10 | 12 (4) | 15 (3) | 17 (1) | 17 (1) |
| BXOR1 | 10 | 17 (3) | 15 (4) | 19 (1) | 18 (2) |
| CFBP2286 | 10 | 11 (3) | 11 (3) | 13 (1) | 12 (2) |
| CFBP7331 | 10 | 10 (4) | 12 (3) | 14 (2) | 15 (1) |
| CFBP7341 | 10 | 10 (4) | 11 (2) | 11 (2) | 14 (1) |
| CFBP7342 | 10 | 11 (2) | 11 (2) | 10 (4) | 13 (1) |
| L8 | 10 | 18 (4) | 24 (2) | 24 (2) | 28 (1) |
| RS105 | 10 | 12 (4) | 14 (1) | 14 (1) | 14 (1) |
| avg. rank | 10 | 3.6 | 2.5 | 1.6 | 1.5 |
| B8-12 | 20 | 23 (4) | 27 (2) | 25 (3) | 29 (1) |
| BLS256 | 20 | 22 (4) | 26 (2) | 26 (2) | 28 (1) |
| BLS279 | 20 | 19 (4) | 22 (2) | 21 (3) | 27 (1) |
| BXOR1 | 20 | 18 (4) | 20 (3) | 23 (2) | 26 (1) |
| CFBP2286 | 20 | 13 (4) | 17 (2) | 14 (3) | 18 (1) |
| CFBP7331 | 20 | 15 (3) | 15 (3) | 16 (2) | 19 (1) |
| CFBP7341 | 20 | 14 (2) | 13 (3) | 12 (4) | 16 (1) |
| CFBP7342 | 20 | 16 (1) | 16 (1) | 13 (4) | 14 (3) |
| L8 | 20 | 29 (4) | 35 (2) | 30 (3) | 39 (1) |
| RS105 | 20 | 17 (3) | 21 (1) | 17 (3) | 20 (2) |
| avg. rank | 20 | 3.3 | 2.1 | 2.9 | 1.3 |
| B8-12 | 50 | 45 (1) | 41 (3) | 40 (4) | 45 (1) |
| BLS256 | 50 | 41 (3) | 40 (4) | 43 (2) | 52 (1) |
| BLS279 | 50 | 36 (2) | 33 (3) | 33 (3) | 47 (1) |
| BXOR1 | 50 | 33 (2) | 31 (4) | 33 (2) | 36 (1) |
| CFBP2286 | 50 | 19 (4) | 20 (3) | 23 (2) | 25 (1) |
| CFBP7331 | 50 | 24 (2) | 24 (2) | 21 (4) | 26 (1) |
| CFBP7341 | 50 | 22 (2) | 22 (2) | 18 (4) | 23 (1) |
| CFBP7342 | 50 | 26 (2) | 25 (3) | 24 (4) | 27 (1) |
| L8 | 50 | 51 (3) | 53 (2) | 48 (4) | 59 (1) |
| RS105 | 50 | 29 (2) | 26 (4) | 30 (1) | 29 (2) |
| avg. rank | 50 | 2.3 | 3.0 | 3.0 | 1.1 |
| B8-12 | Genes AUC | 1331 (3) | 1411 (2) | 1297 (4) | 1563 (1) |
| BLS256 | Genes AUC | 1210 (4) | 1345 (3) | 1446 (2) | 1587 (1) |
| BLS279 | Genes AUC | 1018 (4) | 1127 (2) | 1102 (3) | 1529 (1) |
| BXOR1 | Genes AUC | 1129 (3) | 1072 (4) | 1203 (2) | 1331 (1) |
| CFBP2286 | Genes AUC | 655 (4) | 759 (3) | 809 (2) | 913 (1) |
| CFBP7331 | Genes AUC | 789 (2) | 770 (4) | 788 (3) | 998 (1) |
| CFBP7341 | Genes AUC | 762 (2) | 699 (3) | 643 (4) | 877 (1) |
| CFBP7342 | Genes AUC | 877 (1) | 839 (3) | 742 (4) | 860 (2) |
| L8 | Genes AUC | 1568 (4) | 1859 (2) | 1631 (3) | 2125 (1) |
| RS105 | Genes AUC | 864 (4) | 969 (2) | 920 (3) | 1047 (1) |
| avg. rank | Genes AUC | 3.1 | 2.8 | 3.0 | 1.1 |

**Supplementary Table I.** Performance evaluation on the level of target genes for ten *Xoc* strains. For each strain and each approach, we list the number of predicted target genes that are also up-regulated in the infection for different cutoffs on the number of predictions per TALE, i.e., prediction ranks. For each threshold, we additionally report the average rank of each tool.

| strain | # predictions | Target.Finder | TALgetter | Talvez | PrediTALE |
|---|---|---|---|---|---|
| B8-12 | 1 | 6 (2) | 5 (3) | 4 (4) | 7 (1) |
| BLS256 | 1 | 5 (2) | 5 (2) | 5 (2) | 7 (1) |
| BLS279 | 1 | 4 (2) | 3 (4) | 4 (2) | 5 (1) |
| BXOR1 | 1 | 4 (2) | 2 (4) | 5 (1) | 3 (3) |
| CFBP2286 | 1 | 4 (2) | 4 (2) | 4 (2) | 5 (1) |
| CFBP7331 | 1 | 4 (1) | 2 (4) | 3 (2) | 3 (2) |
| CFBP7341 | 1 | 4 (1) | 2 (3) | 3 (2) | 2 (3) |
| CFBP7342 | 1 | 5 (2) | 2 (4) | 4 (3) | 6 (1) |
| L8 | 1 | 9 (2) | 8 (3) | 6 (4) | 12 (1) |
| RS105 | 1 | 3 (2) | 3 (2) | 3 (2) | 4 (1) |
| avg. rank | 1 | 1.8 | 3.1 | 2.4 | 1.5 |
| B8-12 | 10 | 11 (4) | 13 (2) | 16 (1) | 13 (2) |
| BLS256 | 10 | 10 (4) | 11 (3) | 17 (1) | 15 (2) |
| BLS279 | 10 | 8 (4) | 10 (3) | 12 (1) | 12 (1) |
| BXOR1 | 10 | 13 (3) | 11 (4) | 15 (1) | 14 (2) |
| CFBP2286 | 10 | 9 (4) | 10 (2) | 12 (1) | 10 (2) |
| CFBP7331 | 10 | 7 (4) | 9 (1) | 9 (1) | 9 (1) |
| CFBP7341 | 10 | 7 (3) | 9 (1) | 7 (3) | 9 (1) |
| CFBP7342 | 10 | 10 (1) | 10 (1) | 9 (4) | 10 (1) |
| L8 | 10 | 14 (4) | 18 (2) | 18 (2) | 20 (1) |
| RS105 | 10 | 8 (4) | 10 (2) | 11 (1) | 10 (2) |
| avg. rank | 10 | 3.5 | 2.1 | 1.6 | 1.5 |
| B8-12 | 20 | 15 (4) | 19 (1) | 17 (3) | 19 (1) |
| BLS256 | 20 | 15 (4) | 18 (2) | 17 (3) | 19 (1) |
| BLS279 | 20 | 12 (4) | 15 (2) | 14 (3) | 19 (1) |
| BXOR1 | 20 | 14 (3) | 14 (3) | 17 (2) | 18 (1) |
| CFBP2286 | 20 | 10 (4) | 13 (1) | 12 (3) | 13 (1) |
| CFBP7331 | 20 | 9 (4) | 10 (2) | 10 (2) | 11 (1) |
| CFBP7341 | 20 | 8 (3) | 9 (2) | 8 (3) | 10 (1) |
| CFBP7342 | 20 | 11 (1) | 10 (3) | 11 (1) | 10 (3) |
| L8 | 20 | 18 (4) | 24 (1) | 20 (3) | 24 (1) |
| RS105 | 20 | 11 (4) | 15 (1) | 12 (3) | 15 (1) |
| avg. rank | 20 | 3.5 | 1.8 | 2.6 | 1.2 |
| B8-12 | 50 | 24 (1) | 22 (3) | 20 (4) | 23 (2) |
| BLS256 | 50 | 21 (2) | 20 (4) | 21 (2) | 24 (1) |
| BLS279 | 50 | 20 (2) | 18 (3) | 17 (4) | 24 (1) |
| BXOR1 | 50 | 19 (4) | 21 (1) | 20 (2) | 20 (2) |
| CFBP2286 | 50 | 12 (4) | 13 (3) | 16 (1) | 15 (2) |
| CFBP7331 | 50 | 12 (3) | 13 (2) | 10 (4) | 14 (1) |
| CFBP7341 | 50 | 11 (3) | 13 (2) | 9 (4) | 14 (1) |
| CFBP7342 | 50 | 13 (2) | 13 (2) | 14 (1) | 13 (2) |
| L8 | 50 | 24 (3) | 26 (2) | 23 (4) | 28 (1) |
| RS105 | 50 | 17 (2) | 16 (4) | 17 (2) | 18 (1) |
| avg. rank | 50 | 2.6 | 2.6 | 2.8 | 1.4 |
| B8-12 | TALEs AUC | 862 (3) | 878 (2) | 822 (4) | 923 (1) |
| BLS256 | TALEs AUC | 772 (4) | 825 (3) | 878 (2) | 965 (1) |
| BLS279 | TALEs AUC | 664 (4) | 704 (2) | 704 (2) | 937 (1) |
| BXOR1 | TALEs AUC | 759 (3) | 749 (4) | 830 (2) | 840 (1) |
| CFBP2286 | TALEs AUC | 507 (4) | 575 (3) | 612 (2) | 623 (1) |
| CFBP7331 | TALEs AUC | 447 (4) | 502 (2) | 457 (3) | 561 (1) |
| CFBP7341 | TALEs AUC | 427 (3) | 498 (2) | 386 (4) | 544 (1) |
| CFBP7342 | TALEs AUC | 545 (1) | 522 (4) | 534 (3) | 543 (2) |
| L8 | TALEs AUC | 960 (4) | 1115 (2) | 978 (3) | 1188 (1) |
| RS105 | TALEs AUC | 575 (4) | 665 (2) | 600 (3) | 717 (1) |
| avg. rank | TALEs AUC | 3.4 | 2.6 | 2.8 | 1.1 |

**Supplementary Table J.** Performance evaluation on the level of TALEs for ten *Xoo* strains. For each strain and each approach, we list the number of TALEs with at least one predicted target gene that is also up-regulated in the infection for different cutoffs on the number of predictions per TALE, i.e., prediction ranks. For each threshold, we additionally report the average rank of each tool.

| strain | # predictions | Target.Finder | TALgetter | Talvez | PrediTALE |
|---|---|---|---|---|---|
| B8-12 | 1 | 6 (2) | 5 (3) | 4 (4) | 7 (1) |
| BLS256 | 1 | 5 (2) | 5 (2) | 5 (2) | 7 (1) |
| BLS279 | 1 | 4 (2) | 3 (4) | 4 (2) | 5 (1) |
| BXOR1 | 1 | 4 (2) | 2 (4) | 5 (1) | 3 (3) |
| CFBP2286 | 1 | 4 (2) | 4 (2) | 4 (2) | 5 (1) |
| CFBP7331 | 1 | 4 (1) | 2 (4) | 3 (2) | 3 (2) |
| CFBP7341 | 1 | 4 (1) | 2 (3) | 3 (2) | 2 (3) |
| CFBP7342 | 1 | 5 (2) | 2 (4) | 4 (3) | 6 (1) |
| L8 | 1 | 9 (2) | 8 (3) | 6 (4) | 12 (1) |
| RS105 | 1 | 3 (2) | 3 (2) | 3 (2) | 4 (1) |
| avg. rank | 1 | 1.8 | 3.1 | 2.4 | 1.5 |
| B8-12 | 10 | 16 (4) | 19 (2) | 23 (1) | 18 (3) |
| BLS256 | 10 | 15 (4) | 16 (3) | 24 (1) | 20 (2) |
| BLS279 | 10 | 12 (4) | 15 (3) | 17 (1) | 17 (1) |
| BXOR1 | 10 | 17 (3) | 15 (4) | 19 (1) | 18 (2) |
| CFBP2286 | 10 | 11 (3) | 11 (3) | 13 (1) | 12 (2) |
| CFBP7331 | 10 | 10 (4) | 12 (3) | 14 (2) | 15 (1) |
| CFBP7341 | 10 | 10 (4) | 11 (2) | 11 (2) | 14 (1) |
| CFBP7342 | 10 | 11 (2) | 11 (2) | 10 (4) | 13 (1) |
| L8 | 10 | 18 (4) | 24 (2) | 24 (2) | 28 (1) |
| RS105 | 10 | 12 (4) | 14 (2) | 15 (1) | 14 (2) |
| avg. rank | 10 | 3.6 | 2.6 | 1.6 | 1.6 |
| B8-12 | 20 | 23 (4) | 27 (2) | 26 (3) | 29 (1) |
| BLS256 | 20 | 22 (4) | 26 (2) | 26 (2) | 28 (1) |
| BLS279 | 20 | 19 (4) | 22 (2) | 21 (3) | 27 (1) |
| BXOR1 | 20 | 18 (4) | 20 (3) | 23 (2) | 27 (1) |
| CFBP2286 | 20 | 13 (4) | 17 (2) | 14 (3) | 18 (1) |
| CFBP7331 | 20 | 15 (3) | 15 (3) | 16 (2) | 19 (1) |
| CFBP7341 | 20 | 14 (2) | 13 (3) | 12 (4) | 16 (1) |
| CFBP7342 | 20 | 17 (1) | 16 (2) | 13 (4) | 14 (3) |
| L8 | 20 | 29 (4) | 35 (2) | 30 (3) | 39 (1) |
| RS105 | 20 | 17 (4) | 21 (1) | 18 (3) | 20 (2) |
| avg. rank | 20 | 3.4 | 2.2 | 2.9 | 1.3 |
| B8-12 | 50 | 45 (1) | 41 (3) | 41 (3) | 45 (1) |
| BLS256 | 50 | 41 (3) | 41 (3) | 43 (2) | 52 (1) |
| BLS279 | 50 | 37 (2) | 33 (4) | 34 (3) | 47 (1) |
| BXOR1 | 50 | 35 (3) | 31 (4) | 36 (2) | 38 (1) |
| CFBP2286 | 50 | 20 (4) | 21 (3) | 23 (2) | 26 (1) |
| CFBP7331 | 50 | 25 (2) | 24 (3) | 22 (4) | 26 (1) |
| CFBP7341 | 50 | 22 (2) | 22 (2) | 18 (4) | 23 (1) |
| CFBP7342 | 50 | 27 (1) | 25 (3) | 24 (4) | 27 (1) |
| L8 | 50 | 52 (3) | 53 (2) | 49 (4) | 59 (1) |
| RS105 | 50 | 30 (2) | 26 (4) | 31 (1) | 30 (2) |
| avg. rank | 50 | 2.3 | 3.1 | 2.9 | 1.1 |
| B8-12 | Genes AUC | 1331 (4) | 1411 (2) | 1342 (3) | 1563 (1) |
| BLS256 | Genes AUC | 1210 (4) | 1346 (3) | 1446 (2) | 1587 (1) |
| BLS279 | Genes AUC | 1046 (4) | 1127 (3) | 1130 (2) | 1529 (1) |
| BXOR1 | Genes AUC | 1163 (3) | 1072 (4) | 1242 (2) | 1395 (1) |
| CFBP2286 | Genes AUC | 668 (4) | 782 (3) | 809 (2) | 921 (1) |
| CFBP7331 | Genes AUC | 803 (3) | 770 (4) | 817 (2) | 998 (1) |
| CFBP7341 | Genes AUC | 762 (2) | 699 (3) | 643 (4) | 877 (1) |
| CFBP7342 | Genes AUC | 911 (1) | 839 (3) | 742 (4) | 860 (2) |
| L8 | Genes AUC | 1596 (4) | 1859 (2) | 1659 (3) | 2125 (1) |
| RS105 | Genes AUC | 892 (4) | 969 (2) | 965 (3) | 1063 (1) |
| avg. rank | Genes AUC | 3.3 | 2.9 | 2.7 | 1.1 |

**Supplementary Table K.** Performance evaluation on the level of target genes for ten *Xoc* strains. For each strain and each approach, we list the number of predicted target genes that are also up-regulated in the infection (q-value $< 0.05$, log fold change $> 2$) for different cutoffs on the number of predictions per TALE, i.e., prediction ranks. For each threshold, we additionally report the average rank of each tool.

| strain | # predictions | Target.Finder | TALgetter | Talvez | PrediTALE |
|---|---|---|---|---|---|
| B8-12 | 1 | 6 (2) | 5 (3) | 4 (4) | 7 (1) |
| BLS256 | 1 | 5 (2) | 5 (2) | 5 (2) | 7 (1) |
| BLS279 | 1 | 4 (2) | 3 (4) | 4 (2) | 5 (1) |
| BXOR1 | 1 | 4 (2) | 2 (4) | 5 (1) | 3 (3) |
| CFBP2286 | 1 | 4 (2) | 4 (2) | 4 (2) | 5 (1) |
| CFBP7331 | 1 | 4 (1) | 2 (4) | 3 (2) | 3 (2) |
| CFBP7341 | 1 | 4 (1) | 2 (3) | 3 (2) | 2 (3) |
| CFBP7342 | 1 | 5 (2) | 2 (4) | 4 (3) | 6 (1) |
| L8 | 1 | 9 (2) | 8 (3) | 6 (4) | 12 (1) |
| RS105 | 1 | 3 (2) | 3 (2) | 3 (2) | 4 (1) |
| avg. rank | 1 | 1.8 | 3.1 | 2.4 | 1.5 |
| B8-12 | 10 | 11 (4) | 13 (2) | 17 (1) | 13 (2) |
| BLS256 | 10 | 10 (4) | 11 (3) | 17 (1) | 15 (2) |
| BLS279 | 10 | 8 (4) | 10 (3) | 12 (1) | 12 (1) |
| BXOR1 | 10 | 13 (3) | 11 (4) | 15 (1) | 14 (2) |
| CFBP2286 | 10 | 9 (4) | 10 (2) | 12 (1) | 10 (2) |
| CFBP7331 | 10 | 7 (4) | 9 (1) | 9 (1) | 9 (1) |
| CFBP7341 | 10 | 7 (3) | 9 (1) | 7 (3) | 9 (1) |
| CFBP7342 | 10 | 10 (1) | 10 (1) | 9 (4) | 10 (1) |
| L8 | 10 | 14 (4) | 18 (2) | 18 (2) | 20 (1) |
| RS105 | 10 | 8 (4) | 10 (2) | 12 (1) | 10 (2) |
| avg. rank | 10 | 3.5 | 2.1 | 1.6 | 1.5 |
| B8-12 | 20 | 15 (4) | 19 (1) | 18 (3) | 19 (1) |
| BLS256 | 20 | 15 (4) | 18 (2) | 17 (3) | 19 (1) |
| BLS279 | 20 | 12 (4) | 15 (2) | 14 (3) | 19 (1) |
| BXOR1 | 20 | 14 (3) | 14 (3) | 17 (2) | 18 (1) |
| CFBP2286 | 20 | 10 (4) | 13 (1) | 12 (3) | 13 (1) |
| CFBP7331 | 20 | 9 (4) | 10 (2) | 10 (2) | 11 (1) |
| CFBP7341 | 20 | 8 (3) | 9 (2) | 8 (3) | 10 (1) |
| CFBP7342 | 20 | 11 (1) | 10 (3) | 11 (1) | 10 (3) |
| L8 | 20 | 18 (4) | 24 (1) | 20 (3) | 24 (1) |
| RS105 | 20 | 11 (4) | 15 (1) | 13 (3) | 15 (1) |
| avg. rank | 20 | 3.5 | 1.8 | 2.6 | 1.2 |
| B8-12 | 50 | 24 (1) | 22 (3) | 20 (4) | 23 (2) |
| BLS256 | 50 | 21 (2) | 20 (4) | 21 (2) | 24 (1) |
| BLS279 | 50 | 21 (2) | 18 (3) | 18 (3) | 24 (1) |
| BXOR1 | 50 | 19 (4) | 21 (1) | 20 (2) | 20 (2) |
| CFBP2286 | 50 | 13 (4) | 14 (3) | 16 (1) | 16 (1) |
| CFBP7331 | 50 | 13 (2) | 13 (2) | 11 (4) | 14 (1) |
| CFBP7341 | 50 | 11 (3) | 13 (2) | 9 (4) | 14 (1) |
| CFBP7342 | 50 | 13 (2) | 13 (2) | 14 (1) | 13 (2) |
| L8 | 50 | 25 (3) | 26 (2) | 24 (4) | 28 (1) |
| RS105 | 50 | 18 (1) | 16 (4) | 17 (3) | 18 (1) |
| avg. rank | 50 | 2.4 | 2.6 | 2.8 | 1.3 |
| B8-12 | TALEs AUC | 862 (4) | 878 (2) | 864 (3) | 923 (1) |
| BLS256 | TALEs AUC | 772 (4) | 825 (3) | 878 (2) | 965 (1) |
| BLS279 | TALEs AUC | 692 (4) | 704 (3) | 732 (2) | 937 (1) |
| BXOR1 | TALEs AUC | 759 (3) | 749 (4) | 830 (2) | 840 (1) |
| CFBP2286 | TALEs AUC | 520 (4) | 598 (3) | 612 (2) | 631 (1) |
| CFBP7331 | TALEs AUC | 461 (4) | 502 (2) | 486 (3) | 561 (1) |
| CFBP7341 | TALEs AUC | 427 (3) | 498 (2) | 386 (4) | 544 (1) |
| CFBP7342 | TALEs AUC | 545 (1) | 522 (4) | 534 (3) | 543 (2) |
| L8 | TALEs AUC | 988 (4) | 1115 (2) | 1006 (3) | 1188 (1) |
| RS105 | TALEs AUC | 603 (4) | 665 (2) | 642 (3) | 717 (1) |
| avg. rank | TALEs AUC | 3.5 | 2.7 | 2.7 | 1.1 |

**Supplementary Table L.** Performance evaluation on the level of TALEs for ten *Xoo* strains. For each strain and each approach, we list the number of TALEs with at least one predicted target gene that is also up-regulated in the infection (q-value $< 0.05$, log fold change $> 2$) for different cutoffs on the number of predictions per TALE, i.e., prediction ranks. For each threshold, we additionally report the average rank of each tool.

| measure | Target Finder | TALgetter | Talvez | PrediTALE | Quade | TALgetter vs TALESF | Talvez vs Target Finder | Talvez vs TALgetter | PrediTALE vs Target Finder | PrediTALE vs TALgetter | PrediTALE vs Talvez |
|---|---|---|---|---|---|---|---|---|---|---|---|
| TALEs R1 | 1.8 | 3.1 | 2.4 | 1.5 | ** | — | - | | | +++ | ++ |
| TALEs R10 | 3.5 | 2.1 | 1.6 | 1.5 | *** | + | +++ | ++ | +++ | | |
| TALEs R20 | 3.5 | 1.8 | 2.6 | 1.2 | *** | +++ | + | - | +++ | | +++ |
| TALEs R50 | 2.4 | 2.6 | 2.8 | 1.3 | ** | | | | ++ | +++ | +++ |
| TALEs AUC | 3.5 | 2.7 | 2.7 | 1.1 | *** | ++ | ++ | | +++ | +++ | +++ |
| Genes R1 | 1.8 | 3.1 | 2.4 | 1.5 | ** | — | - | | | +++ | ++ |
| Genes R10 | 3.6 | 2.6 | 1.6 | 1.6 | *** | + | +++ | + | +++ | + | |
| Genes R20 | 3.4 | 2.2 | 2.9 | 1.3 | *** | +++ | + | | +++ | ++ | +++ |
| Genes R50 | 2.3 | 3.1 | 2.9 | 1.1 | ** | | | | +++ | +++ | +++ |
| Genes AUC | 3.3 | 2.9 | 2.7 | 1.1 | *** | | + | | +++ | +++ | +++ |

**Supplementary Table M.** Testing the significance of differences in prediction performance (q-value $< 0.05$, log fold change $> 2$). For each tool and each measure (TALEs/Genes; rank cutoff), we report the average rank per tool, the significance of the Quade test (*:$< 0.05$; **:$< 0.01$; ***:$< 0.001$), and the significance of the pairwise comparison in a post-hoc test. Here, '+' and '-' indicate that the first tool has gained a significantly better or worse performance than the second one, respectively. The number of symbols encodes the significance level in analogy to the Quade test.

| strain | # predictions | Target.Finder | TALgetter | Talvez | PrediTALE |
|---|---|---|---|---|---|
| B8-12 | 1 | 8 (2) | 8 (2) | 5 (4) | 11 (1) |
| BLS256 | 1 | 7 (3) | 8 (2) | 5 (4) | 10 (1) |
| BLS279 | 1 | 6 (3) | 7 (2) | 5 (4) | 8 (1) |
| BXOR1 | 1 | 5 (2) | 5 (2) | 6 (1) | 4 (4) |
| CFBP2286 | 1 | 5 (3) | 6 (2) | 4 (4) | 7 (1) |
| CFBP7331 | 1 | 6 (2) | 7 (1) | 5 (3) | 5 (3) |
| CFBP7341 | 1 | 6 (1) | 2 (3) | 3 (2) | 2 (3) |
| CFBP7342 | 1 | 5 (4) | 7 (1) | 6 (3) | 7 (1) |
| L8 | 1 | 11 (2) | 9 (3) | 6 (4) | 14 (1) |
| RS105 | 1 | 5 (3) | 7 (2) | 4 (4) | 9 (1) |
| avg. rank | 1 | 2.5 | 2.0 | 3.3 | 1.7 |
| B8-12 | 10 | 22 (4) | 26 (3) | 33 (1) | 30 (2) |
| BLS256 | 10 | 18 (4) | 26 (3) | 29 (1) | 28 (2) |
| BLS279 | 10 | 18 (4) | 27 (3) | 28 (2) | 31 (1) |
| BXOR1 | 10 | 23 (4) | 28 (1) | 28 (1) | 28 (1) |
| CFBP2286 | 10 | 17 (3) | 16 (4) | 18 (2) | 19 (1) |
| CFBP7331 | 10 | 14 (4) | 21 (3) | 23 (2) | 24 (1) |
| CFBP7341 | 10 | 14 (2) | 12 (4) | 14 (2) | 17 (1) |
| CFBP7342 | 10 | 14 (4) | 18 (2) | 18 (2) | 22 (1) |
| L8 | 10 | 28 (4) | 38 (2) | 36 (3) | 40 (1) |
| RS105 | 10 | 16 (4) | 20 (3) | 23 (2) | 26 (1) |
| avg. rank | 10 | 3.7 | 2.8 | 1.8 | 1.2 |
| B8-12 | 20 | 36 (4) | 42 (2) | 40 (3) | 48 (1) |
| BLS256 | 20 | 31 (4) | 40 (2) | 36 (3) | 46 (1) |
| BLS279 | 20 | 32 (4) | 41 (2) | 38 (3) | 45 (1) |
| BXOR1 | 20 | 36 (3) | 35 (4) | 38 (1) | 37 (2) |
| CFBP2286 | 20 | 25 (3) | 26 (2) | 22 (4) | 29 (1) |
| CFBP7331 | 20 | 27 (4) | 30 (1) | 30 (1) | 30 (1) |
| CFBP7341 | 20 | 20 (2) | 17 (3) | 16 (4) | 21 (1) |
| CFBP7342 | 20 | 24 (2) | 24 (2) | 23 (4) | 28 (1) |
| L8 | 20 | 43 (4) | 57 (2) | 47 (3) | 58 (1) |
| RS105 | 20 | 28 (3) | 31 (2) | 28 (3) | 35 (1) |
| avg. rank | 20 | 3.3 | 2.2 | 2.9 | 1.1 |
| B8-12 | 50 | 75 (4) | 78 (2) | 78 (2) | 84 (1) |
| BLS256 | 50 | 72 (4) | 78 (3) | 79 (2) | 96 (1) |
| BLS279 | 50 | 67 (4) | 71 (3) | 72 (2) | 85 (1) |
| BXOR1 | 50 | 59 (3) | 56 (4) | 64 (1) | 63 (2) |
| CFBP2286 | 50 | 38 (2) | 33 (4) | 36 (3) | 39 (1) |
| CFBP7331 | 50 | 50 (2) | 46 (3) | 45 (4) | 51 (1) |
| CFBP7341 | 50 | 37 (1) | 33 (3) | 30 (4) | 37 (1) |
| CFBP7342 | 50 | 47 (3) | 48 (2) | 42 (4) | 52 (1) |
| L8 | 50 | 85 (4) | 102 (2) | 88 (3) | 103 (1) |
| RS105 | 50 | 47 (4) | 49 (3) | 51 (2) | 57 (1) |
| avg. rank | 50 | 3.1 | 2.9 | 2.7 | 1.1 |
| B8-12 | Genes AUC | 2123 (4) | 2382 (2) | 2293 (3) | 2647 (1) |
| BLS256 | Genes AUC | 1865 (4) | 2260 (2) | 2211 (3) | 2675 (1) |
| BLS279 | Genes AUC | 1829 (4) | 2207 (2) | 2176 (3) | 2614 (1) |
| BXOR1 | Genes AUC | 1963 (3) | 1909 (4) | 2102 (2) | 2125 (1) |
| CFBP2286 | Genes AUC | 1221 (4) | 1240 (2) | 1235 (3) | 1453 (1) |
| CFBP7331 | Genes AUC | 1507 (4) | 1537 (2) | 1528 (3) | 1708 (1) |
| CFBP7341 | Genes AUC | 1194 (2) | 977 (3) | 928 (4) | 1217 (1) |
| CFBP7342 | Genes AUC | 1385 (3) | 1438 (2) | 1324 (4) | 1665 (1) |
| L8 | Genes AUC | 2525 (4) | 3182 (2) | 2762 (3) | 3339 (1) |
| RS105 | Genes AUC | 1398 (4) | 1637 (2) | 1571 (3) | 1901 (1) |
| avg. rank | Genes AUC | 3.6 | 2.3 | 3.1 | 1.0 |

**Supplementary Table N.** Performance evaluation on the level of target genes for ten *Xoc* strains. For each strain and each approach, we list the number of predicted target genes that are also up-regulated in the infection (q-value $< 0.01$, log fold change $> 1$) for different cutoffs on the number of predictions per TALE, i.e., prediction ranks. For each threshold, we additionally report the average rank of each tool.

| strain | # predictions | Target.Finder | TALgetter | Talvez | PrediTALE |
|---|---|---|---|---|---|
| B8-12 | 1 | 8 (2) | 8 (2) | 5 (4) | 11 (1) |
| BLS256 | 1 | 7 (3) | 8 (2) | 5 (4) | 10 (1) |
| BLS279 | 1 | 6 (3) | 7 (2) | 5 (4) | 8 (1) |
| BXOR1 | 1 | 5 (2) | 5 (2) | 6 (1) | 4 (4) |
| CFBP2286 | 1 | 5 (3) | 6 (2) | 4 (4) | 7 (1) |
| CFBP7331 | 1 | 6 (2) | 7 (1) | 5 (3) | 5 (3) |
| CFBP7341 | 1 | 6 (1) | 2 (3) | 3 (2) | 2 (3) |
| CFBP7342 | 1 | 5 (4) | 7 (1) | 6 (3) | 7 (1) |
| L8 | 1 | 11 (2) | 9 (3) | 6 (4) | 14 (1) |
| RS105 | 1 | 5 (3) | 7 (2) | 4 (4) | 9 (1) |
| avg. rank | 1 | 2.5 | 2.0 | 3.3 | 1.7 |
| B8-12 | 10 | 15 (4) | 16 (3) | 21 (1) | 20 (2) |
| BLS256 | 10 | 13 (4) | 16 (3) | 19 (1) | 19 (1) |
| BLS279 | 10 | 13 (4) | 16 (3) | 18 (2) | 19 (1) |
| BXOR1 | 10 | 15 (4) | 17 (3) | 18 (1) | 18 (1) |
| CFBP2286 | 10 | 13 (3) | 13 (3) | 15 (1) | 15 (1) |
| CFBP7331 | 10 | 10 (4) | 13 (2) | 12 (3) | 14 (1) |
| CFBP7341 | 10 | 9 (2) | 9 (2) | 9 (2) | 10 (1) |
| CFBP7342 | 10 | 11 (3) | 13 (2) | 11 (3) | 15 (1) |
| L8 | 10 | 17 (4) | 23 (1) | 20 (3) | 22 (2) |
| RS105 | 10 | 12 (4) | 13 (3) | 17 (1) | 17 (1) |
| avg. rank | 10 | 3.6 | 2.5 | 1.8 | 1.2 |
| B8-12 | 20 | 20 (4) | 25 (1) | 23 (2) | 23 (2) |
| BLS256 | 20 | 18 (4) | 22 (1) | 20 (3) | 22 (1) |
| BLS279 | 20 | 18 (4) | 23 (1) | 20 (3) | 22 (2) |
| BXOR1 | 20 | 22 (1) | 18 (4) | 22 (1) | 21 (3) |
| CFBP2286 | 20 | 15 (4) | 17 (2) | 17 (2) | 18 (1) |
| CFBP7331 | 20 | 15 (2) | 15 (2) | 16 (1) | 15 (2) |
| CFBP7341 | 20 | 10 (2) | 10 (2) | 10 (2) | 11 (1) |
| CFBP7342 | 20 | 14 (2) | 13 (3) | 12 (4) | 16 (1) |
| L8 | 20 | 21 (4) | 28 (1) | 23 (3) | 26 (2) |
| RS105 | 20 | 17 (4) | 20 (1) | 18 (3) | 20 (1) |
| avg. rank | 20 | 3.1 | 1.8 | 2.4 | 1.6 |
| B8-12 | 50 | 26 (4) | 27 (2) | 27 (2) | 28 (1) |
| BLS256 | 50 | 26 (2) | 26 (2) | 26 (2) | 27 (1) |
| BLS279 | 50 | 24 (3) | 25 (2) | 24 (3) | 26 (1) |
| BXOR1 | 50 | 23 (2) | 21 (4) | 24 (1) | 23 (2) |
| CFBP2286 | 50 | 19 (3) | 17 (4) | 20 (2) | 21 (1) |
| CFBP7331 | 50 | 17 (3) | 18 (2) | 17 (3) | 20 (1) |
| CFBP7341 | 50 | 13 (3) | 15 (2) | 13 (3) | 18 (1) |
| CFBP7342 | 50 | 19 (3) | 20 (1) | 19 (3) | 20 (1) |
| L8 | 50 | 28 (3) | 29 (1) | 27 (4) | 29 (1) |
| RS105 | 50 | 19 (4) | 21 (2) | 21 (2) | 23 (1) |
| avg. rank | 50 | 3.0 | 2.2 | 2.5 | 1.1 |
| B8-12 | TALEs AUC | 1031 (4) | 1145 (2) | 1118 (3) | 1152 (1) |
| BLS256 | TALEs AUC | 971 (4) | 1064 (2) | 1022 (3) | 1091 (1) |
| BLS279 | TALEs AUC | 954 (4) | 1058 (2) | 1008 (3) | 1109 (1) |
| BXOR1 | TALEs AUC | 973 (3) | 909 (4) | 1023 (1) | 1005 (2) |
| CFBP2286 | TALEs AUC | 735 (4) | 766 (3) | 819 (2) | 899 (1) |
| CFBP7331 | TALEs AUC | 677 (4) | 753 (2) | 723 (3) | 799 (1) |
| CFBP7341 | TALEs AUC | 498 (4) | 559 (2) | 500 (3) | 613 (1) |
| CFBP7342 | TALEs AUC | 693 (3) | 764 (2) | 679 (4) | 863 (1) |
| L8 | TALEs AUC | 1146 (4) | 1302 (1) | 1166 (3) | 1269 (2) |
| RS105 | TALEs AUC | 795 (4) | 896 (2) | 870 (3) | 974 (1) |
| avg. rank | TALEs AUC | 3.8 | 2.2 | 2.8 | 1.2 |

**Supplementary Table O.** Performance evaluation on the level of TALEs for ten *Xoo* strains. For each strain and each approach, we list the number of TALEs with at least one predicted target gene that is also up-regulated in the infection (q-value $< 0.01$, log fold change $> 1$) for different cutoffs on the number of predictions per TALE, i.e., prediction ranks. For each threshold, we additionally report the average rank of each tool.

| measure | Target Finder | TALgetter | Talvez | PrediTALE | Quade | TALgetter vs TALESF | Talvez vs Target Finder | Talvez vs TALgetter | PrediTALE vs Target Finder | PrediTALE vs TALgetter | PrediTALE vs Talvez |
|---|---|---|---|---|---|---|---|---|---|---|---|
| TALEs R1 | 2.5 | 2 | 3.3 | 1.7 | ** | | - | − | + | + | +++ |
| TALEs R10 | 3.6 | 2.5 | 1.8 | 1.2 | *** | ++ | +++ | | +++ | + | |
| TALEs R20 | 3.1 | 1.8 | 2.4 | 1.6 | * | +++ | | - | +++ | | |
| TALEs R50 | 3 | 2.2 | 2.5 | 1.1 | ** | | | | +++ | ++ | +++ |
| TALEs AUC | 3.8 | 2.2 | 2.8 | 1.2 | *** | +++ | + | - | +++ | + | +++ |
| Genes R1 | 2.5 | 2 | 3.3 | 1.7 | ** | | - | − | + | + | +++ |
| Genes R10 | 3.7 | 2.8 | 1.8 | 1.2 | *** | + | +++ | + | +++ | +++ | |
| Genes R20 | 3.3 | 2.2 | 2.9 | 1.1 | *** | +++ | | - | +++ | + | +++ |
| Genes R50 | 3.1 | 2.9 | 2.7 | 1.1 | *** | | + | | +++ | +++ | +++ |
| Genes AUC | 3.6 | 2.3 | 3.1 | 1 | *** | +++ | | - | +++ | ++ | +++ |

**Supplementary Table P.** Testing the significance of differences in prediction performance (q-value $< 0.01$, log fold change $> 1$). For each tool and each measure (TALEs/Genes; rank cutoff), we report the average rank per tool, the significance of the Quade test (*:$< 0.05$; **:$< 0.01$; ***:$< 0.001$), and the significance of the pairwise comparison in a post-hoc test. Here, '+' and '-' indicate that the first tool has gained a significantly better or worse performance than the second one, respectively. The number of symbols encodes the significance level in analogy to the Quade test.

| strain | # predictions | Target.Finder | TALgetter | Talvez | PrediTALE |
|---|---|---|---|---|---|
| B8-12 | 1 | 6 (2) | 6 (2) | 4 (4) | 7 (1) |
| BLS256 | 1 | 5 (4) | 6 (2) | 6 (2) | 7 (1) |
| BLS279 | 1 | 4 (2) | 4 (2) | 4 (2) | 5 (1) |
| BXOR1 | 1 | 4 (2) | 3 (3) | 5 (1) | 3 (3) |
| CFBP2286 | 1 | 4 (3) | 5 (1) | 4 (3) | 5 (1) |
| CFBP7331 | 1 | 4 (1) | 3 (2) | 3 (2) | 3 (2) |
| CFBP7341 | 1 | 4 (1) | 2 (3) | 3 (2) | 2 (3) |
| CFBP7342 | 1 | 6 (1) | 3 (4) | 4 (3) | 6 (1) |
| L8 | 1 | 9 (2) | 9 (2) | 7 (4) | 11 (1) |
| RS105 | 1 | 3 (3) | 4 (1) | 3 (3) | 4 (1) |
| avg. rank | 1 | 2.1 | 2.2 | 2.6 | 1.5 |
| B8-12 | 10 | 20 (4) | 23 (1) | 22 (3) | 23 (1) |
| BLS256 | 10 | 18 (4) | 22 (3) | 24 (2) | 25 (1) |
| BLS279 | 10 | 15 (4) | 19 (2) | 17 (3) | 21 (1) |
| BXOR1 | 10 | 18 (3) | 17 (4) | 22 (2) | 23 (1) |
| CFBP2286 | 10 | 12 (4) | 13 (2) | 13 (2) | 14 (1) |
| CFBP7331 | 10 | 11 (4) | 13 (2) | 13 (2) | 16 (1) |
| CFBP7341 | 10 | 10 (4) | 12 (2) | 11 (3) | 15 (1) |
| CFBP7342 | 10 | 12 (1) | 12 (1) | 10 (4) | 12 (1) |
| L8 | 10 | 22 (4) | 29 (2) | 27 (3) | 31 (1) |
| RS105 | 10 | 14 (3) | 17 (2) | 14 (3) | 18 (1) |
| avg. rank | 10 | 3.5 | 2.1 | 2.7 | 1.0 |
| B8-12 | 20 | 29 (4) | 30 (2) | 30 (2) | 32 (1) |
| BLS256 | 20 | 26 (4) | 30 (2) | 33 (1) | 30 (2) |
| BLS279 | 20 | 24 (3) | 24 (3) | 26 (2) | 31 (1) |
| BXOR1 | 20 | 26 (2) | 22 (4) | 26 (2) | 29 (1) |
| CFBP2286 | 20 | 16 (3) | 16 (3) | 19 (2) | 20 (1) |
| CFBP7331 | 20 | 18 (2) | 15 (4) | 16 (3) | 22 (1) |
| CFBP7341 | 20 | 18 (2) | 14 (3) | 14 (3) | 19 (1) |
| CFBP7342 | 20 | 22 (1) | 18 (2) | 15 (4) | 16 (3) |
| L8 | 20 | 35 (4) | 40 (2) | 37 (3) | 44 (1) |
| RS105 | 20 | 20 (4) | 21 (2) | 21 (2) | 23 (1) |
| avg. rank | 20 | 2.9 | 2.7 | 2.4 | 1.3 |
| B8-12 | 50 | 43 (4) | 45 (3) | 47 (1) | 47 (1) |
| BLS256 | 50 | 42 (4) | 44 (3) | 53 (2) | 56 (1) |
| BLS279 | 50 | 38 (3) | 37 (4) | 40 (2) | 47 (1) |
| BXOR1 | 50 | 35 (4) | 36 (3) | 41 (2) | 42 (1) |
| CFBP2286 | 50 | 22 (4) | 23 (3) | 25 (2) | 26 (1) |
| CFBP7331 | 50 | 26 (3) | 27 (2) | 25 (4) | 28 (1) |
| CFBP7341 | 50 | 25 (1) | 24 (2) | 21 (4) | 24 (2) |
| CFBP7342 | 50 | 28 (3) | 32 (1) | 28 (3) | 29 (2) |
| L8 | 50 | 50 (4) | 54 (3) | 58 (2) | 63 (1) |
| RS105 | 50 | 28 (3) | 27 (4) | 35 (1) | 29 (2) |
| avg. rank | 50 | 3.3 | 2.8 | 2.3 | 1.3 |
| B8-12 | Genes AUC | 1469 (4) | 1626 (2) | 1624 (3) | 1685 (1) |
| BLS256 | Genes AUC | 1388 (4) | 1544 (3) | 1772 (2) | 1799 (1) |
| BLS279 | Genes AUC | 1239 (4) | 1342 (3) | 1353 (2) | 1675 (1) |
| BXOR1 | Genes AUC | 1249 (3) | 1233 (4) | 1417 (2) | 1436 (1) |
| CFBP2286 | Genes AUC | 817 (4) | 879 (3) | 950 (2) | 1008 (1) |
| CFBP7331 | Genes AUC | 871 (3) | 891 (2) | 846 (4) | 1059 (1) |
| CFBP7341 | Genes AUC | 857 (2) | 811 (3) | 736 (4) | 924 (1) |
| CFBP7342 | Genes AUC | 1008 (1) | 970 (2) | 859 (4) | 913 (3) |
| L8 | Genes AUC | 1756 (4) | 2033 (2) | 2005 (3) | 2255 (1) |
| RS105 | Genes AUC | 999 (4) | 1083 (3) | 1177 (1) | 1140 (2) |
| avg. rank | Genes AUC | 3.3 | 2.7 | 2.7 | 1.3 |

**Supplementary Table Q.** Performance evaluation on the level of target genes for ten *Xoc* strains when filtering for predictions of TALE boxes on the same strand as the downstream gene. For each strain and each approach, we list the number of predicted target genes that are also up-regulated in the infection for different cutoffs on the number of predictions per TALE, i.e., prediction ranks. For each threshold, we additionally report the average rank of each tool.

| strain | # predictions | Target.Finder | TALgetter | Talvez | PrediTALE |
|---|---|---|---|---|---|
| B8-12 | 1 | 6 (2) | 6 (2) | 4 (4) | 7 (1) |
| BLS256 | 1 | 5 (4) | 6 (2) | 6 (2) | 7 (1) |
| BLS279 | 1 | 4 (2) | 4 (2) | 4 (2) | 5 (1) |
| BXOR1 | 1 | 4 (2) | 3 (3) | 5 (1) | 3 (3) |
| CFBP2286 | 1 | 4 (3) | 5 (1) | 4 (3) | 5 (1) |
| CFBP7331 | 1 | 4 (1) | 3 (2) | 3 (2) | 3 (2) |
| CFBP7341 | 1 | 4 (1) | 2 (3) | 3 (2) | 2 (3) |
| CFBP7342 | 1 | 6 (1) | 3 (4) | 4 (3) | 6 (1) |
| L8 | 1 | 9 (2) | 9 (2) | 7 (4) | 11 (1) |
| RS105 | 1 | 3 (3) | 4 (1) | 3 (3) | 4 (1) |
| avg. rank | 1 | 2.1 | 2.2 | 2.6 | 1.5 |
| B8-12 | 10 | 14 (4) | 16 (1) | 16 (1) | 16 (1) |
| BLS256 | 10 | 13 (4) | 14 (3) | 17 (1) | 17 (1) |
| BLS279 | 10 | 11 (4) | 13 (2) | 12 (3) | 15 (1) |
| BXOR1 | 10 | 13 (3) | 13 (3) | 16 (1) | 16 (1) |
| CFBP2286 | 10 | 10 (4) | 11 (2) | 12 (1) | 11 (2) |
| CFBP7331 | 10 | 7 (4) | 9 (1) | 8 (3) | 9 (1) |
| CFBP7341 | 10 | 7 (3) | 9 (1) | 7 (3) | 9 (1) |
| CFBP7342 | 10 | 10 (1) | 9 (2) | 9 (2) | 9 (2) |
| L8 | 10 | 17 (4) | 21 (1) | 20 (3) | 21 (1) |
| RS105 | 10 | 10 (4) | 12 (2) | 11 (3) | 13 (1) |
| avg. rank | 10 | 3.5 | 1.8 | 2.1 | 1.2 |
| B8-12 | 20 | 19 (1) | 18 (4) | 19 (1) | 19 (1) |
| BLS256 | 20 | 17 (4) | 18 (3) | 19 (2) | 20 (1) |
| BLS279 | 20 | 16 (3) | 15 (4) | 17 (2) | 20 (1) |
| BXOR1 | 20 | 17 (3) | 16 (4) | 18 (2) | 19 (1) |
| CFBP2286 | 20 | 12 (3) | 12 (3) | 13 (1) | 13 (1) |
| CFBP7331 | 20 | 9 (3) | 10 (2) | 9 (3) | 12 (1) |
| CFBP7341 | 20 | 10 (2) | 10 (2) | 8 (4) | 12 (1) |
| CFBP7342 | 20 | 13 (1) | 11 (3) | 12 (2) | 11 (3) |
| L8 | 20 | 21 (3) | 24 (2) | 21 (3) | 25 (1) |
| RS105 | 20 | 14 (2) | 14 (2) | 14 (2) | 15 (1) |
| avg. rank | 20 | 2.5 | 2.9 | 2.2 | 1.2 |
| B8-12 | 50 | 23 (2) | 25 (1) | 22 (3) | 22 (3) |
| BLS256 | 50 | 19 (4) | 24 (1) | 21 (3) | 22 (2) |
| BLS279 | 50 | 20 (2) | 20 (2) | 19 (4) | 22 (1) |
| BXOR1 | 50 | 20 (4) | 21 (3) | 22 (2) | 23 (1) |
| CFBP2286 | 50 | 13 (4) | 17 (1) | 16 (2) | 16 (2) |
| CFBP7331 | 50 | 12 (3) | 12 (3) | 13 (1) | 13 (1) |
| CFBP7341 | 50 | 12 (2) | 12 (2) | 11 (4) | 13 (1) |
| CFBP7342 | 50 | 14 (2) | 14 (2) | 15 (1) | 13 (4) |
| L8 | 50 | 24 (4) | 27 (1) | 27 (1) | 27 (1) |
| RS105 | 50 | 17 (4) | 18 (1) | 18 (1) | 18 (1) |
| avg. rank | 50 | 3.1 | 1.7 | 2.2 | 1.7 |
| B8-12 | TALEs AUC | 901 (4) | 971 (1) | 912 (3) | 924 (2) |
| BLS256 | TALEs AUC | 797 (4) | 924 (2) | 916 (3) | 981 (1) |
| BLS279 | TALEs AUC | 757 (4) | 800 (2) | 775 (3) | 949 (1) |
| BXOR1 | TALEs AUC | 776 (4) | 818 (3) | 888 (1) | 887 (2) |
| CFBP2286 | TALEs AUC | 552 (4) | 657 (3) | 664 (1) | 662 (2) |
| CFBP7331 | TALEs AUC | 432 (4) | 517 (2) | 469 (3) | 562 (1) |
| CFBP7341 | TALEs AUC | 458 (3) | 514 (2) | 421 (4) | 557 (1) |
| CFBP7342 | TALEs AUC | 589 (1) | 535 (4) | 553 (2) | 540 (3) |
| L8 | TALEs AUC | 994 (4) | 1163 (2) | 1099 (3) | 1186 (1) |
| RS105 | TALEs AUC | 638 (4) | 732 (2) | 704 (3) | 752 (1) |
| avg. rank | TALEs AUC | 3.6 | 2.3 | 2.6 | 1.5 |

**Supplementary Table R.** Performance evaluation on the level of TALEs for ten *Xoo* strains when filtering for predictions of TALE boxes on the same strand as the downstream gene. For each strain and each approach, we list the number of TALEs with at least one predicted target gene that is also up-regulated in the infection for different cutoffs on the number of predictions per TALE, i.e., prediction ranks. For each threshold, we additionally report the average rank of each tool.

| measure | Target Finder | TALgetter | Talvez | PrediTALE | Quade | TALgetter vs TALESF | Talvez vs Target Finder | Talvez vs TALgetter | PrediTALE vs Target Finder | PrediTALE vs TALgetter | PrediTALE vs Talvez |
|---|---|---|---|---|---|---|---|---|---|---|---|
| Genes R1 | 2.1 | 2.2 | 2.6 | 1.5 | | | | | | | |
| Genes R10 | 3.5 | 2.1 | 2.7 | 1 | *** | ++ | ++ | | +++ | +++ | +++ |
| Genes R20 | 2.9 | 2.7 | 2.4 | 1.3 | * | | | | +++ | +++ | ++ |
| Genes R50 | 3.3 | 2.8 | 2.3 | 1.3 | ** | | ++ | + | +++ | +++ | |
| Genes AUC | 3.3 | 2.7 | 2.7 | 1.3 | *** | + | + | | +++ | +++ | +++ |
| TALEs R1 | 2.1 | 2.2 | 2.6 | 1.5 | | | | | | | |
| TALEs R10 | 3.5 | 1.8 | 2.1 | 1.2 | *** | +++ | +++ | | +++ | | + |
| TALEs R20 | 2.5 | 2.9 | 2.2 | 1.2 | ** | | | | +++ | +++ | +++ |
| TALEs R50 | 3.1 | 1.7 | 2.2 | 1.7 | * | +++ | | | ++ | | |
| TALEs AUC | 3.6 | 2.3 | 2.6 | 1.5 | *** | +++ | + | | +++ | + | +++ |

**Supplementary Table S.** Testing the significance of differences in prediction performance when filtering for predictions of TALE boxes on the same strand as the downstream gene. For each tool and each measure (TALEs/Genes; rank cutoff), we report the average rank per tool, the significance of the Quade test (*:$<$ 0.05; **:$<$ 0.01; ***:$<$ 0.001), and the significance of the pairwise comparison in a post-hoc test. Here, '+' and '-' indicate that the first tool has gained a significantly better or worse performance than the second one, respectively. The number of symbols encodes the significance level in analogy to the Quade test.

**Supplementary Table T.** Gene abundances and sleuth output for *Xoo* strains. [Available as separate XLS file `RNA-seq-Xoo.xls`]

**Supplementary Table U.** Gene abundances and sleuth output for *Xoc* strains. [Available as separate XLS file `RNA-seq-Xoc.xls`]

**Supplementary Table V.** Complete list of top 20 predictions for all approaches and *Xoo* and *Xoc* strains.
[Available as separate XLS file `predictions.xls`]

**Supplementary Table W.** Results of genome-wide predictions for *Xoo* strains. [Available as separate XLS file `GenomeWidePredictions_Xoo.xls`]

**Supplementary Table X.** Results of genome-wide predictions for *Xoc* strains. [Available as separate XLS file `GenomeWidePredictions_Xoc.xls`]
